# TRPM8 protein dynamics correlates with ligand structure and cellular function

**DOI:** 10.1101/2025.05.13.653789

**Authors:** Mubark D. Mebrat, Dustin D. Luu, Jacob K. Hilton, Minjoo Kim, Kaitlyn Parrott, Brian R. Cherry, Marcia Levitus, V. Blair Journigan, Wade D. Van Horn

**Affiliations:** School of Molecular Sciences, Arizona State University, Tempe, AZ 85281, United States; The Biodesign Institute Center for Personalized Diagnostics, Arizona State University, Tempe, AZ 85281, United States; The Biodesign Institute Center for Single Molecule Biophysics, Arizona State University, Tempe, AZ, 85287, United States; The Magnetic Resonance Research Center, Arizona State University, Tempe, AZ, 85287, United States; Department of Pharmaceutical Sciences, Computational Chemical Genomics Screening Center, University of Pittsburgh, Pittsburgh, PA 15260, United States; Department of Chemistry, University of Pittsburgh, Pittsburgh, PA 15260, United States

## Abstract

Protein dynamics have emerged as a key feature associated with function in various systems. Here, NMR-based studies coupled with computational cheminformatics and cellular function are leveraged to identify a relationship between the human cold and menthol receptor TRPM8 dynamics, chemical structure, and cellular potency. TRPM8 is a validated target for a variety of pain indications but generally has been clinically limited by on-target side effects impacting thermosensing and thermoregulation. This study shows that cheminformatic analysis of a TRPM8 regulating small molecule ligand library correlates with cellular function. Electrophysiology studies further validate the relationship and show a correlation between chemical structure and functional features such as compound potency. Solution NMR studies of the TRPM8 voltage sensing-like domain, which houses the canonical menthol ligand binding site, show that ligand binding conformationally selects NMR-detected TRPM8 dynamics in a manner that quantitatively correlates with chemical structure. The relationship between chemical structure and protein dynamics can be used predictively, where a chemical structure is predictive of dynamics in a latent reduced dimensionality space. Moreover, the robustness of the conformational selection of the dynamic ensemble is evaluated by varying related and divergent chemotypes, signal-to-noise sensitivity, and sample bias. Taken together, this study identifies that protein dynamics can serve as a quantifiable bridge between chemical structure and cellular function, which has implications for drug discovery in difficult systems.

## Introduction

Transient receptor potential melastatin member 8 (TRPM8) is a polymodally activated ion channel that functions as the primary human cold sensor and is activated by menthol and other cooling compounds.^1–3^ TRPM8 has been implicated in pain, cancer, obesity, and other pathophysiologies.^4–9^ As a result, TRPM8 has garnered interest as a promising target for a variety of relevant clinical indications.^10–14^ While several TRPM8 small molecules have been developed to selectively and potently inhibit TRPM8 with clinical efficacy, they generally fail in clinical trials.^15,16^ These failures are commonly attributed to on-target adverse effects on thermosensing and thermoregulation that are thought to be related to the polymodal nature of TRPM8 activation by cold and chemical ligands.^16–19^ Identifying mode selective antagonists is a hypothesized imperative for thermal neutral and clinically successful TRPM8 targeting therapeutics.^19^ Alternatively, the thermogenic side effect could be leveraged to induce therapeutic hypothermia in treating traumatic brain injury, cardiac arrest, and acute myocardial infarction.^20,21^

Recently, phenazopyridine (PAP), an orally administered over-the-counter urinary tract analgesic, has been shown to inhibit TRPM8 at concentrations near those found in patient urine.^22^ TRPM8-expressing sensory neurons innervate the bladder and urinary tract, suggesting that TRPM8 inhibition may relate to the PAP mechanism of action. Independent of the role of TRPM8 in PAP-dependent analgesic effects, the widespread use of PAP indicates that TRPM8 inhibition is possible without thermally adverse side effects, further validating TRPM8 for pain and other pathophysiological indications.

Structure-based drug design is a hallmark of modern drug discovery, where protein structures identify the orthosteric binding pocket of the target protein, enabling rational and efficient identification of lead compounds.^23^ In recent years, avian, mouse, and human TRPM8 structures have been determined by cryogenic electron microscopy (cryo-EM) with structural resolution ranging from 2.5 to 4.3 Å.^24–30^ Several of the TRPM8 structures have ligands bound to the orthosteric menthol binding site, which is located in the transmembrane voltage-sensing-like domain (VSLD, transmembrane helices S1-S4).^24–26,28,30^ A few compounds have been shown to bind TRPM8 allosterically on the exterior of the VSLD, in what is the canonical TRPV1–capsaicin binding site.^26,30^ The VSLD has a canonical four-helix bundle structure^31^ with a membrane-embedded cavity in the inner membrane leaflet that serves as a binding pocket for the cognate agonist menthol and other agonists and antagonists.^25,28,29,32^ Mechanistically, menthol binding to the VSLD is coupled to channel opening (gating) of the adjacent pore domain (transmembrane helices S5-S6, Figure 1).^33,34^ The current cryo-EM structures are invaluable for understanding TRPM8 ligand binding and generating testable mechanistic hypotheses. Nevertheless, the ligand-bound TRPM8 structures provide little insight into the discrete VSLD structural changes that happen upon binding, which are ultimately transduced to open and close the pore domain. This suggests that ligands may modulate TRPM8 protein dynamics, a feature that is not clearly captured by existing static structures. In particular, the structural data are consistent with the idea of conformational selection, where ligand binding redistributes the protein conformational ensemble, often with functional outcomes.^35–37^

**Figure 1.**
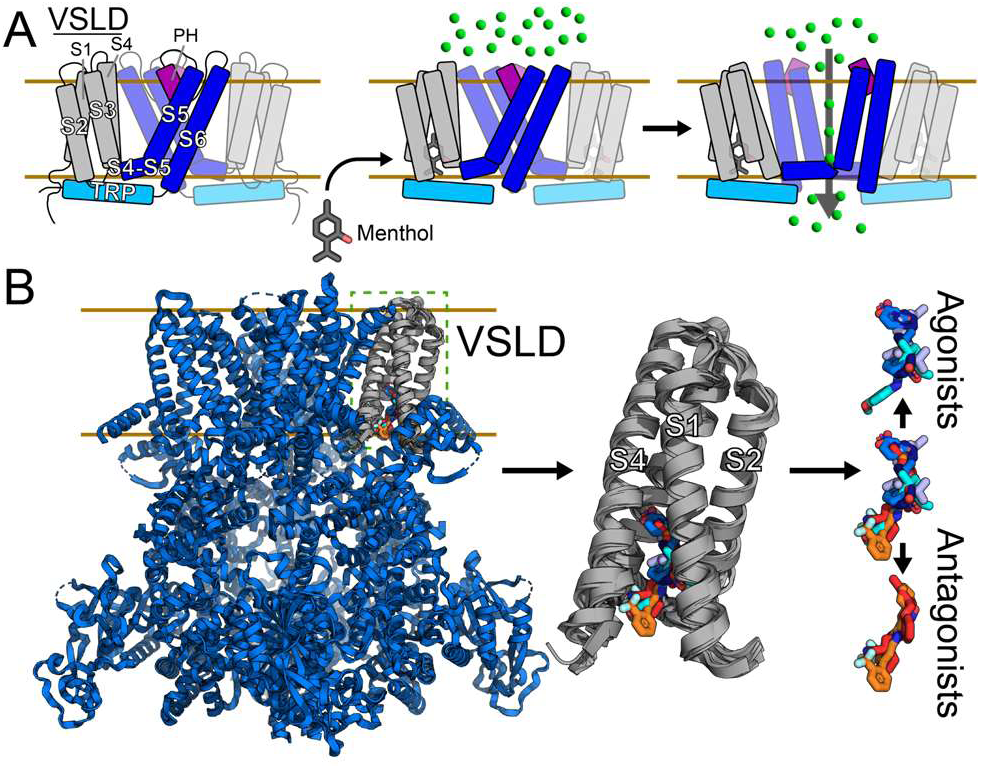
Structural analysis of TRPM8 ligand activation suggests a conformational selection mechanism. (A) Schematic of the TRPM8 transmembrane domain (helices S1-S6) gating mechanism. The orthosteric menthol binding site in the voltage-sensing like domain (VSLD) is transduced to conformational change in the pore domain (S5-S6), whereby initiating signal transduction. (B) Analysis of Cryo-EM structures reveals minimal VSLD structural differences (≤1.21 Å RMSD) between TRPM8 ortholog structures in the presence and absence of activating agonists and inhibiting antagonists. Minimal structural differences in the presence of diverse chemotypes with alternate functional outcomes are consistent with a protein dynamics-based mechanism of conformational selection.

It is increasingly clear that protein dynamics regulate biomolecular function, giving rise to the idea of functional dynamics, where non-random local and allosteric fluctuations over various timescales are crucial to function.^38–40^ Solution nuclear magnetic resonance spectroscopy (NMR) has emerged as a capable technique in quantifying the conformational ensemble and measuring protein dynamics across a wide range of timescales, from picoseconds to weeks.^39–45^ NMR studies are insightful because the data encode diverse protein signatures from the local environment, secondary and tertiary structures, and various dynamic modes. Many varieties of solution NMR techniques provide insight into the conformational ensemble, including ^19^F NMR studies.

Among many isotopically compatible NMR nuclei, ^19^F is particularly well-suited to probe dynamics.^46,47^ ^19^F probes have a particularly broad chemical shift range that covers hundreds of ppm relative to the tens of ppm range for the quintessential NMR ^1^H nucleus.^46^ Accordingly, small spectral differences that arise from conformational changes and protein motions are often readily detected in ^19^F NMR data. Additionally, ^19^F is bioorthogonal, which gives rise to direct and facile signal detection with no contaminating background signal. The benefits of ^19^F studies have been widely used, including to illuminate the functional dynamics of various GPCRs.^48–52^

In this study, we sought to probe TRPM8 functional dynamics. First, we evaluated the general concept of the structure-activity relationship (SAR) for TRPM8. Cheminformatic analysis of a concise and rationally generated library indicates general SAR assumptions hold for TRPM8 despite a central role of TRPM8 dynamics in function. Similarly, unsupervised analysis of a subset of this TRPM8 compound library correlates well with electrophysiology data, validating the relationship between chemical structure and cellular function for this system. Next, we characterized the isolated human TRPM8 VSLD with solution NMR and other biophysical techniques to show that it retains key biological features, including the ability to bind various chemical ligands and undergo temperature-dependent conformational changes. We also established a fluorine-labeled tryptophan hTRPM8-VSLD system that likewise recapitulates TRPM8 chemical binding and cold-sensing features. Lastly, we show that ^19^F NMR-detected TRPM8 dynamics can serve as an intermediate that links chemical structure and cellular function. In particular, signatures of ^19^F dynamic data correlate to both ligand chemical structure and the cellular function as measured from half maximal effective and inhibitory concentrations (*EC*_50_, *IC*_50_). Moreover, TRPM8 dynamics can be leveraged predictively, which has implications for identifying chemical compounds with functional desired features.

## Results and Discussion

### Structural analysis of TRPM8 agonist and antagonist binding

Cryo-EM determined TRPM8 structures have proven pivotal to elucidating ligand binding sites, identifying therapeutically relevant inactive conformational states, and providing unparalleled insight into TRPM8 molecular mechanisms.^53,54^ To date, 29 cryo-EM TRPM8 structures have been determined (Table S1).^24–30^ Among those are 13 that include either an agonist (WS-12, Icilin, cryosim-3) or antagonist (AMTB, TC-I 2014) in the orthosteric VSLD binding site (Figure S1A).^24–26,28^ Evaluation of the ligand-bound TRPM8 structures reveals an interesting feature: the VSLD from bound structures have minimal variance with the average, maximum, and minimum backbone RMSD relative to the unbound human TRPM8 VSLD (pdb 8BDC) of 0.80 Å, 1.16 Å, and 0.51 Å, respectively (Table S2). This is despite the structures being from several species and nominally in distinct activated or inhibited states (Figure 1B). The high VSLD structural homogeneity suggests that large-scale conformations of the VSLD are not required for function and suggests that chemical compounds may modulate TRPM8 protein dynamics, a feature that is not easily captured by static structures. Analysis of all current 29 TRPM8 structures, where an RMSD dissimilarity matrix of the full-length structures was constructed (Table S3) and subjected to a multidimensional scaling (MDS) transform. This analysis supports the idea that relatively small-scale conformational changes and/or alterations of protein dynamics contribute to functional outcomes (Figure S1B), something that has been noted in previous studies.^54–57^

### Cheminformatics links ligand chemical structure and TRPM8 activity

Because orthosteric binding of functionally divergent chemical ligands does not clearly emerge from a meta-analysis of the current TRPM8 structural biology, we sought to assess the linkage between chemical properties of TRPM8 agonists and antagonists with cellular function. Given the increasing breadth of TRPM8-regulating small molecules^16,58,59^ we aimed to investigate the relationship between chemotype and functional activity using cheminformatic and data science tools. Multidimensional scaling (MDS) clustering is an unsupervised dimensionality reduction algorithm that evaluates chemical structural similarity. Using the ChemMine Tools Suite^60^, we selected a representative set of 25 TRPM8-regulating agonists and antagonists (Figure S2), including well-characterized compounds such as menthol, those used in cryo-EM structural biology studies, as well as recently identified clinically relevant compounds like VBJ103^61,62^ and PAP.^22^ These compounds were chosen for their chemotype diversity and relevance in current TRPM8 research.

The MDS clustering algorithm uses a chemical structure representation that is input in ASCII format via the Simplified Molecular-Input-Line-Entry System (SMILES) and is suitable for efficient in silico analysis.^60,63^ Similarity is evaluated in matrix form based on compound atom descriptor features using the Tanimoto coefficient.^60,64^ MDS output visually represents distances (dissimilarities) between compounds, with the first axis (V1) capturing the greatest variance in MDS space.

Similarity clustering MDS analysis of the 25-compound virtual TRPM8 library is shown in Figure 2A. Assigning accepted TRPM8 functional attributes of agonist or antagonist to MDS scatter plot identifies that along the primary variance axis, V1, cheminformatic similarity analysis generally discriminates agonist from antagonist. We note that icilin is an apparent outlier. MDS reflects a simplified representation of higher dimensionality space; primarily chemical structure features in this case. Accordingly, the icilin peculiarity in MDS space is complicated; however, we note that icilin is reported to require Ca^2+^ as a cofactor for TRPM8 activation, so it does have unusual functional features.^25,65^ Nevertheless, cheminformatic analysis of TRPM8 ligands identifies a relationship between chemical structure and functional activity.

**Figure 2.**
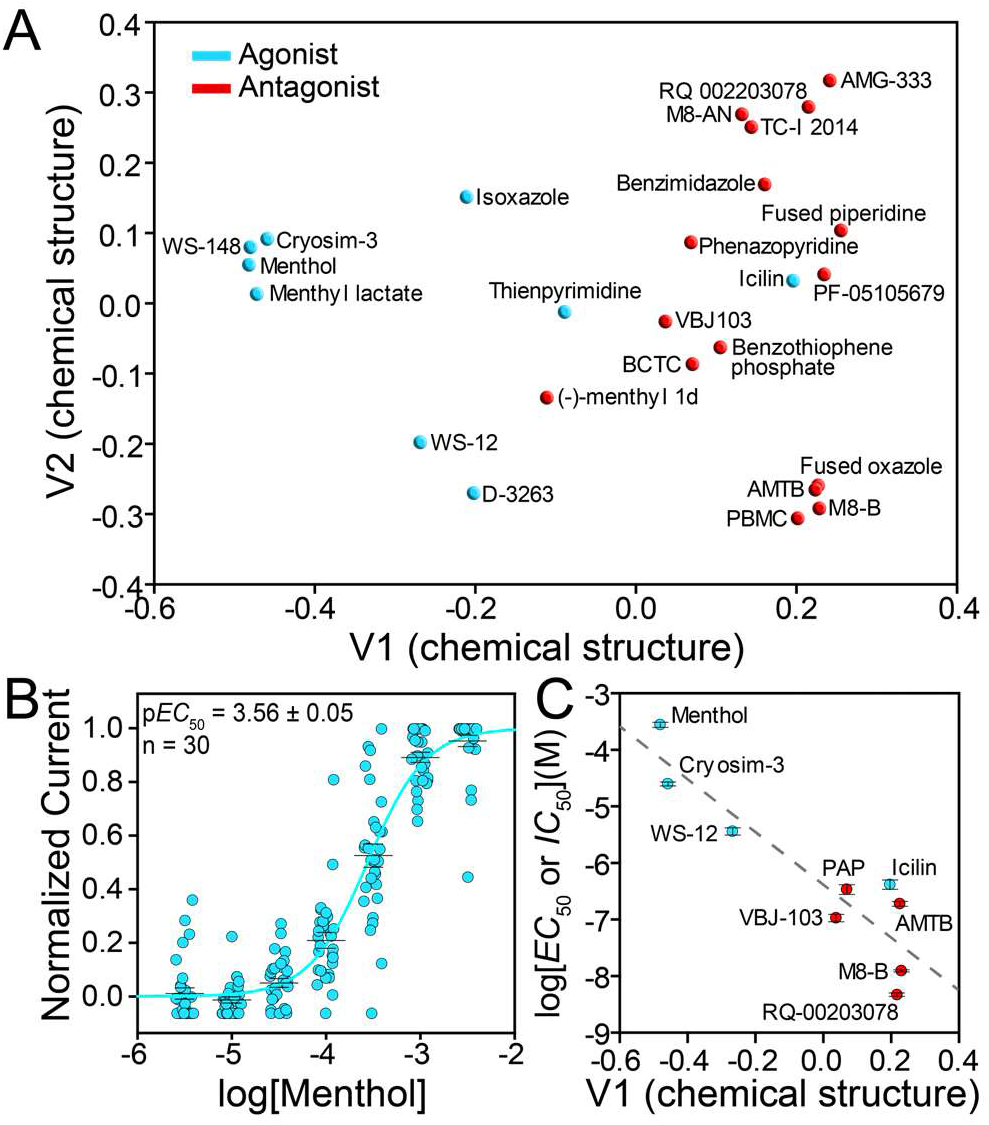
Cheminformatic analysis of TRPM8 chemical compounds is predictive of function and correlates with experimental cellular data. (A) Evaluation of chemical structure in MDS space by similarity clustering is predictive of agonist (blue) and antagonist (red). (B) Menthol activaton of TRPM8 data obtained from automated patch-clamp electrophysiology and (C) correlation plot between functional data (*EC*_50_ and *IC*_50_) and cheminformatic MDS primary variance axis V1 component (R^2^, 0.826; p-value, 0.0007)

To evaluate the cheminformatic MDS similarity clustering quantitatively, we used the k-nearest neighbor (KNN) machine learning algorithm. KNN is a supervised learning classifier that was trained on the 25 TRPM8 compound library (Figure S2) to identify agonists and antagonists. A quantitative assessment of the cheminformatics similarity clustering was done by k-fold cross-validation, which calculates the classification loss by training library subsets. The k-fold cross-validation loss was 0.08, suggesting good performance for a small library and that the cheminformatics analysis chemical structure is predictive of TRPM8 function.

### Quantifying the correlation between cheminformatic chemical structure analysis and cellular function

To further validate and quantify the correlation between chemical structure and agonism or antagonism experimentally, we generated a standardized functional data set using a single-cell automated patch-clamp electrophysiology (APC) of full-length hTRPM8 in a model human cell line. Eleven compounds from the cheminformatic library were selected to obtain human TRPM8 *EC*_50_ and *IC*_50_ values. The need for a standardized data set for quantitative analysis is exemplified by the wide range of literature-reported menthol *EC*_50_ values, which vary between 3 µM and 196 µM ^29,65–71^. The compounds chosen for functional analysis reflect chemical diversity and availability. *IC*_50_ values were determined in the presence of 500 µM menthol stimulation. A minimum of 13 replicates were performed for each compound, with each replicate assessing seven compound concentrations. Each data point from each replicate was averaged from seven current measurements derived from voltage ramps beginning 25 s after exposure at each compound concentration (Figure S3). The current for each point was taken from the difference in current at 100 mV and the holding potential at –60 mV. The cognate menthol agonist data are shown in Figure 2B. Of the 11 compounds evaluated (Figures S4 and S5), two compounds were not well-behaved in our hands. BCTC solubility prevented complete inhibition of menthol-evoked currents, and the agonist D-3263 did not wash out. The standardized APC *EC*_50_*/ IC*_50_ values are tabulated in Table 1.

**Table 1:** p*EC*_50_/*IC*_50_ of TRPM8 Compounds from Automated Patch-Clamp Electrophysiology.

| Compound | $pEC_{50}/pIC_{50}$<br>($EC_{50}/IC_{50}$) | Replicates |
| --- | --- | --- |
| Menthol | **3.56 ± 0.05**<br>(276 μM) | **30** |
| Icilin<sup>a</sup> | **6.39 ± 0.08**<br>(412 nM) | **13** |
| WS-12<sup>a</sup> | **5.44 ± 0.06**<br>(4 μM) | **13** |
| Cryosim-3<sup>a</sup> | **4.60 ± 0.03**<br>(24.8 μM) | **16** |
| D-3263<sup>a,b</sup> | n/a | **4** |
| AMTB<sup>c</sup> | **6.72 ± 0.04**<br>(188 nM) | **12** |
| M8-B<sup>c</sup> | **7.91 ± 0.02**<br>(12 nM) | **12** |
| PAP<sup>c</sup> | **6.5 ± 0.9**<br>(338 nM) | **12** |
| VBJ103<sup>c</sup> | **7.0 ± 0.7**<br>(106 nM) | **14** |
| RQ-00203078<sup>c</sup> | **8.33 ± 0.03**<br>(5 nM) | **14** |
| BCTC<sup>c,d</sup> | n/a | **17** |

<sup>a</sup>Agonist, <sup>b</sup>The D-3263 compound had issues with washing out which made it difficult to do leak subtraction (baseline correction), <sup>c</sup>antagonist, $IC_{50}$ in the presence of 500 μM menthol, <sup>d</sup>data shown in supporting estimated $IC_{50}$ was >100 mM, <sup>d</sup> Full inhibition was unable to be reach as a result of solubility issues.

To quantitatively evaluate the relationship between chemical structure and TRPM8 function, we plotted the resulting *EC*_50_ and *IC*_50_ values against the V1 primary variance axis from cheminformatic similarity analysis. The hybrid data plot was fit to a linear regression model (Figure 2C). The simple coefficient of determination (R^2^) metric was used to quantify the relationship between cellular electrophysiology function and chemical species. The R^2^ value between *EC*_50_/*IC*_50_ cellular functional data and chemical structure cheminformatics values indicates that the majority of variance (82.6%) is explained in the regression model and establishes a straightforward and quantitative relationship between TRPM8 cellular function and in silico ligand analyses. A p-value of 0.0007 supports the conclusion that the correlation between chemical structure and cellular function is meaningful.

The connection between TRPM8 chemical structure and protein function and opacity from current cryo-EM TRPM8 structural biology suggests that protein dynamics may play an important role in TRPM8 function. Structural biology is traditionally thought to bridge a gap between molecular details and function. Given the paucity of TRP channel X-ray crystallography structures, including none for any full-length TRPM family channel, we hypothesized that TRPM8 dynamics might play an outsized functional role. To test this hypothesis, we established a system to detect changes in the TRPM8 conformational ensemble that arise from the presence of diverse chemical ligands.

### The isolated human TRPM8 VSLD resides in a biologically relevant state

Previous NMR studies of the heat, capsaicin, and proton receptor TRPV1 extensively characterized and identified that the isolated hTRPV1-VSLD was folded with the correct membrane topology, agonist binding competent, and that temperature-dependent conformational changes could be leveraged to make predictive mutations that modulate TRPV1 cellular function.^72^ Similarly, NMR and microscale thermophoresis-based menthol and WS-12 binding studies of the isolated human TRPM8-VSLD, pre-S1, and 9^th^ helix turn helix (HTH9) regions (residues Pro672-Pro855) validated electrophysiology results that this region contained the orthosteric binding site prior to TRPM8 cryo-EM structures.^34^

Building upon these results, we engineered a human TRPM8-VSLD construct to investigate ligand-induced protein dynamics. The construct encompasses the pre-S1 amphipathic helix through the S4 transmembrane helix (Pro716-Pro855). Heterologous human TRPM8-VSLD expression was evaluated under 63 initial conditions, which include three IPTG concentrations (0.3 mM, 0.5 mM, and 1 mM), three growth temperatures (18 °C, 25 °C, and 37 °C), and seven *E. coli* expression strains. Expression levels were quantified by antibody-detected dot blot analysis (Figure S6 and Table S4). Outcomes of the first round of expression identified an initial set of expression conditions. A focused second round of expression optimization fine-tuned the IPTG induction concentration, providing the final expression conditions of 18 °C growth temperature, 0.2 mM IPTG to induce expression from the BL21 (DE3) Lobstr cell line (see methods for details).

Optimized expression and purification results in pure and homogenous hTRPM8-VSLD (Figure S7) reconstituted in lysopalmitoylphosphatidylglycerol (LPPG) lysolipid micelles. NMR ^1^H-^15^N TROSY-HSQC spectra show well-resolved resonances with good proton dispersion, supporting that the human TRPM8-VSLD is well-folded (Figure S7C). Far-UV circular dichroism (CD) data further confirms the expected α-helical nature of the VSLD (Figure S7D). To confirm that the isolated hTRPM8-VSLD is in a biologically relevant state, we first assessed menthol binding. A ^1^H-^15^N TROSY-HSQC NMR-detected titration was recorded at physiological temperature (37 °C; Figure 3A). Chemical shift perturbation (CSP) for several resonances was observed with increasing menthol concentrations, consistent with ligand binding. These data were further analyzed using principal component analysis (PCA) of the NMR spectra.^73^ Principal component 1 (PC1) was plotted agonist menthol concentration and fit to a specific binding model. The binding affinity (*K*_d_) for menthol was determined to be 85 ± 25 mmol % (Figure 3B).

**Figure 3.**
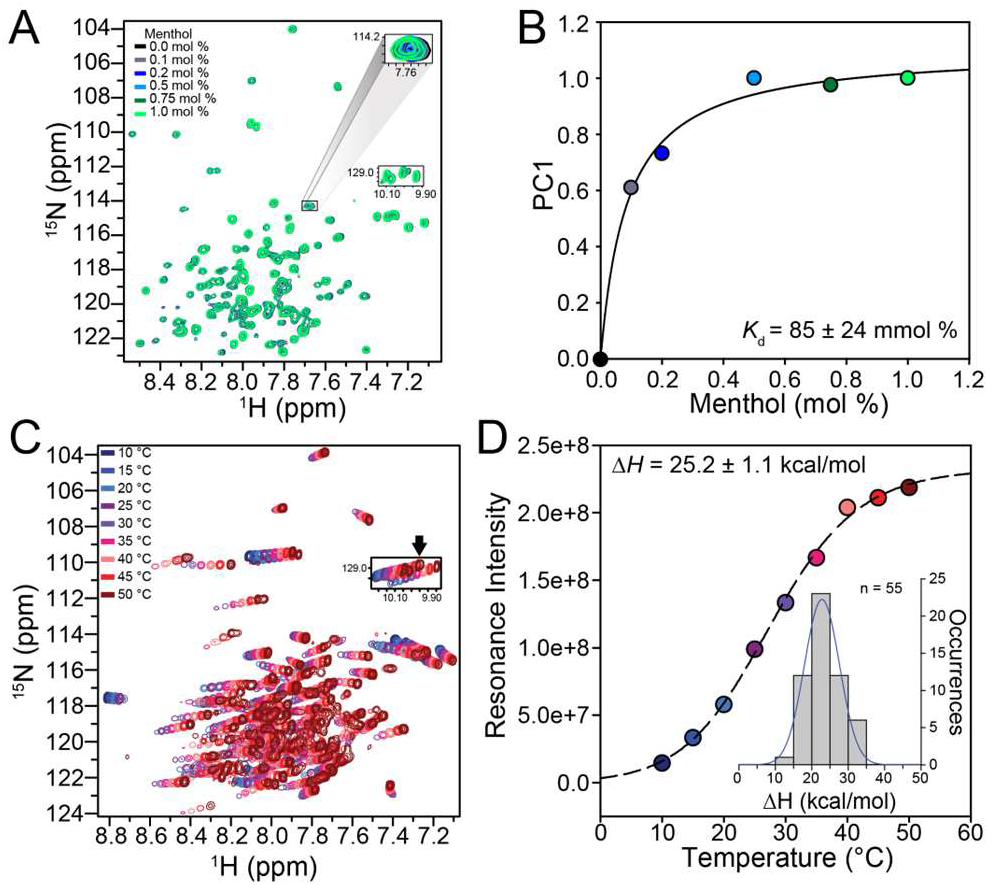
The isolated hTRPM8 VSLD in solution retains key biologically relevant features. (A) NMR-detected titration of the VSLD with the cognate agonist menthol identifies chemical shift perturbations consistent with binding. (B) Global analysis of the titration data in (A) shows specific binding to menthol consistent with the VSLD retaining the orthosteric binding pocket. (C) An NMR-detected temperature titration shows chemical shift perturbations that, when analyzed (D), identify a two-state temperature-dependent conformational change indicative that the VSLD contributes to cold-sensing and retains this phenotype in the isolated state. Inset in (D) is analysis of *ΔH* for 55 resonances, the mean of which is 22.8 ± 0.1 kcal/mol.

To evaluate if the TRPM8-VSLD displays similar temperature-dependent behavior as TRPV1-VSLD, as previously predicted^74^, we collected an NMR-detected temperature titration. 2D ^1^H-^15^N TROSY-HSQC spectra were collected from 15 °C to 50 °C in 5 °C increments (Figure 3C). Chemical shift trajectories for 55 resonances were unambiguously trackable, and the intensities of these resonances at each temperature were measured. To quantify an estimate of the thermosensitivity, we fit the data to a two-state sigmodal model to extract the change in enthalpy (ΔH), an established proxy of the temperature dependence of a conformational change.^18,72^ The aggregated 55 ΔH values were fit to a Gaussian distribution to obtain a mean thermosensitivity (ΔH_avg_) of 22.8 ± 0.1 kcal/mol (Figure 3D). The results of the NMR temperature titration were further validated using intrinsic tryptophan fluorescence, which was probed from 6.5 °C to 44.6 °C in ∼5 °C increments. The fluorescence data yielded an ΔH of 23 ± 8 kcal/mol (Figure S8). The relatively high thermosensitivity of the hTRPM8-VSLD suggests a role beyond housing the orthosteric site, though further characterization is beyond the scope of this study.

### hTRPM8-VSLD ligand-dependent conformational ensemble mirrors activation and inhibition

To probe underlying differences in protein dynamics associated with ligand binding, we employed ^19^F NMR, which provides high sensitivity to subtle conformational changes.^75,76^ A non-perturbing ^19^F label was biosynthetically incorporated to label four endogenous tryptophan residues. The 5-fluorotryptophan residues are located in pre-S1 (W725), S1 (W740), S2 (W786), and S3 (W798) of the hTRPM8-VSLD (Figure 4A). These probes are serendipitously located either in the same transmembrane helices or proximally, within 6 Å (Cα), to key residues (e.g., Y745, R842, D802, and G805) in the orthosteric binding site. The apo human TRPM8-VSLD ^19^F NMR spectrum shows four peaks corresponding to the expected four labeled tryptophan residues (Figure 4B). Peaks were unambiguously assigned by mutagenesis, where each tryptophan was mutated to phenylalanine (W725F, W740F, W786F, and W798F), and the resulting mutant spectrum was used to assign peaks (Figure S9). Distributed across TRPM8-VSLD, these probes report on the hTRPM8-VSLD dynamics. Insight into protein dynamics arises partly from the sensitivity of the chemical shift information, which reflects protein structural states or the ensembles of conformational states.^41,42,77^ ^19^F NMR resonances represent an average of the ensemble of states that is biased by ligand binding or other stimuli.

**Figure 4.**
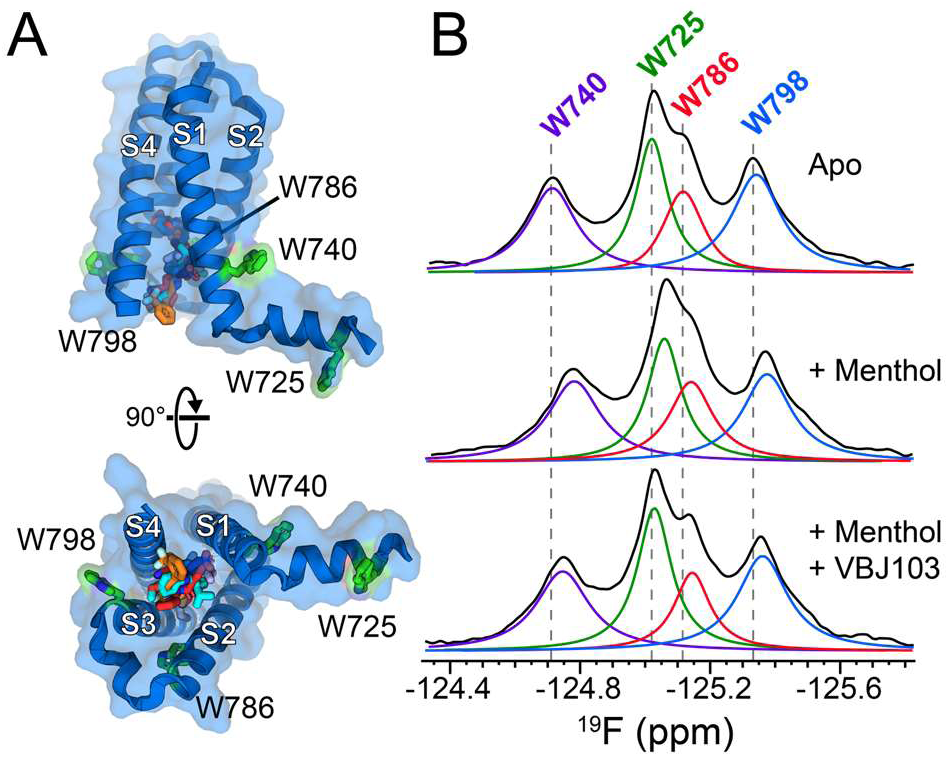
^19^F NMR TRPM8 VSLD spectral changes reflect chemical binding from an agonist and antagonist. (A) Structural representation of the TRPM8 VSLD highlight four tryptophan residues (green), which are located in the pre-S1 (W725), S1 (W740), S2 (W786), and S3 (W798) helices (pdb: 7WRA). The endogenous tryptophan residues are well positioned to report on the orthosteric menthol binding site. (B) 5-fluoro-tryptophan ^19^F NMR spectra of hTRPM8-VSLD in the presence of menthol and the inhibitor VBJ103. Menthol induces chemical shift perturbation in all resonances, suggestive that of an activated state conformational ensemble is detected with the ^19^F probes. Addition of VBJ103 in the presence of menthol shift resonance positions towards the apo state consistent with an inhibited conformational state that reflects the apo/resting state.

To validate that ^19^F NMR data recapitulated features of traditional ^1^H-^15^N NMR biomolecular experiments, we determined *K*_d_ values for four compounds (menthol, icilin, (-)-menthyl 1d, and VBJ103; Figure S10) and evaluated thermosensitivity. A series of ^19^F NMR spectra were collected (Figure S10A) and analyzed using PCA. The binding affinities measured by ^19^F NMR and ^15^N-HSQC generally agree for menthol, icilin, VBJ103, and (-)-menthyl-1d (Figures S10B-E). We additionally measured the *K*_d_ for WS-12 and D-3263 with ^19^F NMR (Figure S10F, G). ^19^F NMR-detected thermosensitivity (ΔH) was also in agreement with ^1^H-^15^N HSQC and intrinsic tryptophan fluorescence. ^19^F-labeled hTRPM8-VSLD NMR spectra were collected from 15 °C to 50 °C in 5 °C increments (Figure S11A). The resulting spectra were deconvoluted, and temperature-dependent peak intensities were obtained. However, due to extensive overlap between the W725 and W786 resonances, these intensities could not be accurately measured. Nevertheless, the W740 and W798 resonances were well-resolved and accurately deconvoluted. W740 and W798 have thermosensitivity values of 20 ± 4 and 34 ± 13 kcal/mol (Figure S11B, Figure S11C, respectively). The ^19^F-derived ΔH values agree with those from ^1^H-^15^N TROSY-HSQC and intrinsic fluorescence (Figure S11D). These data show that hTRPM8-VSLD binding and thermosensitivity values obtained from ^19^F probes reflect those obtained from traditional ^1^H-^15^N backbone experiments and validate the use of ^19^F NMR studies.

A feature of TRPM8 function is an interdependence between activation modes. For example, it has been shown that menthol alters TRPM8 cold activation.^67^ To evaluate if the origins of this feature could be traced to the TRPM8-VSLD, we evaluated coupling between menthol binding and cold-dependent conformational changes by measuring the menthol *K*_d_ over a range of temperatures (15 °C to 45 °C in 5 °C increments). Temperature-dependent affinity analysis reveals a non-linear van’t Hoff plot (Figure S12A), which suggests that menthol activation is allosterically modulated by temperature within the VSLD.^74^

Lastly, to further tie ^19^F spectral and functional features, a series of ^19^F-detected hTRPM8-VSLD NMR spectra were qualitatively evaluated where the apo state spectrum was compared to that in the presence of the cognate agonist menthol and in the presence of the potent antagonist VBJ103 (Figure 4B). Menthol and VBJ103 with menthol were added at saturating concentrations (∼2.5× *K*_d_). The addition of menthol induced chemical shift perturbations in all four tryptophan resonances, with the most significant chemical shift changes occurring in W740 and, to a lesser degree, in W725, W786, and W798 (Table S5). Adding the VBJ103 antagonist in the presence of menthol reverts the spectral changes towards the apo-like spectrum. Presumably, these spectral features reflect that VBJ103 induces changes in protein dynamics away from the menthol-bound activated ensemble and towards a more apo-like conformation state. These data also suggest that VBJ103 competitively binds hTRPM8-VSLD to stabilize an inactive-, resting-, or closed-like state. These ^19^F NMR-detected ligand titration data capture subtle spectral changes that arise from ligand-dependent differences in the conformational ensemble, effectively enabling us to probe hTRPM8-VSLD functional dynamics.

### hTRPM8-VSLD conformation dynamics correlates with ligand structure cheminformatics

To establish if there is a relationship between chemical structure and hTRPM8-VSLD protein dynamics, we rationally generated a menthol-derived set of agonists and antagonists (Figure 5A). The set includes three agonists (menthol, menthyl lactate, and WS-12) and two antagonists ((-)-menthyl 1d and VBJ103).^61,78^ We also included icilin as a control for a divergent non-menthol-based chemotype. Cheminformatic hierarchical clustering validated and quantified the rationally expected chemotype relationships (Figure 5A). Cheminformatics similarity clustering of this menthol-derived compound set is shown in Figure 5B and corroborates the designed library.

**Figure 5.**
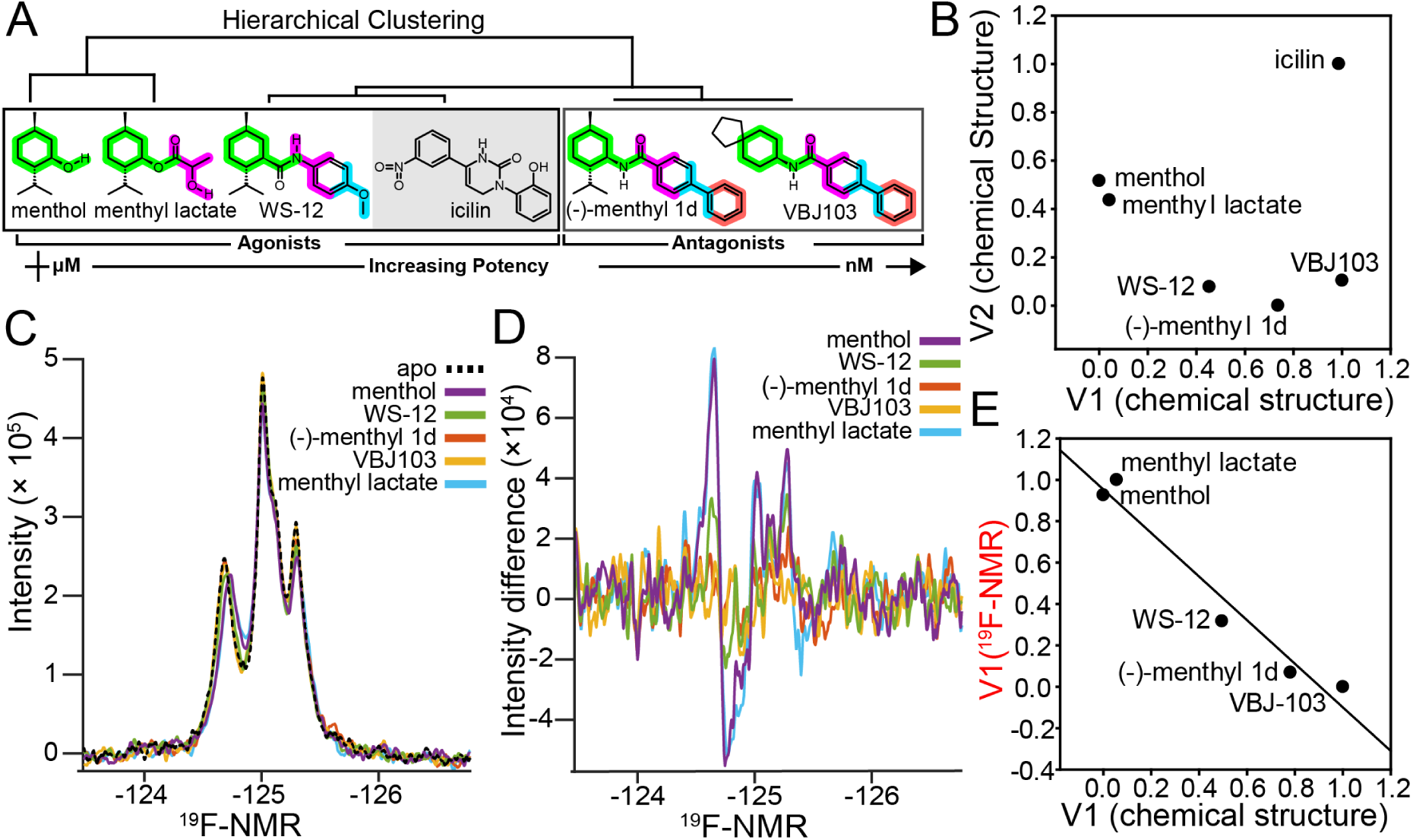
^19^F NMR-detected conformation dynamics correlate with ligand chemical structure. (A) Chemoinformatic hierarchical clustering similarity tree validates the relationship of a rationally designed chemical library that builds on a menthol-based chemotype. Note that icilin is included as a non-menthol control. (B) Menthol-derived clustering of the library in (A) by multidimensional scaling (MDS) recapitulates the chemotype trajectory. (C) 5-fluoro-tryptophan ^19^F NMR spectra were collected at saturating concentrations for the chemotype library. (D) ^19^F NMR difference spectra produced by subtracting the apo-hTRPM8-VSLD spectrum from all ligand-bound spectra to produce a dissimilarity matrix for MDS analysis. (E) MDS data show a strong correlation (R^2^, 0.957; p-value, 0.0038) between chemical structure and ^19^F NMR reported conformational dynamics.

To evaluate if protein dynamics correlates with the ligand structure, ^19^F hTRPM8-VSLD NMR spectra were collected at 37 °C in the absence of ligand (apo-state) and at an estimated saturating ligand concentration (∼2.5× *K*_d_) of the menthol-related compound set that included menthol, menthyl lactate, WS-12, (-)-menthyl 1d, and VBJ103 samples (Figure 5C). To quantitatively assess the effects of the protein dynamics that arise from this set of TRPM8 ligands, ^19^F NMR difference spectra were generated by subtracting the apo-state spectrum from the respective ligand-bound equivalent sample (Figure 5D). The NMR difference spectra were used to generate a dissimilarity matrix and subjected to MDS similarity clustering of the experimental data analogously to the cheminformatic analysis done earlier (Figures 2A and 5B). The chemical structure-based primary variance axis (V1, Figure S13A) was plotted against the primary variance derived from the experimental ^19^F NMR data (Figure S13B) to establish a correlation between chemical structure and hTRPM8 protein dynamics (Figure 5E). A clear linear relationship appears in the data, which was fit to a linear model where the R^2^ coefficient of determination establishes a strong correlation of 0.957 and a p-value of 0.0038. We interpret this data to mean that protein dynamics can potentially be leveraged to identify which compound is engaged in binding. Alternatively, because of the correlation, a chemical structure can be leveraged to make predictions about the ^19^F NMR-detected protein dynamics.

### Predicting conformational dynamic signatures from chemical structure

To validate the predictive relationship between small molecule structure and hTRPM8-VSLD protein dynamics, we used MDS similarity clustering to identify D-3263. This menthol-related TRPM8 agonist has been evaluated clinically to treat prostate hyperplasia.^79^ This analysis revealed that D-3263 shares the highest similarity in MDS space with WS-12 (Figure S13C). We hypothesized that D-3263 would modulate the protein dynamics of the TRPM8-VSLD in a way that is most similar to that of WS-12. To test this, we collected ^19^F NMR hTPRM8-VSLD spectra (Figure 6A) and subjected them to the same similarity clustering protocol in MDS space (Figure S13D) using the difference NMR spectra (Figure 6B). Comparison of the principal variance axes from the chemical structure and experimental ^19^F NMR-based protein dynamics were assessed and fit to a linear model (Figure 6C). As hypothesized, these data support that D-3263 conformationally selects TRPM8 dynamics in a manner most similar to the cooling agonist WS-12.

**Figure 6.**
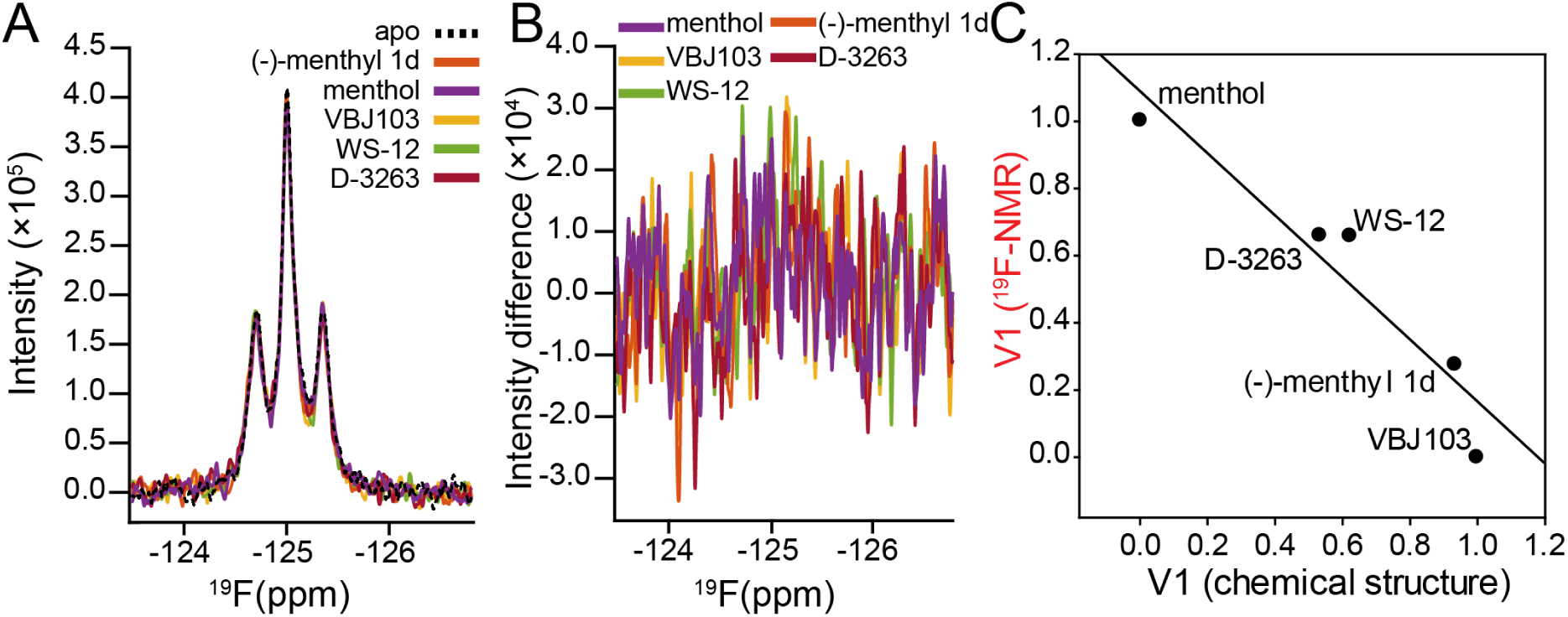
Predicting NMR-detected chemometrics beyond a menthol-based chemotype. Decreased protein concentration to challenge and include D-3263. Second round of chemometric analysis predicts D-3263. (A) lower concentration ^19^F NMR spectra collected at saturated agonist and antagonist. (B) Apo-hTRPM8-VSLD spectrum were substracted from all ligand-bound spectra to obtain difference matric and MDS were implemented on the spectra difference data.(C) MDS data show strong correlation (R^2^, 0.815; p-value, 0.0358) between structure and conformational dynamics and identifies D-3263 as situated closer to WS-12.

As part of this predictive experiment, we deliberately decreased the hTRPM8-VSLD concentration by ∼30 % to lower the signal-to-noise of the data and test the limits of the correlation. Comparison of the difference spectra in Figures 5D and 6B illustrate the decreased signal-to-noise. Despite this, the correlation between chemical structure and protein dynamics remains, albeit with a lower correlation. Nevertheless, the linear model explains 81.5% of the correlation (Figure 6C). Notably, the second round of NMR studies validates the predictive nature between small molecule structure and protein dynamics.

### Extending and validating the link between NMR-detected protein dynamics and preclinically relevant TRPM8 chemical scaffolds

To further validate and assess the robustness of the relationship between chemical structure and protein dynamics, we expanded the hybrid computational and experimental analysis to include compounds with substantial chemotype diversity. We performed a third round of ^19^F NMR experiments, which included menthol, WS-12, and VBJ103 as potential anchors and expanded the list also to include cyrosim-3, phenazopyridine (PAP), BCTC, AMTB, and M8-B.

A fundamental assumption of the previous NMR analyses is that principal axes of variance (V1) reflected the hypothesized relationship to TRPM8 conformation dynamics. To validate that assumption, we opted to vary the percentage of DMSO in our samples to introduce spectral perturbation bias (Figure 7A), an experimental aspect that had been controlled in the previous analyses. It has been established that DMSO can alter ^19^F NMR spectra.^80–83^ To evaluate the perturbation of DMSO on the TRPM8 VSLD, a control experiment was done where the addition of a small amount of DMSO (0.63%, Figure S14) was sufficient to induce small chemical shift changes (∼Δδ: 0.007-0.016) and a 5-8% decrease in peak intensity. We hypothesized that varying the DMSO concentration deliberately, by up to 3 fold, will induce spectral changes orthogonally to the data features that arise from ligand-protein dynamics selection. Additionally, because the magnitude of the DMSO-induced spectral changes is larger than those that arise from compound addition–conformational selection, the principal variance axis will poorly correlate with chemical structure. As expected, the NMR-derived MDS principal variance axes (V1, Figure 7B) showed a weak correlation with chemical structure and ^19^F NMR spectra (R^2^, 0.398, Figure S15). However, as predicted, the V2 MDS component, which reflects the secondary variance axis of the MDS analysis, has a strong correlation with chemical structure (R^2^, 0.880; p-value, 0.006; Figure 7C,). We ascribe the low correlation between V1 components to the different concentrations of DMSO in these samples. Nevertheless, the ^19^F NMR experimental data still retain the functional dynamics information, albeit in the second variance axis component (V2). Chemometric analysis of the chemical structure and ^19^F NMR protein dynamics signatures are sufficiently robust to overcome systematic external perturbation, even in the presence of diverse chemotypes.

**Figure 7.**
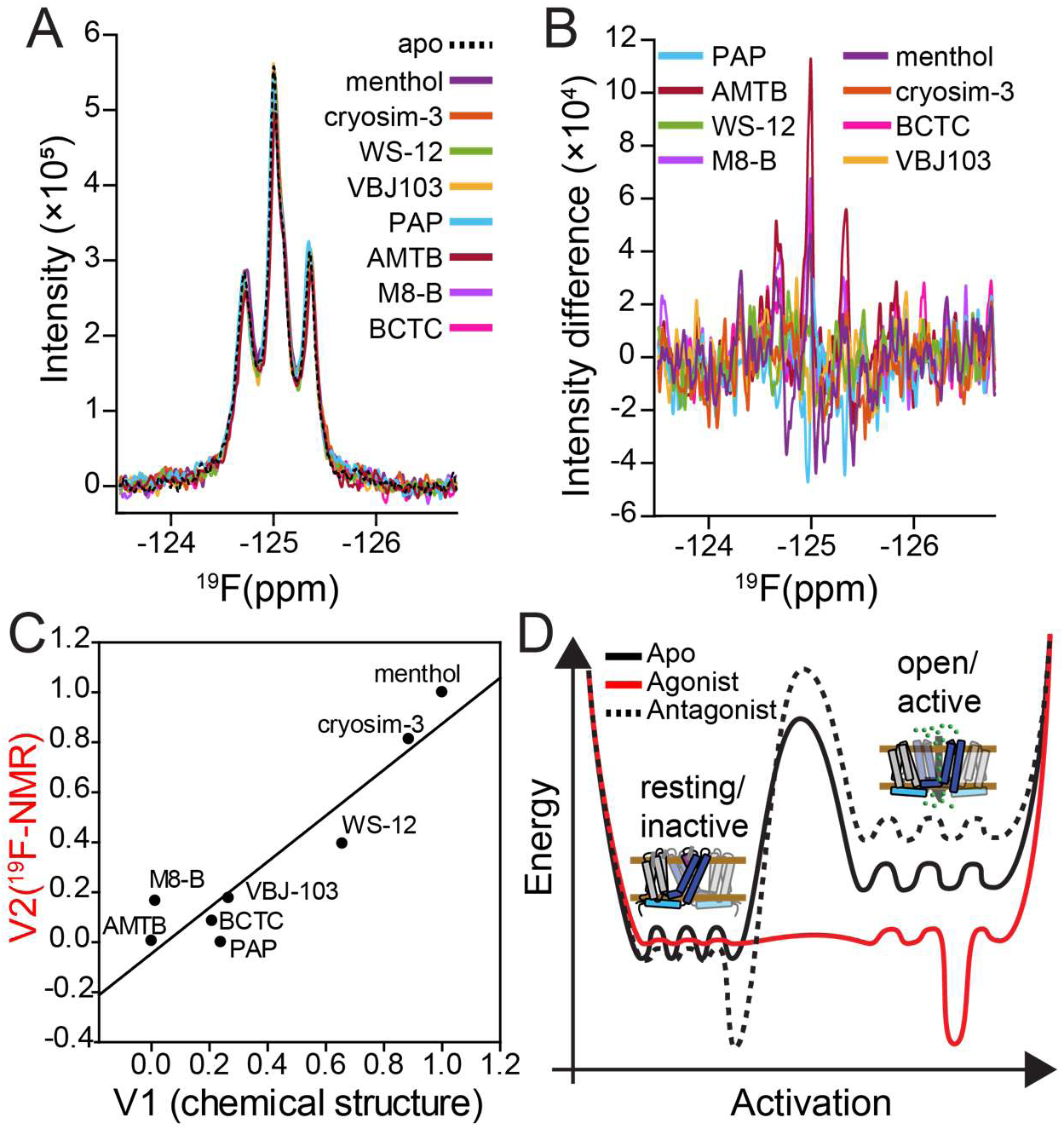
Validation of the correlation of NMR-detected chemometrics for diverse chemotypes. Third round of chemometric analysis includes compounds beyond menthol derivatives. (A) ^19^F NMR spectra in the presence menthol, WS-12, VBJ103, cryosim-3, PAP, AMTB, M8-B and BCTC. In addition, variability in DMSO to validate to force compound dependent variation into component V2. (B) Difference spectra used for MDS analysis. (C) Correlation plot between conformational dynamics and chemical structure show a strong correlation (R^2^, 0.880; p-value, 0.006). (D) A proposed TRPM8 energy landscape where relatitvley accessible substates underlie small magnitude conformational changes that efficiently interconvert enabling funcational dynamics by conformaitonal selection.

### Insights into TRPM8 protein dynamics

A primary assumption in this study is that protein dynamics is what underlies the bridge between the ^19^F NMR data and ligand structure. To test this hypothesis, we measured dynamics directly. The ^19^F NMR data herein shows individual resonances and subtle chemical shift position changes upon the addition of a variety of ligand chemotypes (c.f. Figures 5C, 6A, and 7A), which is consistent with TRPM8 VSLD conformational exchange rates in fast to intermediate exchange regime on the NMR timescale. To quantitatively validate that ligands modulate hTRPM8 protein dynamics, we conducted a series of ¹⁹F-CPMG relaxation dispersion experiments of the hTPM8 VSLD in the apo, agonist-bound, and antagonist-bound states.^84^ The agonist used was menthol, the canonical TRPM8 activator, which is placed structurally in the orthosteric binding pocket by the chemically related WS-12 cryo-EM structure.^25^ The antagonist AMTB is also structurally resolved to the orthosteric VSLD binding site (Figure S16A).^28,30^ CPMG experiments were performed in triplicate at 600 and 850 MHz using CPMG frequencies ranging from 250 Hz to 5000 Hz to generate relaxation dispersion profiles. Of the four ^19^F resonances in the VSLD, only W740 showed a dispersion profile. The lack of dispersion profiles for W725, W786, and W798 is consistent with fast exchange and consistent with previous studies.^55^ While W740 is not directly involved in ligand binding, it is very close to the binding site. W740 is located in the S1 helix, on the opposite side of the helix, pointing away from the orthosteric site (Figure S16A). The W740 relaxation dispersion curves were fit to extract the chemical exchange rate (*k*_ex_). The fitted dispersion curves for the apo state, antagonist-bound, and agonist-bound conditions revealed ligand-dependent changes in *k*_ex_ (Figure S16B-D). Specifically, the basal apo state *k*_ex_ was measured as 7×10^3^ ± 3×10^3^ s⁻¹, which increased to 12×10^3^ ± 4×10^3^ s⁻¹ in the presence of AMTB antagonist, and decreased to 4×10^3^ ± 3×10^3^ s⁻¹ with menthol binding. These results support that ligand binding modulates the rate of conformational exchange of a ^19^F W740 probe, which resides near the binding site.

To further evaluate the role of TRPM8 dynamics and validate the CPMG data, we performed 2D lineshape analysis on ¹H-¹⁵N TROSY-HSQC-detected titrations of agonists (menthol and icilin; Figure S17A-B) and antagonists (VBJ103 and (-)-menthyl 1d; Figure S17C-D) titration using the TITAN software package.^85,86^ Currently, backbone NMR resonance assignments do not exist for the hTRPM8 VSLD, so residue identification is not possible. However, we performed lineshape analysis of nine representative well-isolated peaks distributed across the titration spectra (Figure S17E). It is well established that exchange kinetics contribute to NMR transverse relaxation rates (*R*₂), which is manifest in resonance linewidth (λ). We fit individual resonances (Figure S17F) and quantified the change in linewidth (Δλ) as a probe of protein dynamics. Linewidths in both direct and indirect dimensions were extracted and Δλ values were calculated (Table S6 and S7).

Menthol binding resulted in minor changes in linewidth with an average difference (Δλ_AVG_) for all nine selected peaks of 2 ± 8 Hz, consistent with minor changes in backbone dynamics. The more potent agonist icilin caused general resonance sharpening (–172 ± 11 Hz), suggesting a shift toward a faster exchange regime for the backbone amides. In contrast, the antagonist VBJ103 caused substantial line broadening with a Δλ_AVG_ of 154 ± 7 Hz, consistent with a shift towards decreased exchange regimes. A similar trend was observed for the antagonist (-)-menthyl 1d. Together, the ¹⁹F-CPMG and 2D lineshape analyses offer complementary perspectives of TRPM8 protein dynamics. The ¹⁹F probe on the W740 sidechain, near the orthosteric binding site, reports that agonist binding decreases *k*_ex_, suggesting slower exchange rates at the sidechain level near the binding site. Whereas antagonist binding increases *k*_ex_, implying faster sidechain motions for W740. Conversely, the lineshape analysis, which is sensitive to backbone motions, reveals that agonists (particularly icilin) can increase backbone exchange rates, while TRPM8 antagonists generally attenuate dynamics. These nuanced findings suggest that ligands differentially affect sidechain and backbone dynamics, highlighting the complexity of conformational modulation in hTRPM8.

A potential explanation for differential effects of agonist and antagonist on backbone and side chain dynamics could be attributed to entropy–entropy compensation, but requires further investigation.^56,87^ Ligand binding can significantly perturb local changes in side-chain dynamics that are balanced by adjustments in flexibility elsewhere (near or distant).^55,56,87,88^ Moreover, internal conformational entropy measurements have shown high correlation with total binding entropy, supporting the notion that fast-timescale exchange at side-chain may play an important functional role.^88^ In the case of TRPM8, the measured dynamics is in qualitative agreement with the cryo-EM structural biology, which shows relatively small magnitude conformational changes (Figures 1B and S1, Tables S2-3). Figure 7D schematizes a TRPM8 protein dynamics landscape where functional dynamics are accessed through small-scale conformational changes that interconvert relatively rapidly because of low free energy barriers. We note that such a dynamic profile could be a functional requirement for temperature-sensing receptors like TRPM8, which evolved to sense small temperature changes.^1^

### NMR-detected protein dynamics also correlate with cellular function

We have established and quantified the correlation between hTRPM8 function and chemical structure (Figure 2C) and, through three rounds of chemometric analysis, established a robust correlation between hTRPM8-VSLD apparent protein dynamics and chemical structures (Figures 5-7). Under the assumption that ligands impart functional effects by modulating protein dynamics, we evaluated the correlation between protein dynamics and cellular function. A correlation between ligand-biased dynamics and cellular function was established between the electrophysiology and the ^19^F NMR data from the DMSO biasing experiment. The correlation still holds despite the deliberate experimental DMSO bias (R^2^, 0.791; p-value, 0.0074; Figure S18. Accordingly, we conclude that TRPM8 protein dynamics provide insights that bridge the relationship between chemical structure and cellular function.

In an attempt to identify the spectral features that underlie the origins of the protein dynamics–cellular function correlation, we measured discrete ^19^F spectral features. Specifically, we compared *EC*_50_ and *IC*_50_ values from Table 1 with spectral features of peak intensity and linewidth (frequencies) from Figs. 5 and 7 data sets. There is a correlation between both W798 peak intensity and linewidth with efficacy (Figure S19A-D). Curiously, no similar trends were observed for W725, W740, and W786, as shown for W740 in Figure S19E-F. To test if orthosteric site proximity was responsible for the correlation with W798 resonance features and cellular function, we calculated the distances between all bound ligands in the cryo-EM structures (Table S2) and all equivalent positions of the VSLD tryptophan fluorine probes. On average, the fluorine probe in W798 is the closest to orthosterically bound ligands, though only by a small margin (<2Å) relative to the next closest ^19^F probe site W740. While there is a slight distance bias that might underlie the positive correlation with W798, several structures place ^19^F probes of W740 and W786 closer to bound ligands in a structure-dependent manner. Accordingly, we don’t interpret spectral features and functional correlations to be solely a function of ligand proximity. Likely, the relationship is more complex, and the presence or absence of the correlation also depends on the relative locations to the orthosteric site in addition to the timescales and amplitudes of dynamics each ^19^F probe experiences. Future studies are needed to evaluate this hypothesis.

## Conclusion

TRPM8 is a validated drug target for several pathophysiologies. However, its utility has been limited by a purported need for mode-selective thermoneutral molecules. Here, we employ a cheminformatics approach and identify a relationship between molecular structure and NMR-detected TRPM8 functional dynamics that might be useful in circumventing current obstacles. Here, we validated the TRPM8 structure-activity relationship (SAR) between chemical structure and cellular electrophysiology function for TRPM8 (Figure 2). MDS similarity clustering discriminates between agonist and antagonist compounds, thus indicating that chemical structure can be used to predict, at least coarsely, between agonist and antagonist. We further validated the TRPM8–SAR relationship by collecting a set of *EC*_50_ and *IC*_50_ values from automated patch-clamp electrophysiology. The analysis shows a robust correlation between chemical structure and efficacy. Given the validated TRPM8 chemical structure-activity relationship and the cyro-EM structural similarity, we hypothesized that protein dynamics by ligand-based conformational selection might underlie the observed SAR.

Although functional^32,65,66,89^ and structural^24–26,28^ studies have established the orthosteric site, the role of protein dynamics in TRPM8 function has remained opaque. To probe this, we optimized the expression of a human TRPM8-VSLD construct, evaluated it with a traditional solution NMR, and demonstrated that it is binding competent and sensitive to temperature changes. Using ^19^F NMR, we investigated ligand-induced hTRPM8-VSLD protein dynamics. We selected menthol and menthol-derived agonists and antagonists and collected ligand-bound TRPM8-VSLD ^19^F NMR spectra. Our data suggests that small molecules modulate TRPM8-VSLD conformational ensembles consistent with the concept of functional dynamics. Correlations between MDS similarity clustering on small molecule structures and ^19^F NMR spectra, which reflect TRPM8-VSLD protein dynamics, revealed that induced changes in protein dynamics that correlate strongly with chemical structures. Exploiting the correlation between small molecule structure and TRPM8-VSLD protein dynamics, we were able to predict that D-3263 would influence the conformational ensemble analogously to WS-12. Furthermore, we probed the robustness of the correlation between TRPM8 protein dynamics and cheminformatics using diverse chemotypes and perturbing biases.

By combining cheminformatics, automated patch-clamp electrophysiology, and ^19^F NMR dynamics, our study establishes a dynamics-based TRPM8 small-molecule structure-activity relationship. In particular, our ^19^F NMR data show that ligands impart their effect, at least in part, by modulating TRPM8 protein dynamics, which ultimately lead to conformational changes required for channel gating. The integration of protein dynamics with chemical structure and cellular function is anticipated to help efficiently focus the search of chemical space for modulatory compounds with desired functional effects.

## Materials and Methods

### Structural and Cheminformatics Analysis

All chemical structures were analyzed using the *ChemMine Tools* webserver (https://chemminetools.ucr.edu/).^60^ The structural and physiochemical properties of 25 chemical compounds (Figure S2) were retrieved from the PubChem database (https://pubchem.ncbi.nlm.nih.gov/).^90^ Compounds include TRPM8 agonists and antagonists. A default similarity cutoff of 0.4 was used for MDS similarity clustering analysis.

### k-nearest neighbor machine learning evaluation

k-nearest neighbor (KNN) and k-fold cross-validation analysis was performed in Matlab v2022a using the fitcknn and kfoldLoss functions, respectively. The input features for training included the primary and secondary variance axes (V1 and V2) from the ChemMine tools MDS-based similarity clustering and a binary classifier feature to identify agonist or antagonist.^91^ KNN used five nearest neighbors (k), which was defined as 5, and a cosine distance metric. For validation, the number of folds was set to 5.

### Automated Patch Clamp (APC) Electrophysiology

Stably expressing wild type full-length human TRPM8 HEK-293T cells were grown in DMEM (Gibco 11960077) supplemented with 10% fetal bovine serum (Gibco 16000), 4 mM L-glutamine (Gibco 25030), 100 U/mL penicillin: streptomycin (Gibco 15140), 100 µM non-essential amino acid solution (Gibco 11140050), 4 mM glutaMax (Gibco 35050061), 200 µg/mL G418 (Sigma-Aldrich A1720), and 0.12% sodium bicarbonate (Gibco 25080094) and maintained at 37 °C with 8% CO_2_. Cells were cultured in 100 mm dishes. On the day of the experiment, cells were washed twice with phosphate buffer saline solution (PBS), pH 7.4 (Gibco 10010031) before accutase (Gibco A1110501) was added and incubated for 5 minutes at 37 °C, 8% CO_2_. Cells were harvested and transferred to a conical tube and pelleted by centrifuging at 200 *×g* for 1.5 minutes. The supernatant was aspirated. The cell pellet was resuspended in serum-free media (Gibco 11686029) and transferred to a T25 flask and shaken with an orbital shaker at 50 rpm for at least 30 minutes at room temperature to allow for recovery. Before adding the human TRPM8 expressing cells to cell inlet wells of the IonFlux Plate HT – Single Cells (Cell Microsystems 910-0059), cells were centrifuge at 200 *×g* for 1.5 minutes and resuspended in extracellular solution (10 mM HEPES, 145 mM NaCl, 4 mM KCl, 1 mM MgCl_2_, 2 mM CaCl_2_, 10 mM glucose, pH 7.4,) to a cell density of 3-7 × 10^6^ cells/mL. Prior to experimental use, the extracellular solution was pH adjusted with NaOH, and osmolality was measured and adjusted using a Vapro 5600 vapor pressure osmometer (Wescor) and sucrose to 315-330 mOsm. Compounds were dissolved into DMSO before dissolving into extracellular solution. DMSO percentage was kept consistent across dilutions for each compound. Serial dilutions were conducted using the Opentrons OT-2 Robot on 96 well compound plates before being pipetted to single-cell automated patch clamp electrophysiology plates. The cell traps were filled with intracellular solution (10 mM HEPES, 120 mM KCl, 1.75 mM MgCl_2_, 5.374 mM CaCl_2_, 10 mM EGTA, 4 mM NaATP, pH 7.2. The pH was adjusted using KOH and osmolality was adjusted with sucrose to 305-315 mOsm.

Automated patch clamp data were collected using IonFlux Mercury HT with Ionflux HT v5.0. The manufacturer default protocol of “water rinse” and “prime single” was conducted on the plate to wash off the peroxide solution and prime the plate of the solutions and compounds before the experimental protocol, respectively. The experimental protocol is divided into four steps: prime, trap, break, and data acquisition. During the prime step: (1) the main channel was applied 1 psi for t = 0-25 s and 0.4 psi for t = 25-60 s (2) the trap and compound channels were applied at 5 psi for t = 0-20 s and then 1.5 psi for t = 20-55 s followed by only the traps at 2 psi for t = 55-60 s. For the voltage during the prime step, a pulse was applied every 150 ms, where the 0 mV holding potential was applied during t = 0-50 ms, 20 mV was applied during the t = 50-100 ms, and 0 mV during the t = 100-150 ms. During the trap step: (1) the main channel is applied 0.1 psi for t = 0-5 s before applying 0.5 s pulses of 0.2 psi every 5 s during t = 5-135 s (2) the trap channel is applied 6 inHg for t = 0-135 s. For the voltage during the trap step, a pulse was applied every 70 ms, where the –80 mV holding potential was applied between t = 0-20 ms, –100 mV for t = 20-50 ms, –80 mV for t = 50-70 ms. During the break step: (1) the main channel is applied 0.1 psi for t = 0-100 s (2) the trap channel was applied 6 inHg between t = 0-10 s, vacuum ramp from 10 to 14 inHg from t = 10-40 s, and 6 inHg for t = 40-100 s. For the voltage during the break step, a pulse was applied every 150 ms, where –80 mV holding potential was applied between t = 0-50 ms, –100 mV for t = 50-100 ms, and –80 mV for t = 100-150 ms. During the data acquisition: (1) the main channel is applied at 0.15 psi for t = 0-1350 s, (2) the traps channel is applied 5 inHg for t = 0-3 s and 3 inHg for t = 3-1350 s. For the voltage during the data acquisition, a pulse was applied every 625 ms, where the –60 mV holding potential was applied between t = 0-100 ms, – 70 mV for t = 100-200 ms, –60 mV for t = 200-300 ms, a voltage ramp from –120 mV to 160 mV for t = 300-525 ms and –60 mV for t = 525-625 ms. The pulse program for the data acquisition is shown in Figure S3. The compounds were perfused for 75 s for each concentration. *IC*_50_ values were determined in the presence of 500 µM menthol. The data were analyzed using Ionflux Data Analyzer v5.0. The data were leak subtracted based on the initial –60 mV holding potential and –70 mV voltage steps from the data acquisition. The data was averaged from 7 points after 25 s of perfusion of the compound for each concentration, where the points are from the difference of the current at 100 mV and the holding potential at –60 mV.

The data was normalized for each cell and plotted in SigmaPlot 12.5, where it was then fit to a sigmoidal dose-response (variable slope) eq 1. The data was rescaled based on the max and min and refitted.

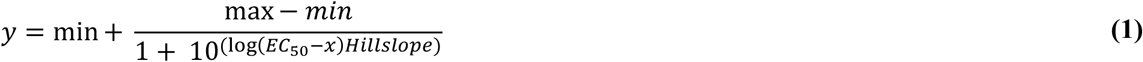

### hTRPM8 VSLD Protein Expression

Wild-type human TRPM8 VSLD (hM8-VSLD, residues 716-855) with an N-terminal thrombin cleavage site (LVPR|GS) and 10×His tag was cloned into pET16b vector. Four mutant hM8-VSLD (W725F, W740F, W786F, and W798F) constructs were generated using Q5 site-directed mutagenesis and verified by Sanger sequencing. WT and mutant hTRPM8-VSLD (W725F, W740F, W786F, and W798F) pET16b plasmid were transformed into BL21 (DE3) Lobstr *E. Coli* expression cell line. Cells were grown in a minimal media 9 [M9; 12.8 g Na_2_HPO_4_·7H_2_O (VWR Life Science), 3 g KH_2_PO_4_ (Thermo Fisher Scientific), 0.5 g NaCl (Thermo Fisher Scientific), 1 mM MgSO_4_·7H_2_O (Sigma Aldrich), 0.1 mM CaCl_2_ (Sigma Aldrich), 0.4% D-glucose (w/v) (Sigma Aldrich), 1x MEM Vitamin solution (100x solution, Corning), 0.5x metals mix (1000x solutions: 10 mM FeCl_3_,10 mM CuSO_4,_ 10 mM MnCl_2_, 10 mM ZnSO_4_), and 1 g NH_4_Cl or ^15^NH_4_Cl (Cambridge Isotope Laboratories) as the sole nitrogen source] and 40 mg 5-Fluoroindole was added per liter 10-15 min before induction for ^19^F labeling of tryptophan residues. Cells were induced at 0.6-0.7 OD_600 nm_ with 0.2 mM isopropyl b-D-1-thiogalactopyranoside (IPTG) and incubated at 18 °C for 36-48 hrs. Cells were harvested at 6,000 *×g* for 15 mins at 4 °C.

### Protein Purification

Cell pellets were resuspended in lysis buffer (75 mM Tris-HCl, 300 mM NaCl, 0.2 mM EDTA, pH 7.7) followed by additions of LDR (0.15 mg/mL lysozyme, 0.015 mg/mL DNase, and 0.015 mg/mL RNase), 1 mM PMSF, and 5 mM magnesium acetate. The lysate was allowed to tumble at room temperature (RT) for 20 min and then sonicated on ice for 8 min with a 55% duty cycle (5 sec on/off) twice. The cell lysate was solubilized with 3% Empigen for 1-2 hr at 4 °C and centrifuged at 38,000 *×g* for 30 min at 4°C. The supernatant was collected and tumbled with Ni-NTA (0.1 ml/g of pellet) for 2 hrs at 4°C. The resin was washed with 25 column volumes of Buffer A (1% Empigen, 40 mM HEPES, 300 mM NaCl, pH 7.5, 2 mM BME) and Buffer B (Buffer A plus 60 mM imidazole). hTRPM8-VSLD was reconstituted into 1-palmitoyl-2-hydroxy-sn-glycero-3-[phospho-rac-(1-glycerol) (LPPG) by performing detergent exchange with Buffer C (0.05 % LPPG, 25 mM Na_2_HPO_4_, pH 7.8) while bound on resin and eluted with 500 mM imidazole in Buffer C. Elution fractions were combined and exchanged into Thrombin Cleavage buffer (20 mM Tris, 100 mM NaCl, pH 8.00) using Millipore Centrifugal Filter Units with 10 kDa cutoff and tumbled in 3 unit of Thrombin for 16-24 hrs at room temperature. Upon cleavage 10×His tag, SDS-PAGE shows a strong hTRPM8-VSLD band shifted down (Figure S7A). Impurities were removed by negative purification with Ni-NTA resin. Flow through and wash from negative purification were concentrated (using Millipore Centrifugal Filter Units with 10kDa cutoff) and further purified using Gel filtration (16 XK GE Healthcare Life Science) in NMR Buffer (25 mM Na_2_HPO_4_, 0.05% LPPG, 0.5 mM EDTA, pH 6.5). SDS-PAGE on size exclusion (SEC) shows a pure, strong, homogenous band (Figure S7B). After gel filtration, hM8-VSLD fractions (lane 8-12, Figure S7B) were pooled, concentrated by ultrafiltration (Amicon, Millipore), and prepared in a 3 mm NMR tube (Bruker) with 5.5 % D_2_O for ^19^F NMR and 2.7 % D_2_O for HSQC experiments. hTRPM8-VSLD concentration was determined using Pierce BCA Protein Assay Kit (ThermoFisher Scientific).

### Far-UV circular dichroism

hTRPM8-VSLD samples for Circular Dichroism (CD) were prepared in an NMR buffer to a final concentration of 0.2 mg/mL. Spectra were collected using a Jasco J-710 spectropolarimeter, with temperature regulation provided by a Peltier device (Jasco). The CD spectra were measured from 190 to 250 nm, using a 100 mDeg sensitivity, a data interval of 0.5 nm, and a scan speed of 50 nm/min. Menthol stock was dissolved in 50% ethanol. Each condition was scanned five times, and the results were averaged for the final spectrum. Blank spectra were recorded in NMR buffer alone or with 10 mol% menthol. Data analysis was performed using CDtoolX.64 software.^92^

### Nuclear magnetic resonance spectroscopy

All spectra data were collected via Bruker Avance II HD 600 MHz ^1^H spectrometer and Avance II HD console equipped with 5 mm Triple Resonance Prodigy Cryoprobe and Bruker 850 MHz ^1^H spectrometer and Avance III HD console equipped with 5 mm TCI Cryoprobe. NMR sample temperatures were calibrated using standard 99% ^2^H methanol to within ∼0.1 °C.

Menthol, icilin, D-3263, (-)-menthyl 1d, WS-12, and VBJ103 titration points were calculated in mol % ^72^, and stocks were prepared in dimethyl sulfoxide (DMSO). The compounds were incubated for 10 min, with hTRPM8-VSLD concentrations ranging from 80 μM to 220 μM in 1.38% LPPG, at 37 °C before data collection. ^1^H-^15^N TROSY-HSQC or ^19^F NMR spectra were collected with 512 to 2048 ns, depending on the sample concentration. For temperature titration, hTRPM8 was incubated at the desired temperature (15°C, 20 °C, 25 °C, 30 °C, 35 °C, 40 °C, 45 °C, and 50 °C) for 10 min. Temperature titration data were fit to eq 2 and 3. ^1^H-^15^N TROSY-HSQC or ^19^F NMR spectra were collected with 1024 to 2048 ns, depending on the sample concentration.

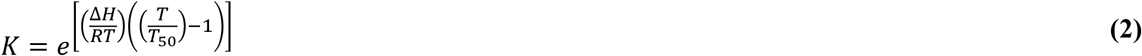

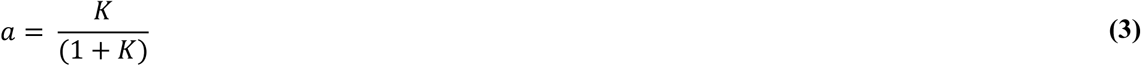

All hTRPM8-VSLD samples used for NMR-detected chemometrics experiments were equally divided to prepare multiple NMR samples (e.g., apo and hTRPM8-VSLD plus compound) in an NMR buffer. hTRPM8-VSLD concentration for round 1, round 2, and round 3 were determined to be 31 μM, 22 μM, and 37 μM in 1.38% LPPG, respectively. All compounds prepared in DMSO or just DMSO were added to hTRPM8-VSLD NMR sample immediately prior to placement into NMR (850 MHz) instrument. All samples were incubated for 10 min at 37 °C before data collection. Spectra were acquired using 2048 number of scans (ns). All Stocks were prepared in DMSO for the following saturating concentrations: menthol (417.5 mmol%), menthyl lactate (270 mmol%), WS-12 (37.5 mmol%), icilin (80 mmol%), D-3263 (405μmol %), VBJ103 (117.5 nmol%), (-)-menthyl 1d (450 nmol%), cryosim-3 (9.4 mmol%), AMTB (93 mmol%), M8-B (62 nmol%), PAP (10 mmol %), and BCTC (6.9 mmol%) were prepared. Saturating concentration for cryosim-3, AMTB, M8-B, PAP, and BCTC were based on an estimate *K*_d_. We used mean literature and APC *EC*_50_ or *IC*_50_ values for menthol, icilin, WS-12, (-)-menthyl 1d, and VBJ103. Functional activity values were compared to ^19^F-NMR based K_d_ values. *EC*_50_ or *IC*_50_-*K*_d_ plot were fitted to a linear modal, yielding a R^2^ 0.722 (Figure S20). Using literature and APC obtained mean *EC*_50_ or *IC*_50_, we determine an estimated menthyl lactate, cryosim-3, AMTB, M8-B, PAP, and BCTC *K*_d_ values. In addition, for round three of chemometric experiments, DMSO content was varied for menthol (3 μL), WS-12 (2 μL), and VBJ-103 (2 μL).

### NMR Data analysis

All 1D ^19^F NMR spectra were processed using Bruker TopSpin 4.1 and MestReNova 14.2 software (Mestrelab Research). All 2D ^1^H-^15^N TROSY-HSQC were processed using NMRpipe.^93^ ^1^H-^15^N TROSY-HSQC-based binding data were analyzed using principal component analysis with TrendNMR 1.8.^73^ ^1^H-^15^N TROSY-HSQC-based temperature titrations were analyzed by extracting peak intensities using ccpNMR Analysis.^94^ ^19^F NMR based temperature titration was analyzed by deconvoluting peaks using MestReNova to obtain peak intensity. ^19^F NMR based isothermal binding was determined using an in-house 1D principal component analysis (PCA) Matlab 2023b script. All binding and temperature titration were fit using SigmaPlot 12.0. All Binding isotherms were fit to hyperbolic Langmuir binding curves (eq 4 one site saturation; eq 5: one site saturation + nonspecific [N_s_]).

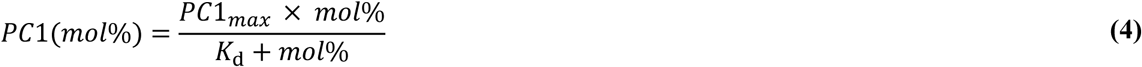

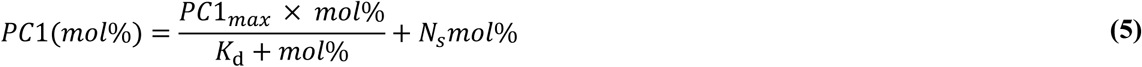

All MDS analyses used Matlab 2023b. The input data for MDS analysis was Bruker TopSpin 4.1 processed ^19^F NMR spectra. The ^19^F NMR difference spectra were generated by subtracting the apo VSLD spectrum from the ligand-bound spectra. Difference spectra were used to create a dissimilarity matrix as input for a classical MDS (cmdscale in Matlab) and computed using an Euclidean distance matrix. The Matlab script is available on the Van Horn Lab GitHub repository: https://github.com/vanhornlab. A two-tailed p-value was calculated using the coefficient of determination (R^2^), number of pair data points (n), and degrees of freedom (n-2) as inputs for an in-house MATLAB script.

### 19F CPMG Relaxation Dispersion

To investigate the dynamics of hTRPM8-VSLD in the apo state and in the presence of saturating concentrations of menthol (agonist) and AMTB (antagonist), triplicate ^19^F relaxation dispersion experiments^84^ were performed at 25°C using samples prepared in the previously stated NMR buffer on a 600 MHz spectrometer. Each experiment was recorded with a relaxation delay (T_cpmg_) of 4 ms and included 15 CPMG frequencies (νCPMG): 250, 500, 750, 1000, 1250, 1500, 1750, 2000, 2250, 2500, 3000, 3500, 4000, 4500, and 5000 Hz. All acquired spectra were processed with a 40 Hz line broadening filter. Spectral deconvolution was performed using TopSpin 4.4.1 with the SOLA algorithm. All spectra were fitted with a Gaussian/Lorentzian ratio set to 0 and Haeberlen–Mehring–Spiess convention. Topspin SOLA CPMG relaxation dispersion data analysis and fit to the Carver-Richards equation were conducted using GUARDD (v.2017.04.25)^95^ to extract chemical exchange rate (*k*_ex_). All error estimates were obtained using a Monte Carlo bootstrap.

### 2D Lineshape analysis

Two-dimensional lineshape fitting of HSQC spectra was performed for titration data of menthol, icilin, menthyl 1d, and VBJ103. Data were collected at 37°C on an 850 MHz spectrometer in previously state NMR buffer. Spectra were processed with an apodization function of 4 Hz in the direct dimension and 8 Hz in the indirect dimension. A total of 9 peaks, selected based on their relative isolation and distribution across the spectra, were used for fitting (Figure S17).The data were fit using a two-state exchange model in TITAN (v1.6)^96^, to extract linewidth (λ) for both direct and indirect dimensions. Change in linewidth between apo and ligand bound state (Δλ = λ_ligand_ − λ_apo_) were calculated for all peaks. All error estimation performed via bootstrap analysis using 100 replicates.

### Temperature-dependent intrinsic tryptophan fluorescence

Fluorescence emission spectra of tryptophan were measured on a QM-4/2005SR Spectrofluorometer (PTI, NJ) using a 295 nm excitation wavelength to minimize the excitation and fluorescence contributions of tyrosine.^97^ The sample temperature was controlled over a range of approximately 6-50°C using a water circulation system, and the temperature was measured directly in the cuvette with a calibrated thermocouple. Light scattering contributions were accounted for by subtracting a blank sample containing 25 mM Na_2_HPO_4_, 0.5 mM EDTA, and 1.099% LPPG from hTRPM8-VLSD sample measurements. The blank signals were relatively small and did not change in magnitude or position when the temperature was changed. Tryptophan fluorescence is sensitive to temperature and the polarity of its environment. A control experiment with free tryptophan in 25 mM Na_2_HPO_4_ and 0.5 mM EDTA showed that as the temperature increased, the fluorescence intensity decreased with no spectral shift. hTRPM8-VLSD samples exhibited a reduced fluorescence intensity and a spectral shift toward higher wavelengths when temperature was increased. The spectral shift was characterized by the change in fluorescence spectra, which were described by calculating the sum of the product of wavelength and normalized fluorescence at that wavelength (eq 6).

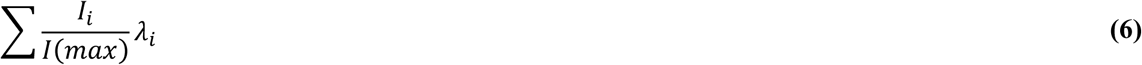

Where λ_i_ is the wavelength, *I*_i_ is the fluorescence intensity at that wavelength, and *I*(*max*) is the intensity at the peak. The fluorescence intensity was normalized using *I*(*max*) to ensure temperature independence in the analysis by accounting for changes in fluorescence efficiency that occur with temperature variations.

As the temperature increased, the fluorescence spectra of the hTRPM8-VLSD shifted toward longer wavelengths, as indicated by an increase in the sum. Temperature titration data were fitted to the eq 3.

## ASSOCIATED CONTENT

The Supporting Information is available free of charge. Supporting information includes structural analysis, chemical structures, supporting functional assay, purification, intrinsic tryptophan fluorescence, NMR, chemometrics analysis, and references.

## AUTHOR INFORMATION

### Corresponding Author

**Wade D. Van Horn** – *School of Molecular Sciences, The Biodesign Institute Center for Personalized Diagnostics, Arizona State University, Tempe, AZ 85281, United States,*

### Present addresses

**Mubark D. Mebrat** – *School of Molecular Sciences, The Biodesign Institute Center for Personalized Diagnostics, Arizona State University, Tempe, AZ 85281, United States,*

**Dustin D. Luu** – *School of Molecular Sciences, The Biodesign Institute Center for Personalized Diagnostics, Arizona State University, Tempe, AZ 85281, United States,*

**Jacob K. Hilton** – *Membrane Transport Biophysics Section, National Institute of Neurological Disorders and Stroke, Bethesda, MD, United States,*

**Minjoo Kim** – *Department of Biomedical Engineering, New York University Tandon School of Engineering, New York, NY, United States,*

**Kaitlyn Parrott** – *School of Molecular Sciences, Arizona State University, Tempe, AZ 85281, United States; The Biodesign Institute Center for Single Molecule Biophysics, Arizona State University, Tempe, AZ, 85287, United States,*

**Brian R. Cherry** – *The Magnetic Resonance Research Center, Arizona State University, Tempe, AZ, 85287, United States,*

**Marcia Levitus** – *School of Molecular Sciences, Arizona State University, Tempe, AZ 85281, United States; The Biodesign Institute Center for Single Molecule Biophysics, Arizona State University, Tempe, AZ, 85287, United State,*

1. **V. Blair Journigan** – *Department of Pharmaceutical Sciences, Department of Chemistry, Computational Chemical Genomics Screening Center, University of Pittsburgh, Pittsburgh, PA 15260, United States,*

## ACKNOWLEDGMENT

Stably expressing human TRPM8 HEK-293T cells was a generous gift from Professor Thomas Voets.

---

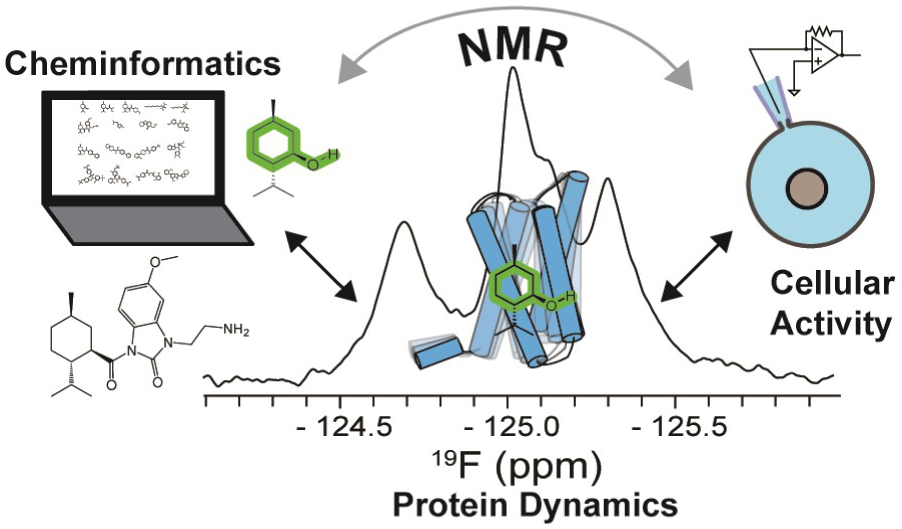
Table of Content Graphics.

## REFERENCES

(1) Luu, D. D.; Ramesh, N.; Kazan, I. C.; Shah, K. H.; Lahiri, G.; Mana, M. D.; Ozkan, S. B.; Van Horn, W. D. Evidence That the Cold– and Menthol-Sensing Functions of the Human TRPM8 Channel Evolved Separately. Sci Adv 2024, 10 (25), 9228. 10.1126/sciadv.adm9228.

(2) Bautista, D. M.; Siemens, J.; Glazer, J. M.; Tsuruda, P. R.; Basbaum, A. I.; Stucky, C. L.; Jordt, S. E.; Julius, D. The Menthol Receptor TRPM8 Is the Principal Detector of Environmental Cold. Nature 2007, 448 (7150), 204–208. 10.1038/nature05910.

(3) Peier, A. M.; Moqrich, A.; Hergarden, A. C.; Reeve, A. J.; Andersson, D. A.; Story, G. M.; Earley, T. J.; Dragoni, I.; McIntyre, P.; Bevan, S.; Patapoutian, A. A TRP Channel That Senses Cold Stimuli and Menthol. Cell 2002, 108 (5), 705–715. 10.1016/S0092-8674(02)00652-9.

(4) Knowlton, W. M.; Daniels, R. L.; Palkar, R.; McCoy, D. D.; McKemy, D. D. Pharmacological Blockade of TRPM8 Ion Channels Alters Cold and Cold Pain Responses in Mice. PLoS One 2011, 6 (9), e25894. 10.1371/journal.pone.0025894.

(5) Chasman, D. I.; Schürks, M.; Anttila, V.; De Vries, B.; Schminke, U.; Launer, L. J.; Terwindt, G. M.; Van Den Maagdenberg, A. M. J. M.; Fendrich, K.; Völzke, H.; Ernst, F.; Griffiths, L. R.; Buring, J. E.; Kallela, M.; Freilinger, T.; Kubisch, C.; Ridker, P. M.; Palotie, A.; Ferrari, M. D.; Hoffmann, W.; Zee, R. Y. L.; Kurth, T. Genome-Wide Association Study Reveals Three Susceptibility Loci for Common Migraine in the General Population. Nat Genet 2011, 43 (7), 695–698. 10.1038/ng.856.

(6) Yee, N. Roles of TRPM8 Ion Channels in Cancer: Proliferation, Survival, and Invasion. Cancers (Basel) 2015, 7 (4), 2134–2146. 10.3390/cancers7040882.

(7) Kaur, J.; Singh, D. P.; Kumar, V.; Kaur, S.; Bhunia, R. K.; Kondepudi, K. K.; Kuhad, A.; Bishnoi, M. Transient Receptor Potential (TRP) Based Polypharmacological Combination Stimulates Energy Expending Phenotype to Reverse HFD-Induced Obesity in Mice. Life Sci 2023, 324, 121704. 10.1016/j.lfs.2023.121704.

(8) Burgos-Vega, C. C.; Ahn, D. D.-U.; Bischoff, C.; Wang, W.; Horne, D.; Wang, J.; Gavva, N.; Dussor, G. Meningeal Transient Receptor Potential Channel M8 Activation Causes Cutaneous Facial and Hindpaw Allodynia in a Preclinical Rodent Model of Headache. Cephalalgia 2016, 36 (2), 185–193. 10.1177/0333102415584313.

(9) Winchester, W. J.; Gore, K.; Glatt, S.; Petit, W.; Gardiner, J. C.; Conlon, K.; Postlethwaite, M.; Saintot, P.-P.; Roberts, S.; Gosset, J. R.; Matsuura, T.; Andrews, M. D.; Glossop, P. A.; Palmer, M. J.; Clear, N.; Collins, S.; Beaumont, K.; Reynolds, D. S. Inhibition of TRPM8 Channels Reduces Pain in the Cold Pressor Test in Humans. Journal of Pharmacology and Experimental Therapeutics 2014, 351 (2), 259–269. 10.1124/jpet.114.216010.

(10) Knowlton, W. M.; Palkar, R.; Lippoldt, E. K.; McCoy, D. D.; Baluch, F.; Chen, J.; McKemy, D. D. A Sensory-Labeled Line for Cold: TRPM8-Expressing Sensory Neurons Define the Cellular Basis for Cold, Cold Pain, and Cooling-Mediated Analgesia. Journal of Neuroscience 2013, 33 (7), 2837–2848. 10.1523/JNEUROSCI.1943-12.2013.

(11) Liu, B.; Fan, L.; Balakrishna, S.; Sui, A.; Morris, J. B.; Jordt, S. E. TRPM8 Is the Principal Mediator of Menthol-Induced Analgesia of Acute and Inflammatory Pain. Pain 2013, 154 (10), 2169–2177. 10.1016/j.pain.2013.06.043.

(12) Di Donato, M.; Ostacolo, C.; Giovannelli, P.; Di Sarno, V.; Monterrey, I. M. G.; Campiglia, P.; Migliaccio, A.; Bertamino, A.; Castoria, G. Therapeutic Potential of TRPM8 Antagonists in Prostate Cancer. Sci Rep 2021, 11 (1), 1–16. 10.1038/s41598-021-02675-4.

(13) Fakih, D.; Baudouin, C.; Réaux-Le Goazigo, A.; Mélik Parsadaniantz, S. TRPM8: A Therapeutic Target for Neuroinflammatory Symptoms Induced by Severe Dry Eye Disease. Int J Mol Sci 2020, 21 (22), 8756. 10.3390/ijms21228756.

(14) Weyer, A.; Lehto, S. Development of TRPM8 Antagonists to Treat Chronic Pain and Migraine. Pharmaceuticals 2017, 10 (2), 37. 10.3390/ph10020037.

(15) Izquierdo, C.; Martín-Martínez, M.; Gómez-Monterrey, I.; González-Muñiz, R. TRPM8 Channels: Advances in Structural Studies and Pharmacological Modulation. Int J Mol Sci 2021, 22 (16), 8502. 10.3390/ijms22168502.

(16) González-Muñiz, R.; Bonache, M. A.; Martín-Escura, C.; Gómez-Monterrey, I. Recent Progress in TRPM8 Modulation: An Update. Int J Mol Sci 2019, 20 (11), 2618. 10.3390/ijms20112618.

(17) Liu, Y.; Mikrani, R.; He, Y.; Faran Ashraf Baig, M. M.; Abbas, M.; Naveed, M.; Tang, M.; Zhang, Q.; Li, C.; Zhou, X. TRPM8 Channels: A Review of Distribution and Clinical Role. Eur J Pharmacol 2020, 882 (January), 173312. 10.1016/j.ejphar.2020.173312.

(18) Luu, D. D.; Owens, A. M.; Mebrat, M. D.; Van Horn, W. D. A Molecular Perspective on Identifying TRPV1 Thermosensitive Regions and Disentangling Polymodal Activation. Temperature 2023, 10 (1), 67–101. 10.1080/23328940.2021.1983354.

(19) Garami, A.; Shimansky, Y. P.; Rumbus, Z.; Vizin, R. C. L.; Farkas, N.; Hegyi, J.; Szakacs, Z.; Solymar, M.; Csenkey, A.; Chiche, D. A.; Kapil, R.; Kyle, D. J.; Van Horn, W. D.; Hegyi, P.; Romanovsky, A. A. Hyperthermia Induced by Transient Receptor Potential Vanilloid-1 (TRPV1) Antagonists in Human Clinical Trials: Insights from Mathematical Modeling and Meta-Analysis. Pharmacol Ther 2020, 208, 107474. 10.1016/j.pharmthera.2020.107474.

(20) Polderman, K. H.; Noc, M.; Beishuizen, A.; Biermann, H.; Girbes, A. R. J.; Tully, G. W.; Seidman, D.; Albertsson, P. A.; Holmberg, M.; Sterz, F.; Holzer, M. Ultrarapid Induction of Hypothermia Using Continuous Automated Peritoneal Lavage with Ice-Cold Fluids: Final Results of the Cooling for Cardiac Arrest or Acute ST-Elevation Myocardial Infarction Trial. Crit Care Med 2015, 43 (10), 2191–2201. 10.1097/CCM.0000000000001158.

(21) Crompton, E. M.; Lubomirova, I.; Cotlarciuc, I.; Han, T. S.; Sharma, S. D.; Sharma, P. Meta-Analysis of Therapeutic Hypothermia for Traumatic Brain Injury in Adult and Pediatric Patients*. Crit Care Med 2017, 45 (4), 575–583. 10.1097/CCM.0000000000002205.

(22) Luyts, N.; Daniluk, J.; Freitas, A. C. N.; Bazeli, B.; Janssens, A.; Mulier, M.; Everaerts, W.; Voets, T. Inhibition of TRPM8 by the Urinary Tract Analgesic Drug Phenazopyridine. Eur J Pharmacol 2023, 942, 175512. 10.1016/j.ejphar.2023.175512.

(23) Lopez, K. E.; Van Horn, W. D. Targeting TRP Channels: The Emerging Role of Cryogenic Electron Microscopy in Drug Discovery. In TRP Channels as Therapeutic Targets; Elsevier, 2024; pp 35–52. 10.1016/B978-0-443-18653-0.00010-1.

(24) Zhao, C.; Xie, Y.; Xu, L.; Ye, F.; Xu, X.; Yang, W.; Yang, F.; Guo, J. Structures of a Mammalian TRPM8 in Closed State. Nat Commun 2022, 13 (1), 3113. 10.1038/s41467-022-30919-y.

(25) Yin, Y.; Le, S. C.; Hsu, A. L.; Borgnia, M. J.; Yang, H.; Lee, S.-Y. Structural Basis of Cooling Agent and Lipid Sensing by the Cold-Activated TRPM8 Channel. Science (1979) 2019, 363 (6430), eaav9334. 10.1126/science.aav9334.

(26) Yin, Y.; Zhang, F.; Feng, S.; Butay, K. J.; Borgnia, M. J.; Im, W.; Lee, S.-Y. Activation Mechanism of the Mouse Cold-Sensing TRPM8 Channel by Cooling Agonist and PIP _2_. Science (1979) 2022, 378 (6616), eadd1268. 10.1126/science.add1268.

(27) Palchevskyi, S.; Czarnocki-Cieciura, M.; Vistoli, G.; Gervasoni, S.; Nowak, E.; Beccari, A. R.; Nowotny, M.; Talarico, C. Structure of Human TRPM8 Channel. Commun Biol 2023, 6 (1), 1065. 10.1038/s42003-023-05425-6.

(28) Diver, M. M.; Cheng, Y.; Julius, D. Structural Insights into TRPM8 Inhibition and Desensitization. Science (1979) 2019, 365 (6460), 1434–1440. 10.1126/science.aax6672.

(29) Yin, Y.; Wu, M.; Zubcevic, L.; Borschel, W. F.; Lander, G. C.; Lee, S.-Y. Structure of the Cold– and Menthol-Sensing Ion Channel TRPM8. Science (1979) 2018, 359 (6372), 237–241. 10.1126/science.aan4325.

(30) Yin, Y.; Park, C.-G.; Zhang, F.; G. Fedor, J.; Feng, S.; Suo, Y.; Im, W.; Lee, S.-Y. Mechanisms of Sensory Adaptation and Inhibition of the Cold and Menthol Receptor TRPM8. Sci Adv 2024, 10 (31), 2211. 10.1126/sciadv.adp2211.

(31) Kamtekar, S.; Hecht, M. H. The Four-Lielix Bundle: What Determines a Fold? The FASEB Journal 1995, 9 (11), 1013–1022. 10.1096/fasebj.9.11.7649401.

(32) Bandell, M.; Dubin, A. E.; Petrus, M. J.; Orth, A.; Mathur, J.; Sun, W. H.; Patapoutian, A. High-Throughput Random Mutagenesis Screen Reveals TRPM8 Residues Specifically Required for Activation by Menthol. Nat Neurosci 2006, 9 (4), 493–500. 10.1038/nn1665.

(33) Xu, L.; Han, Y.; Chen, X.; Aierken, A.; Wen, H.; Zheng, W.; Wang, H.; Lu, X.; Zhao, Z.; Ma, C.; Liang, P.; Yang, W.; Yang, S.; Yang, F. Molecular Mechanisms Underlying Menthol Binding and Activation of TRPM8 Ion Channel. Nat Commun 2020, 11 (1), 3790. 10.1038/s41467-020-17582-x.

(34) Rath, P.; Hilton, J. K.; Sisco, N. J.; Van Horn, W. D. Implications of Human Transient Receptor Potential Melastatin 8 (TRPM8) Channel Gating from Menthol Binding Studies of the Sensing Domain. Biochemistry 2016, 55 (1), 114–124. 10.1021/acs.biochem.5b00931.

(35) Weikl, T. R.; Paul, F. Conformational Selection in Protein Binding and Function. Protein Science. Blackwell Publishing Ltd November 1, 2014, pp 1508–1518. 10.1002/pro.2539.

(36) Boehr, D. D.; Nussinov, R.; Wright, P. E. The Role of Dynamic Conformational Ensembles in Biomolecular Recognition. Nat Chem Biol 2009, 5 (11), 789–796. 10.1038/nchembio.232.

(37) Vogt, A. D.; Di Cera, E. Conformational Selection Is a Dominant Mechanism of Ligand Binding. Biochemistry 2013, 52 (34), 5723–5729. 10.1021/bi400929b.

(38) Ueda, T.; Imai, S.; Shimada, I. Function-Related Dynamics of GPCRs. Journal of Magnetic Resonance 2022, 336, 107164. 10.1016/j.jmr.2022.107164.

(39) Saccuzzo, E. G.; Mebrat, M. D.; Scelsi, H. F.; Kim, M.; Ma, M. T.; Su, X.; Hill, S. E.; Rheaume, E.; Li, R.; Torres, M. P.; Gumbart, J. C.; Van Horn, W. D.; Lieberman, R. L. Competition between Inside-out Unfolding and Pathogenic Aggregation in an Amyloid-Forming β-Propeller. Nat Commun 2024, 15 (1), 155. 10.1038/s41467-023-44479-2.

(40) Alderson, T. R.; Kay, L. E. NMR Spectroscopy Captures the Essential Role of Dynamics in Regulating Biomolecular Function. Cell 2021, 184 (3), 577–595. 10.1016/j.cell.2020.12.034.

(41) Kleckner, I. R.; Foster, M. P. An Introduction to NMR-Based Approaches for Measuring Protein Dynamics. Biochim Biophys Acta Proteins Proteom 2011, 1814 (8), 942–968. 10.1016/j.bbapap.2010.10.012.

(42) Arthanari, H.; Takeuchi, K.; Dubey, A.; Wagner, G. Emerging Solution NMR Methods to Illuminate the Structural and Dynamic Properties of Proteins. Curr Opin Struct Biol 2019, 58, 294–304. 10.1016/j.sbi.2019.06.005.

(43) Alderson, T. R.; Kay, L. E. Unveiling Invisible Protein States with NMR Spectroscopy. Curr Opin Struct Biol 2020, 60, 39–49. 10.1016/j.sbi.2019.10.008.

(44) Sekhar, A.; Kay, L. E. NMR Paves the Way for Atomic Level Descriptions of Sparsely Populated, Transiently Formed Biomolecular Conformers. Proceedings of the National Academy of Sciences 2013, 110 (32), 12867– 12874. 10.1073/pnas.1305688110.

(45) Palmer, A. G.; Kroenke, C. D.; Patrick Loria, J. Nuclear Magnetic Resonance Methods for Quantifying Microsecond-to-Millisecond Motions in Biological Macromolecules. In Nuclear Magnetic Resonance of Biological Macromolecules – Part B; James, T. L., Dötsch, V., Schmitz, U., Eds.; Methods in Enzymology; Academic Press, 2001; Vol. 339, pp 204–238. 10.1016/S0076-6879(01)39315-1.

(46) Marsh, E. N. G.; Suzuki, Y. Using 19 F NMR to Probe Biological Interactions of Proteins and Peptides. ACS Chem Biol 2014, 9 (6), 1242–1250. 10.1021/cb500111u.

(47) Arntson, K. E.; Pomerantz, W. C. K. Protein-Observed Fluorine NMR: A Bioorthogonal Approach for Small Molecule Discovery. J Med Chem 2016, 59 (11), 5158–5171. 10.1021/acs.jmedchem.5b01447.

(48) Manglik, A.; Kim, T. H.; Masureel, M.; Altenbach, C.; Yang, Z.; Hilger, D.; Lerch, M. T.; Kobilka, T. S.; Thian, F. S.; Hubbell, W. L.; Prosser, R. S.; Kobilka, B. K. Structural Insights into the Dynamic Process of Β2-Adrenergic Receptor Signaling. Cell 2015, 161 (5), 1101–1111. 10.1016/j.cell.2015.04.043.

(49) Ye, L.; Van Eps, N.; Zimmer, M.; Ernst, O. P.; Scott Prosser, R. Activation of the A2A Adenosine G-Protein-Coupled Receptor by Conformational Selection. Nature 2016, 533 (7602), 265–268. 10.1038/nature17668.

(50) Ye, L.; Neale, C.; Sljoka, A.; Lyda, B.; Pichugin, D.; Tsuchimura, N.; Larda, S. T.; Pomès, R.; García, A. E.; Ernst, O. P.; Sunahara, R. K.; Prosser, R. S. Mechanistic Insights into Allosteric Regulation of the A2A Adenosine G Protein-Coupled Receptor by Physiological Cations. Nat Commun 2018, 9 (1), 1372. 10.1038/s41467-018-03314-9.

(51) Zhao, J.; Elgeti, M.; O’Brien, E. S.; Sár, C. P.; EI Daibani, A.; Heng, J.; Sun, X.; White, E.; Che, T.; Hubbell, W. L.; Kobilka, B. K.; Chen, C. Ligand Efficacy Modulates Conformational Dynamics of the µ-Opioid Receptor. Nature 2024, 629 (8011), 474–480. 10.1038/s41586-024-07295-2.

(52) Dixon, A. D.; Inoue, A.; Robson, S. A.; Culhane, K. J.; Trinidad, J. C.; Sivaramakrishnan, S.; Bumbak, F.; Ziarek, J. J. Effect of Ligands and Transducers on the Neurotensin Receptor 1 Conformational Ensemble. J Am Chem Soc 2022, 144 (23), 10241–10250. 10.1021/jacs.2c00828.

(53) Yin, Y.; Lee, S.-Y. Current View of Ligand and Lipid Recognition by the Menthol Receptor TRPM8. Trends Biochem Sci 2020, 45 (9), 806–819. 10.1016/j.tibs.2020.05.008.

(54) Talyzina, I. A.; Nadezhdin, K. D.; Sobolevsky, A. I. Forty Sites of TRP Channel Regulation. Curr Opin Chem Biol 2025, 84, 102550. 10.1016/j.cbpa.2024.102550.

(55) O’Brien, E. S.; Fuglestad, B.; Lessen, H. J.; Stetz, M. A.; Lin, D. W.; Marques, B. S.; Gupta, K.; Fleming, K. G.; Wand, A. J. Membrane Proteins Have Distinct Fast Internal Motion and Residual Conformational Entropy. Angewandte Chemie International Edition 2020, 59 (27), 11108–11114. 10.1002/anie.202003527.

(56) Wankowicz, S. A.; de Oliveira, S. H.; Hogan, D. W.; van den Bedem, H.; Fraser, J. S. Ligand Binding Remodels Protein Side-Chain Conformational Heterogeneity. Elife 2022, 11. 10.7554/eLife.74114.

(57) Zhang, K.; Julius, D.; Cheng, Y. Structural Snapshots of TRPV1 Reveal Mechanism of Polymodal Functionality. Cell 2021, 184 (20), 5138–5150.e12. 10.1016/j.cell.2021.08.012.

(58) Journigan, V. B.; Zaveri, N. T. TRPM8 Ion Channel Ligands for New Therapeutic Applications and as Probes to Study Menthol Pharmacology. Life Sci 2013, 92 (8–9), 425–437. 10.1016/j.lfs.2012.10.032.

(59) Pérez de Vega, M. J.; Gómez-Monterrey, I.; Ferrer-Montiel, A.; González-Muñiz, R. Transient Receptor Potential Melastatin 8 Channel (TRPM8) Modulation: Cool Entryway for Treating Pain and Cancer. J Med Chem 2016, 59 (22), 10006–10029. 10.1021/acs.jmedchem.6b00305.

(60) Backman, T. W. H.; Cao, Y.; Girke, T. ChemMine Tools: An Online Service for Analyzing and Clustering Small Molecules. Nucleic Acids Res 2011, 39 (suppl), W486–W491. 10.1093/nar/gkr320.

(61) Journigan, V. B.; Feng, Z.; Rahman, S.; Wang, Y.; Amin, A. R. M. R.; Heffner, C. E.; Bachtel, N.; Wang, S.; Gonzalez-Rodriguez, S.; Fernández-Carvajal, A.; Fernández-Ballester, G.; Hilton, J. K.; Van Horn, W. D.; Ferrer-Montiel, A.; Xie, X.-Q.; Rahman, T. Structure-Based Design of Novel Biphenyl Amide Antagonists of Human Transient Receptor Potential Cation Channel Subfamily M Member 8 Channels with Potential Implications in the Treatment of Sensory Neuropathies. ACS Chem Neurosci 2020, 11 (3), 268–290. 10.1021/acschemneuro.9b00404.

(62) Gold, M. S.; Pineda-Farias, J. B.; Close, D.; Patel, S.; Johnston, P. A.; Stocker, S. D.; Journigan, V. B. Subcutaneous Administration of a Novel TRPM8 Antagonist Reverses Cold Hypersensitivity While Attenuating the Drop in Core Body Temperature. Br J Pharmacol 2024, 181 (18), 3527–3543. 10.1111/bph.16429.

(63) Sliwoski, G.; Kothiwale, S.; Meiler, J.; Lowe, E. W. Computational Methods in Drug Discovery. Pharmacol Rev 2014, 66 (1), 334–395. 10.1124/pr.112.007336.

(64) Willett, P.; Winterman, V. A Comparison of Some Measures for the Determination of Inter-Molecular Structural Similarity Measures of Inter-Molecular Structural Similarity. Quantitative Structure-Activity Relationships 1986, 5 (1), 18–25. 10.1002/qsar.19860050105.

(65) Chuang, H.; Neuhausser, W. M.; Julius, D. The Super-Cooling Agent Icilin Reveals a Mechanism of Coincidence Detection by a Temperature-Sensitive TRP Channel. Neuron 2004, 43 (6), 859–869. 10.1016/j.neuron.2004.08.038.

(66) Voets, T.; Owsianik, G.; Janssens, A.; Talavera, K.; Nilius, B. TRPM8 Voltage Sensor Mutants Reveal a Mechanism for Integrating Thermal and Chemical Stimuli. Nat Chem Biol 2007, 3 (3), 174–182. 10.1038/nchembio862.

(67) Andersson, D. A.; Chase, H. W. N.; Bevan, S. TRPM8 Activation by Menthol, Icilin, and Cold Is Differentially Modulated by Intracellular PH. Journal of Neuroscience 2004, 24 (23), 5364–5369. 10.1523/JNEUROSCI.0890-04.2004.

(68) Bödding, M.; Wissenbach, U.; Flockerzi, V. Characterisation of TRPM8 as a Pharmacophore Receptor. Cell Calcium 2007, 42 (6), 618–628. 10.1016/j.ceca.2007.03.005.

(69) McKemy, D. D.; Neuhausser, W. M.; Julius, D. Identification of a Cold Receptor Reveals a General Role for TRP Channels in Thermosensation. Nature 2002, 416 (6876), 52–58. 10.1038/nature719.

(70) El-Arabi, A. M.; Salazar, C. S.; Schmidt, J. J. Ion Channel Drug Potency Assay with an Artificial Bilayer Chip. Lab Chip 2012, 12 (13), 2409–2413. 10.1039/c2lc40087a.

(71) Sherkheli, M. A.; Vogt-Eisele, A. K.; Bura, D.; Beltrán Márques, L. R.; Gisselmann, G.; Hatt, H. Characterization Of Selective TRPM8 Ligands And Their Structure Activity Response (S.A.R) Relationship. Journal of Pharmacy & Pharmaceutical Sciences 2010, 13 (2), 242. 10.18433/J3N88N.

(72) Kim, M.; Sisco, N. J.; Hilton, J. K.; Montano, C. M.; Castro, M. A.; Cherry, B. R.; Levitus, M.; Van Horn, W. D. Evidence That the TRPV1 S1-S4 Membrane Domain Contributes to Thermosensing. Nat Commun 2020, 11 (1), 4169. 10.1038/s41467-020-18026-2.

(73) Xu, J.; Van Doren, S. R. Affinities and Comparisons of Enzyme States by Principal Component Analysis of NMR Spectra, Automated Using TREND Software. In Methods in Enzymology; Academic Press Inc., 2018; Vol. 607, pp 217–240. 10.1016/bs.mie.2018.05.016.

(74) Clapham, D. E.; Miller, C. A Thermodynamic Framework for Understanding Temperature Sensing by Transient Receptor Potential (TRP) Channels. Proceedings of the National Academy of Sciences 2011, 108 (49), 19492– 19497. 10.1073/pnas.1117485108.

(75) Mishra, N. K.; Urick, A. K.; Ember, S. W. J.; Schönbrunn, E.; Pomerantz, W. C. Fluorinated Aromatic Amino Acids Are Sensitive 19F NMR Probes for Bromodomain-Ligand Interactions. ACS Chem Biol 2014, 9 (12), 2755– 2760. 10.1021/cb5007344.

(76) Picard, L.-P.; Prosser, R. S. Advances in the Study of GPCRs by 19F NMR. Curr Opin Struct Biol 2021, 69, 169–176. 10.1016/j.sbi.2021.05.001.

(77) Danielson, M. A.; Falke, J. J. Use of 19F NMR to Probe Protein Structure and Conformational Changes. Annu Rev Biophys Biomol Struct 1996, 25, 163–195. 10.1146/annurev.bb.25.060196.001115.

(78) Journigan, V. B.; Alarcón-Alarcón, D.; Feng, Z.; Wang, Y.; Liang, T.; Dawley, D. C.; Amin, A. R. M. R.; Montano, C.; Van Horn, W. D.; Xie, X. Q.; Ferrer-Montiel, A.; Fernández-Carvajal, A. Structural and in Vitro Functional Characterization of a Menthyl TRPM8 Antagonist Indicates Species-Dependent Regulation. ACS Med Chem Lett 2021, 12 (5), 758–767. 10.1021/acsmedchemlett.1c00001.

(79) Duncan, D.; Stewart, F.; Frohlich, M.; Urdal, D. Preclinical Evaluation of the Trpm8 Ion Channel Agonist D-3263 for Benign Prostatic Hyperplasia. Journal of Urology 2009, 181 (4S), 503–503. 10.1016/s0022-5347(09)61422-1.

(80) Ayotte, Y.; Woo, S.; LaPlante, S. R. Practical Considerations and Guidelines for Spectral Referencing for Fluorine NMR Ligand Screening. ACS Omega 2022, 7 (15), 13155–13163. 10.1021/acsomega.2c00613.

(81) Mishra, N. K.; Urick, A. K.; Ember, S. W. J.; Schönbrunn, E.; Pomerantz, W. C. Fluorinated Aromatic Amino Acids Are Sensitive 19 F NMR Probes for Bromodomain-Ligand Interactions. ACS Chem Biol 2014, 9 (12), 2755– 2760. 10.1021/cb5007344.

(82) Li, G. C.; Castro, M. A.; Ukwaththage, T.; Sanders, C. R. Optimizing NMR Fragment-Based Drug Screening for Membrane Protein Targets. J Struct Biol X 2024, 9, 100100. 10.1016/j.yjsbx.2024.100100.

(83) Wallerstein, J.; Akke, M. Minute Additions of DMSO Affect Protein Dynamics Measurements by NMR Relaxation Experiments through Significant Changes in Solvent Viscosity. ChemPhysChem 2019, 20 (2), 326–332. 10.1002/cphc.201800626.

(84) Overbeck, J. H.; Kremer, W.; Sprangers, R. A Suite of 19F Based Relaxation Dispersion Experiments to Assess Biomolecular Motions. J Biomol NMR 2020, 74 (12), 753–766. 10.1007/s10858-020-00348-4.

(85) Waudby, C. A.; Ouvry, M.; Davis, B.; Christodoulou, J. Two-Dimensional NMR Lineshape Analysis of Single, Multiple, Zero and Double Quantum Correlation Experiments. J Biomol NMR 2020, 74 (1), 95–109. 10.1007/s10858-019-00297-7.

(86) Waudby, C. A.; Alfonso, I. An Introduction to One– and Two-Dimensional Lineshape Analysis of Chemically Exchanging Systems. J Magn Reson Open 2023, 16–17, 100102. 10.1016/j.jmro.2023.100102.

(87) Brath, U.; Akke, M. Differential Responses of the Backbone and Side-Chain Conformational Dynamics in FKBP12 upon Binding the Transition-State Analog FK506: Implications for Transition-State Stabilization and Target Protein Recognition. J Mol Biol 2009, 387 (1), 233–244. 10.1016/j.jmb.2009.01.047.

(88) Frederick, K. K.; Marlow, M. S.; Valentine, K. G.; Wand, A. J. Conformational Entropy in Molecular Recognition by Proteins. Nature 2007, 448 (7151), 325–329. 10.1038/nature05959.

(89) Malkia, A.; Pertusa, M.; Fernández-Ballester, G.; Ferrer-Montiel, A.; Viana, F. Differential Role of the Menthol-Binding Residue Y745 in the Antagonism of Thermally Gated TRPM8 Channels. Mol Pain 2009, 5, 1744–8069-5–62. 10.1186/1744-8069-5-62.

(90) Kim, S.; Chen, J.; Cheng, T.; Gindulyte, A.; He, J.; He, S.; Li, Q.; Shoemaker, B. A.; Thiessen, P. A.; Yu, B.; Zaslavsky, L.; Zhang, J.; Bolton, E. E. PubChem 2023 Update. Nucleic Acids Res 2023, 51 (D1), D1373–D1380. 10.1093/nar/gkac956.

(91) Mitchell, J. B. O. Machine Learning Methods in Chemoinformatics. WIREs Computational Molecular Science 2014, 4 (5), 468–481. 10.1002/wcms.1183.

(92) Miles, A. J.; Wallace, B. A. CDtoolX, a Downloadable Software Package for Processing and Analyses of Circular Dichroism Spectroscopic Data. Protein Science 2018, 27 (9), 1717–1722. 10.1002/pro.3474.

(93) Delaglio, F.; Grzesiek, S.; Vuister, G. W.; Zhu, G.; Pfeifer, J.; Bax, A. NMRPipe: A Multidimensional Spectral Processing System Based on UNIX Pipes. J Biomol NMR 1995, 6 (3), 277–293. 10.1007/BF00197809.

(94) Vranken, W. F.; Boucher, W.; Stevens, T. J.; Fogh, R. H.; Pajon, A.; Llinas, M.; Ulrich, E. L.; Markley, J. L.; Ionides, J.; Laue, E. D. The CCPN Data Model for NMR Spectroscopy: Development of a Software Pipeline. *Proteins: Structure*, Function and Genetics 2005, 59 (4), 687–696. 10.1002/prot.20449.

(95) Kleckner, I. R.; Foster, M. P. GUARDD: User-Friendly MATLAB Software for Rigorous Analysis of CPMG RD NMR Data. J Biomol NMR 2012, 52 (1), 11–22. 10.1007/s10858-011-9589-y.

(96) Waudby, C. A.; Ramos, A.; Cabrita, L. D.; Christodoulou, J. Two-Dimensional NMR Lineshape Analysis. Sci Rep 2016, 6 (1), 24826. 10.1038/srep24826.

(97) Royer, C. A. Probing Protein Folding and Conformational Transitions with Fluorescence. Chem Rev 2006, 106 (5), 1769–1784. 10.1021/cr0404390.

