## Supplemental Files for "TRPM8 protein dynamics correlates with ligand structure and cellular function"

### Table of Contents

#### SI figures

Figure S1. Comparison of selected TRPM8 cryo-EM structures

Figure S2. Chemical structures of TRP channel agonists and antagonists used in multidimensional scaling chemometric analysis

Figure S3. Agonist titration of menthol with automated patch clamp

Figure S4. Agonist data from automated patch clamp

Figure S5. Antagonist data from automated patch clamp

Figure S6. Expression test for hTRPM8-VSLD

Figure S7. SDS-PAGE, $^{1}$H-$^{15}$N HSQC spectrum, and CD spectrum show well-folded purified hTRPM8-VSLD

Figure S8. hTRPM8-VSLD thermosensitivity validated using intrinsic tryptophan fluorescence

Figure S9. $^{19}$F NMR spectra of hTRPM8-VSLD with the corresponding tryptophan peak assignments

Figure S10. 1D $^{19}$F NMR replicates 2D $^{1}$H-$^{15}$N TROSY-HSQC binding measurements

Figure S11. hTRPM8-VSLD thermosensitivity validated using $^{19}$F NMR, $^{15}$N-HSQC, and intrinsic tryptophan fluorescence

Figure S12. Menthol binding is coupled to temperature sensitivity

Figure S13. Multidimensional scaling analysis for all chemical structures and ligand-bound hTRPM8 $^{19}$F NMR spectra

Figure S14. DMSO induced 19F NMR hTRPM8-VSLD spectral changes

Figure S15. Round three multidimensional scaling analysis for all chemical structures and ligand-bound hTRPM8 $^{19}$F NMR spectra

Figure S16. $^{19}$F NMR reveals distinct dynamics near the binding site in the presence agonist verses antagonist bound states

Figure S17. 2D $^{1}$H-$^{15}$N HSQC Lineshape analysis shows ligand-dependent backbone dynamics in hTRPM8-VSLD

Figure S18. hTRPM8-VSLD conformational dynamics correlates with function

Figure S19. W798 $^{19}$F NMR peak intensity and linewidth correlate with cellular function

Figure S20. Correlation between functional activity and binding affinity used to estimate $K_d$ values

#### SI Tables

Table S1. Cryo-EM Structures of TRPM8 orthologs

Table S2. Pairwise RMSD between selected TRPM8-VSLD (S1-S4; corresponding the human S733-L853)

Table S3. Pairwise RMSD analysis of all TRPM8 channel structures

Table S4. Expression levels of TRPM8 S1–S4 domain under various conditions

Table S5. Menthol or Menthol+VBJ-103 induced chemical shift change ($\Delta\delta$)

Table S6. $^{1}$H lineshape analysis of $^{1}$H-$^{15}$N HSQC based agonist and antagonist titration

Table S7. $^{15}$N lineshape analysis of $^{1}$H-$^{15}$N HSQC based agonist and antagonist titration

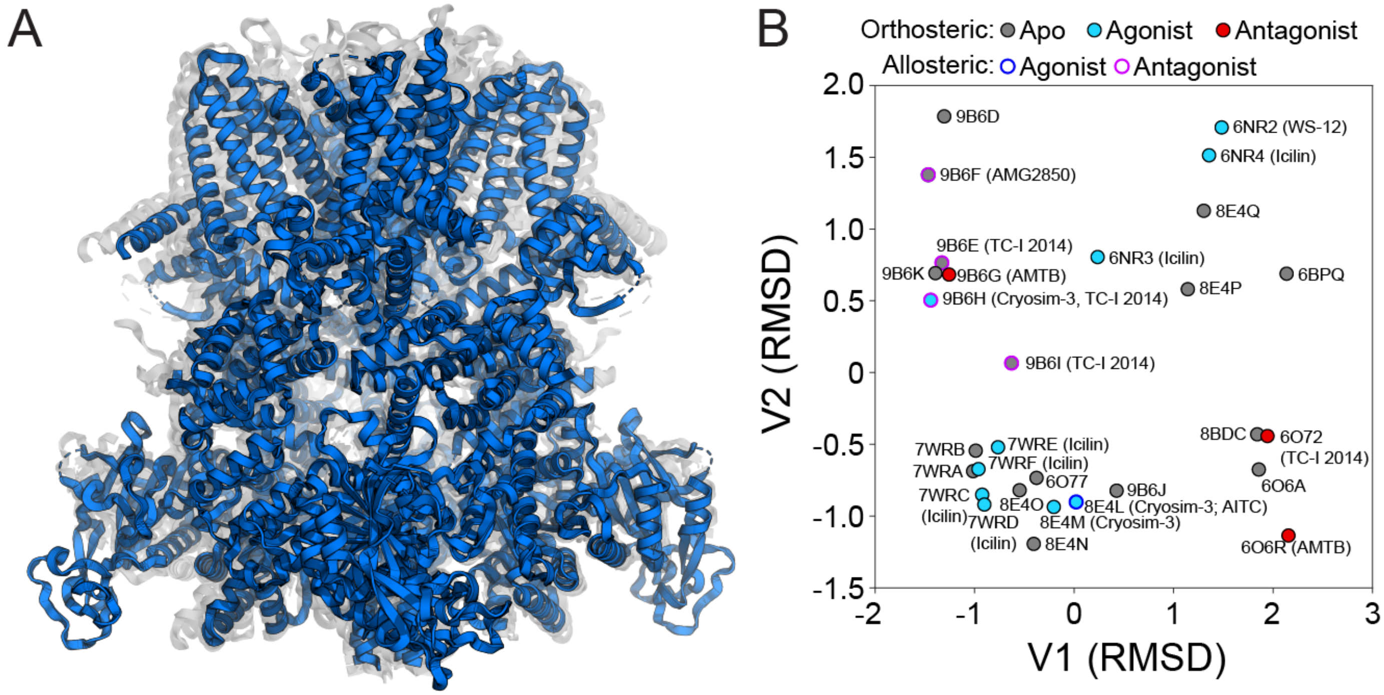

**Figure S1. Comparison of selected TRPM8 cryo-EM structures.** Pymol alignment algorithm CE align of the entire structures was used. (A) In blue is apo human TRPM8 structure (pdb 8BDC) aligned to representative orthosteric ligand-bound TRPM8 structures: cryosim-3 (pdb 8E4L), AMTB (pdb 6O6R), TC-I 2014 (pdb 6O72), WS-12 (pdb 6NR2), and icilin (pdb 7WRD). (B) The RMSD was used to generate a distance matrix for the 29 TRPM8 cryo-EM structures. Each point represents a structure with the orthosteric site in Apo (gray), agonist-bound (blue), or antagonist-bound (red). The outline of the points indicates presences of allosterically bound ligand: agonist (dark blue) or antagonist (purple).

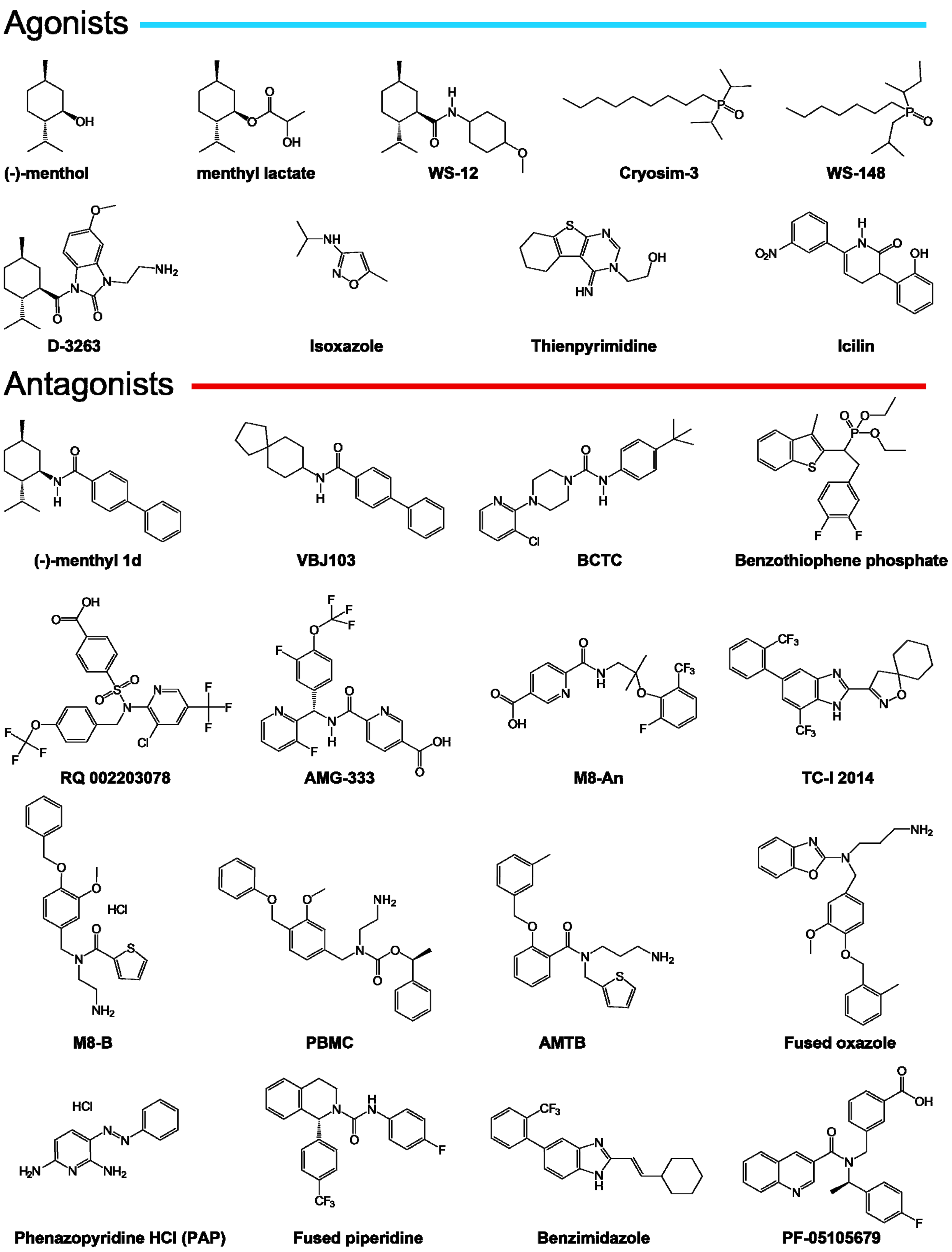

**Figure S2. Chemical structures of TRP channel agonists and antagonists used in multidimensional scaling chemometric analysis.** A library of 25 known TRPM8 regulating agonists and antagonists was identified; including all compounds visualized in TRPM8 structures (WS-12, icilin, AMTB, TC-I 2014, and cryosim-3) and chemotypes beyond menthol derivatives.

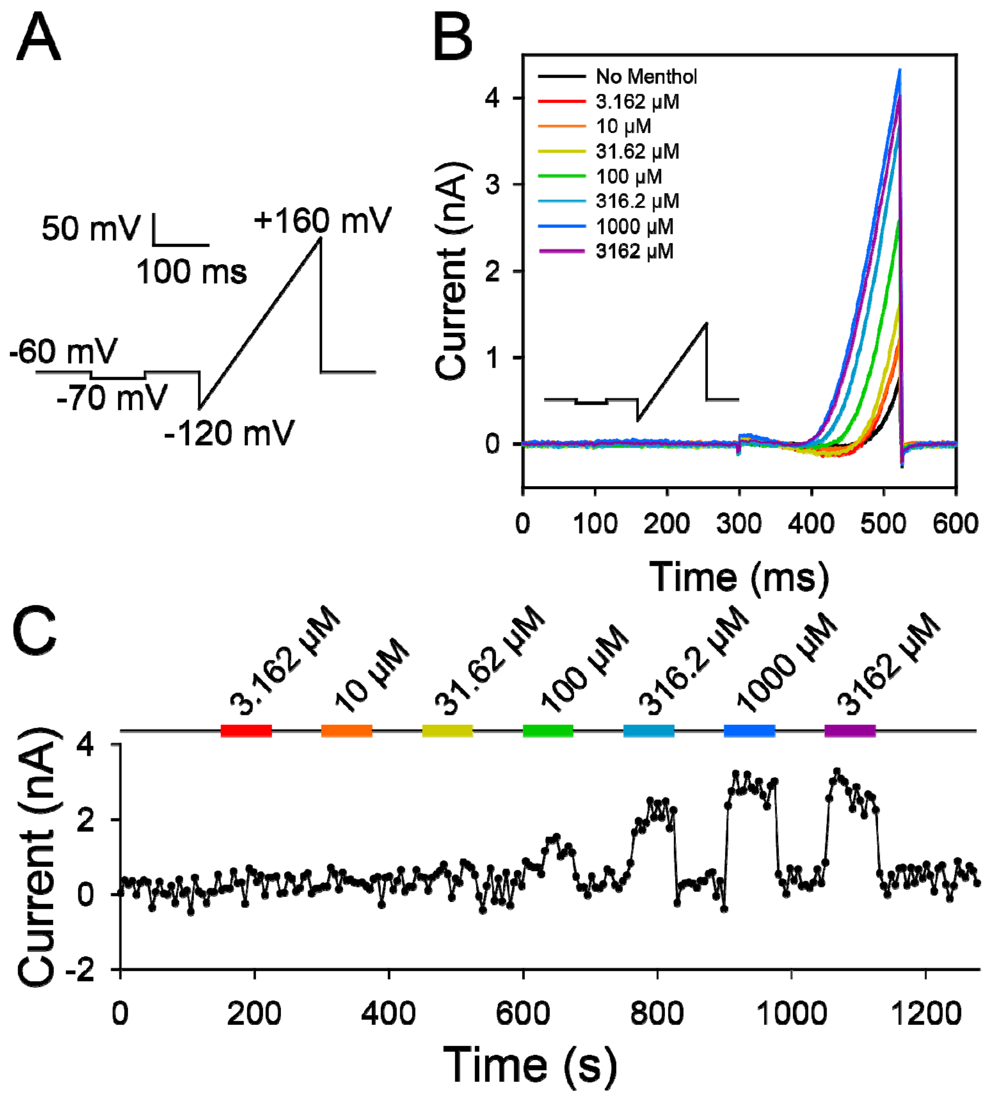

**Figure S3. Agonist titration of menthol with automated patch clamp.** (A) The pulse program used to get the current trace at each point. (B) An overlay of the sweeps at varying concentrations of menthol after 25 s of exposure to each concentration. (C) Representative current trace over time of human TRPM8, stably expressed in HEK203T cells, activated with increasing concentrations of menthol (rainbow) with washout of extracellular solution (black) in between.

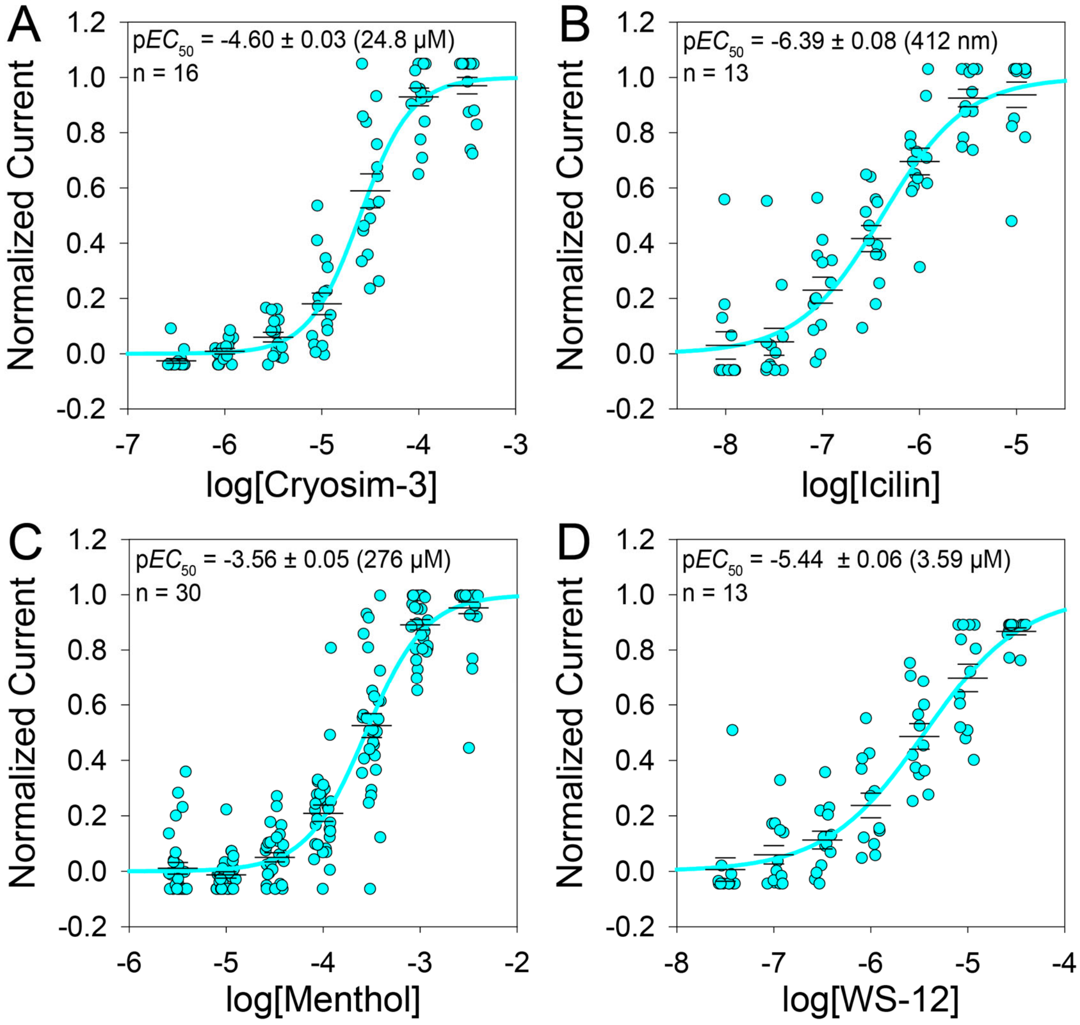

**Figure S4. Agonist data from automated patch clamp.** Jitter and sigmoidal fit of (A) cryosim-3, (B) icilin, (C) menthol, and (D) WS-12 using automated patch clamp with the $EC_{50}$ , respectively. The middle wider bars are the mean and the smaller bar above and below are the standard error of the mean.

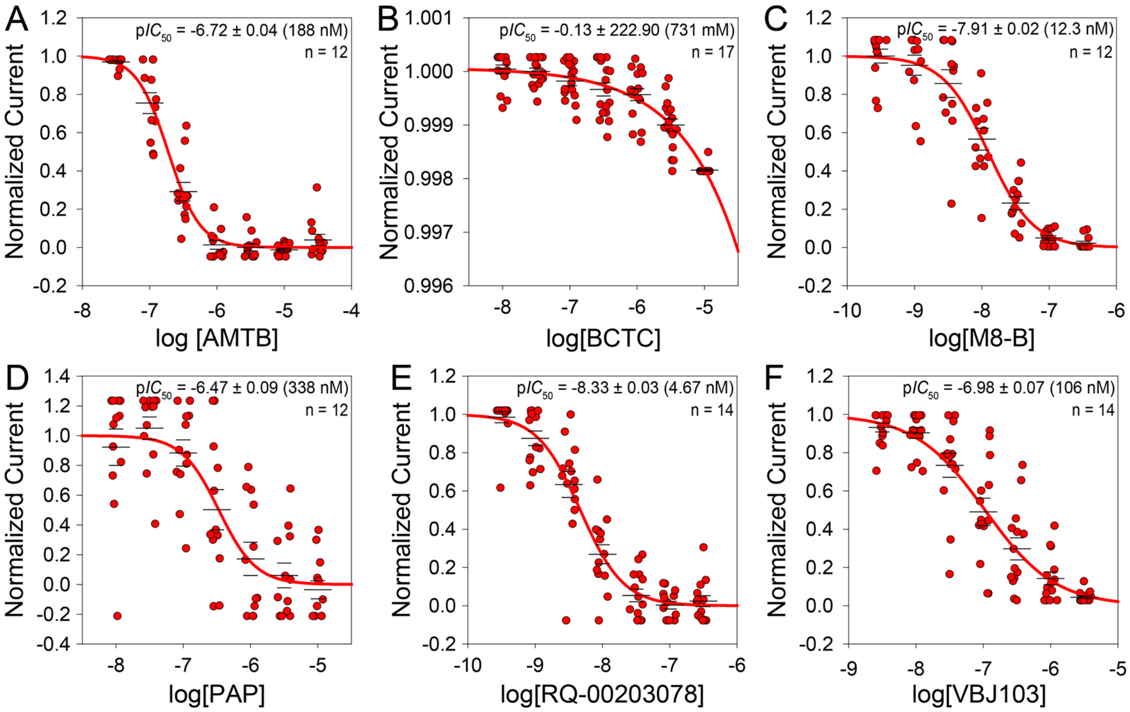

**Figure S5. Antagonist data from automated patch clamp.** Jitter and sigmoidal fit of (A) AMTB, (B) BCTC, (C) M8-B, (D) PAP, (E) RQ-00203078, and (F) VBJ103 using automated patch clamp with the $EC_{50}$ respectively. The middle wider bars are the mean and the smaller bar above and below are the standard error of the mean.

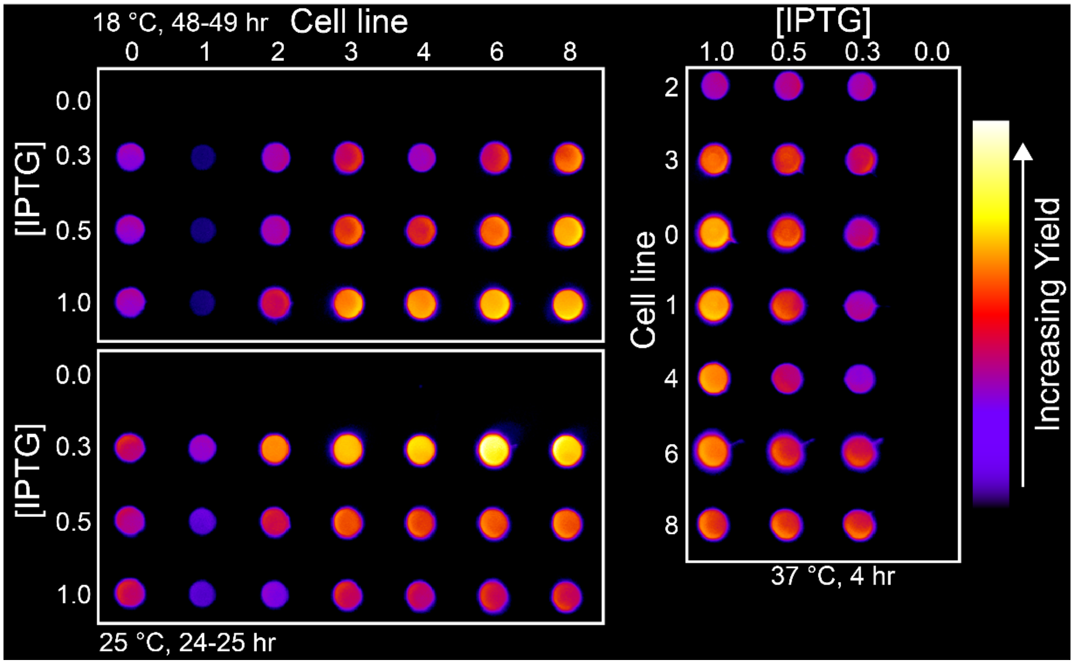

**Figure S6. Expression test for hTRPM8-VSLD**. 63 conditions were tested: three growth temperatures (18 °C, 25 °C, 37 °C), three IPTG concentrations (0.3 mM, 0.5 mM, and 1 mM), and seven cell lines. Cell lines are numbered as: 0, BL21(DE3); 1, BL21(DE3) Star; 2, C41(DE3); 3, Rosetta 2(DE3); 4, C43(DE3) Rosetta2; 6, BL21(DE3) CodonPlus RP; 8 BL21(DE3) LOBSTR. The color gradient from purple to red to yellow represents the lowest (purple) to highest (bright yellow) signal intensity.

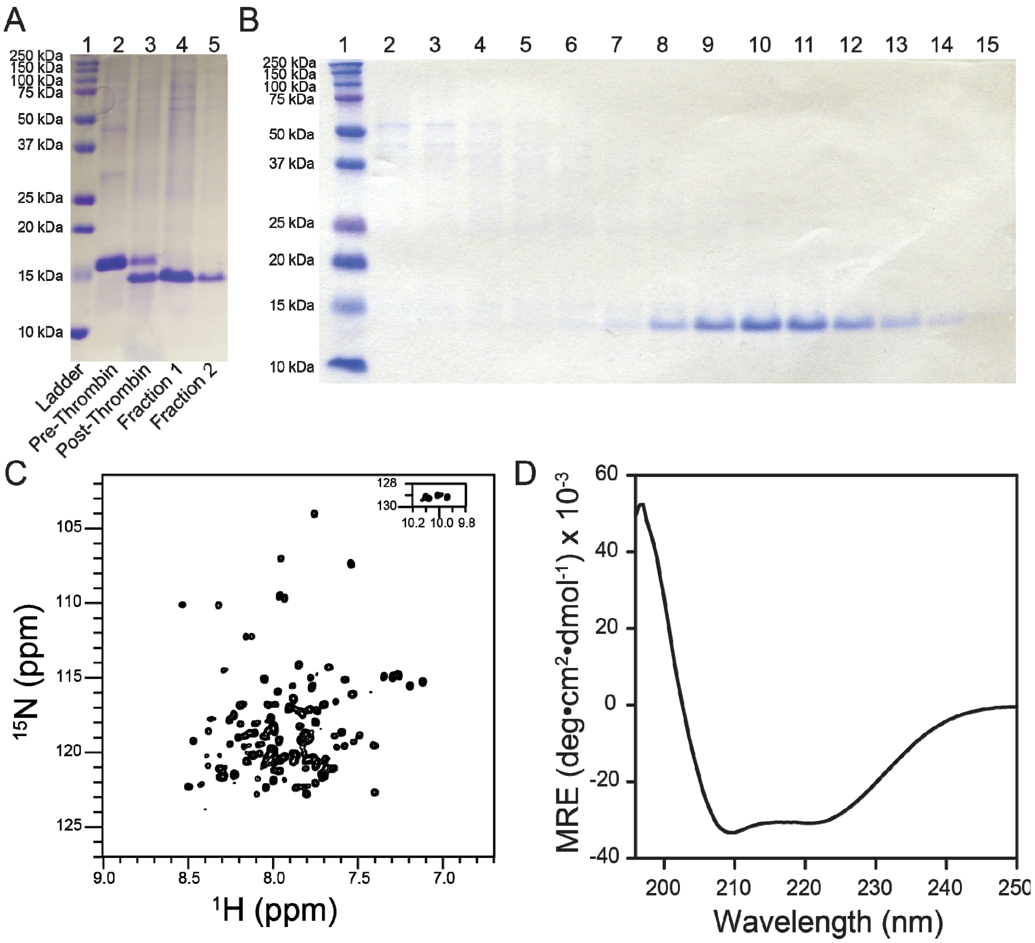

**Figure S7. SDS-PAGE, $^{1}$H-$^{15}$N HSQC spectra,and CD spectrum show well-folded purified hTRPM8-VSLD**. (A) SDS-PAGE of protein ladder (lane 1), pre-thrombin (lane 2), and post-thrombin hTRPM8-VSLD (lanes 3,4, and 5). (B) SDS-PAGE of protein ladder (lane 1), size exclusion fraction (SEC) containing impurities (lane 2-6), and purified hTRPM8-VSLD (lane 7-14). (C) hTRPM8-VSLD $^{1}$H-$^{15}$N TROSY-HSQC spectrum collected at 37 °C. (D) Far-UV circular dichroism spectrum shows purified hTPRM8-VSLD with features minima at 220 and 208 nm and maximums at 195 nm, typical of $\alpha$-helical structure.

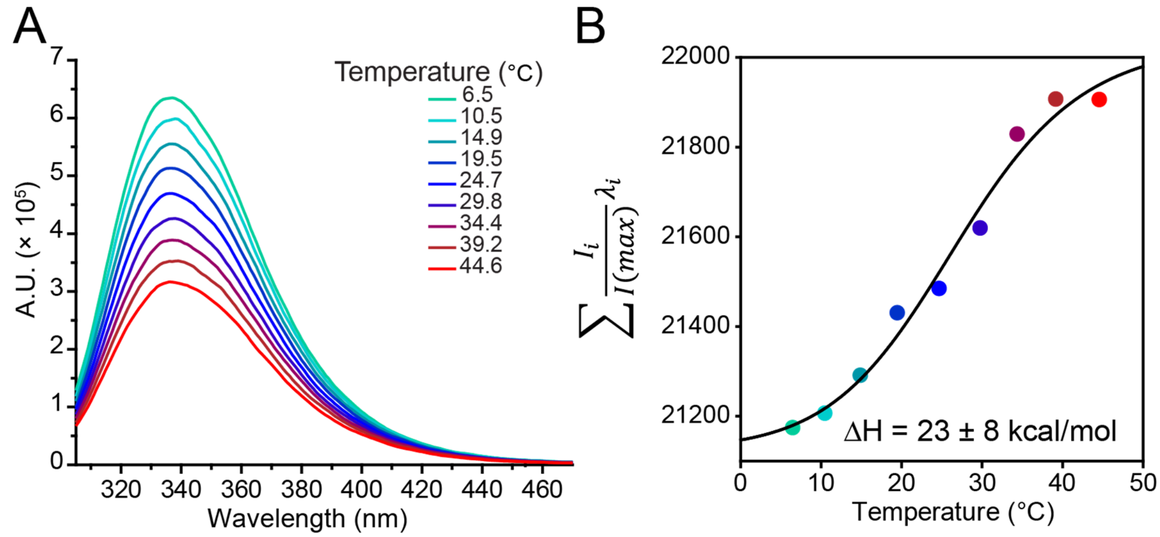

**Figure S8. hTRPM8-VSLD thermosensitivity validated using intrinsic tryptophan fluorescence**. (A) Spectral shift of tryptophan fluorescence as a function of temperature. (B) The spectral shift was quantified using Eq. 5, where $\lambda i$ is the wavelength, and $I_i/(max)$ is the fluorescence at wavelength $\lambda i$ normalized by the intensity at the peak ($I(max)$). The data was fitted to a two-state sigmodal curve. The ΔH contribution was determined to be ΔH = 23 ± 8 kcal/mol.

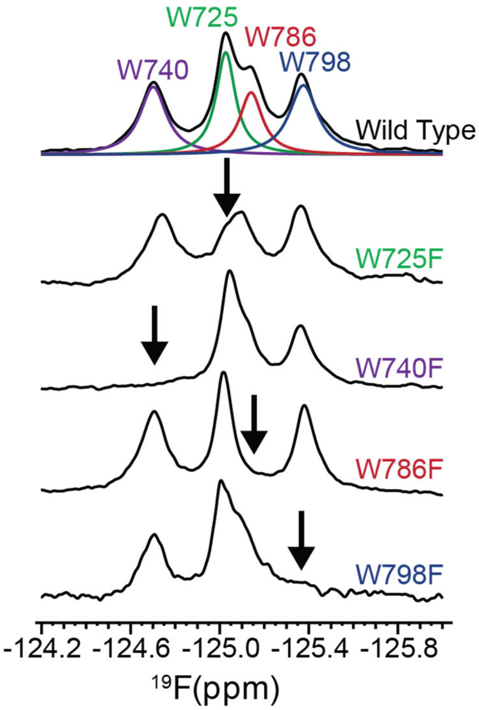

**Figure S9. $^{19}$F NMR spectra of hTRPM8-VSLD with the corresponding tryptophan peak assignments**. Single-point mutation (Trp→Phe) was introduced at each tryptophan residue. $^{19}$F NMR spectra of wild type and four hTRPM8-VSLD mutants with black arrows indicating the mutated tryptophan residue resonance position.

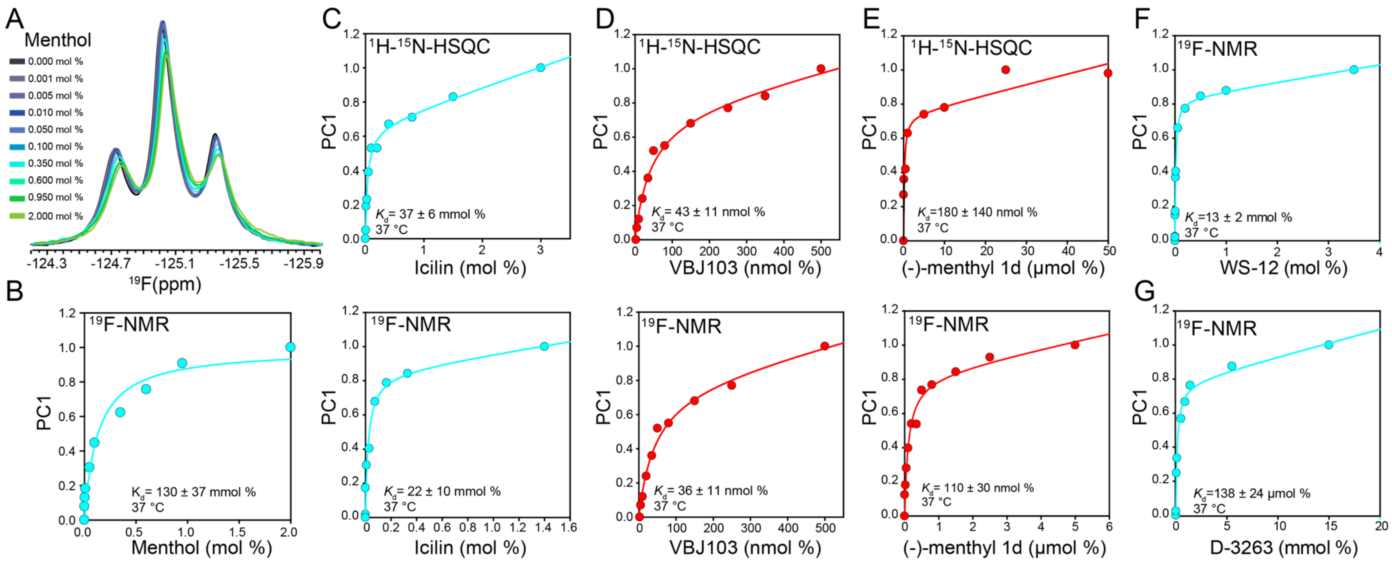

**Figure S10. 1D $^{19}$F NMR replicates 2D $^{1}$H-$^{15}$N TROSY-HSQC binding measurements.** (A) $^{19}$F NMR spectra were collected for each menthol titration point, and spectra were overlaid. Spectra overlay shows a concentration-dependent chemical shift. (F) Principal component analysis (PCA) was conducted on menthol titration $^{19}$F NMR spectral data. PC1 was fitted to a specific non-specific binding curve to determine $K_d$ (110 ± 30 mmol %) consistent with $^{1}$H-$^{15}$N TROSY HSQC data. Titration data were collected using $^{1}$H-$^{15}$N TROSY-HSQC and $^{19}$F NMR for (C) Icilin (HSQC: 37 ± 6 mmol %;$^{19}$F NMR: 22 ± 10 mmol), (D) VBJ103(HSQC:43.5 ± 11 nmol %; $^{19}$F NMR:36 ± 11 nmol %),(E) (-)-menthyl 1d (HSQC: 180 ± 140 nmol %; $^{19}$F NMR:110 ± 30 nmol %), compounds. Binding affinity for (F) WS-12(13± 2 nmol %) and (G) D-3263(138± 24 nmol %) were collected using only $^{19}$F NMR

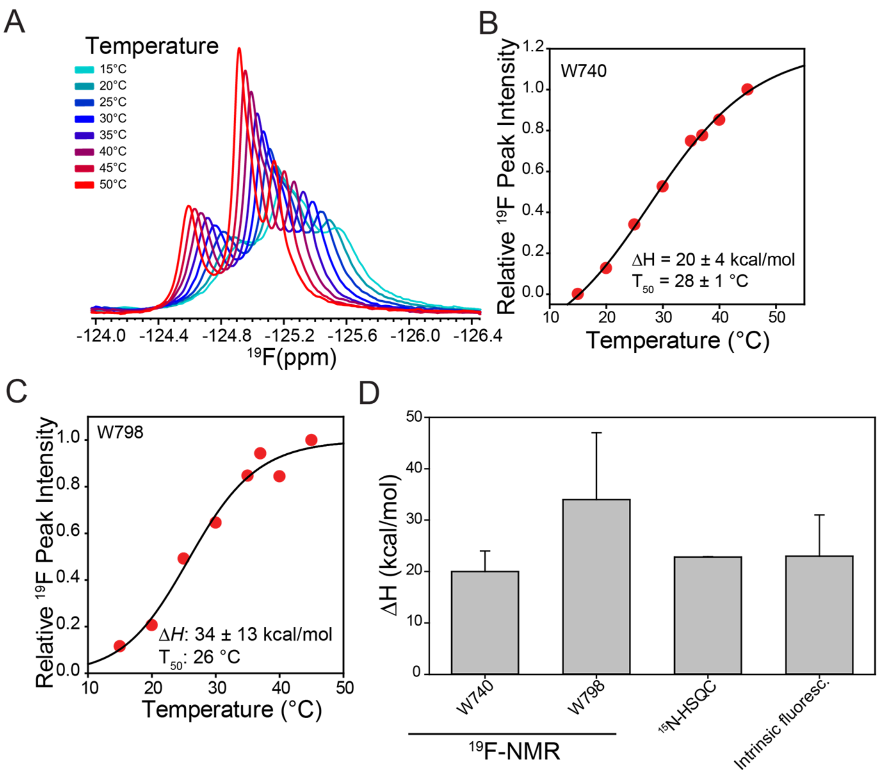

**Figure S11. hTRPM8-VSLD thermosensitivity validated using $^{19}$F NMR, $^{15}$N-HSQC, and intrinsic tryptophan fluorescence.** $^{19}$F NMR spectral were collected from 15 °C to 50 °C with 5 °C increment. (A) $^{19}$F NMR spectral overlaid shows temperature-dependent peak intensity changes. (B) $^{19}$F NMR based temperature-relative peak (W740) intensity response was fitted to a two-state sigmodal curve and obtained $\Delta H$ (20 ± 4 kcal/mol). (C) Change in (W798) intensity as function of temperature was fitted to two-state sigmodal curve to determine ΔH (34 ± 13 kcal/mol). (D) Comparison of $\Delta H$ measurement across $^{19}$F NMR, $^{15}$N-HSQC, and intrinsic tryptophan fluorescence is shown.

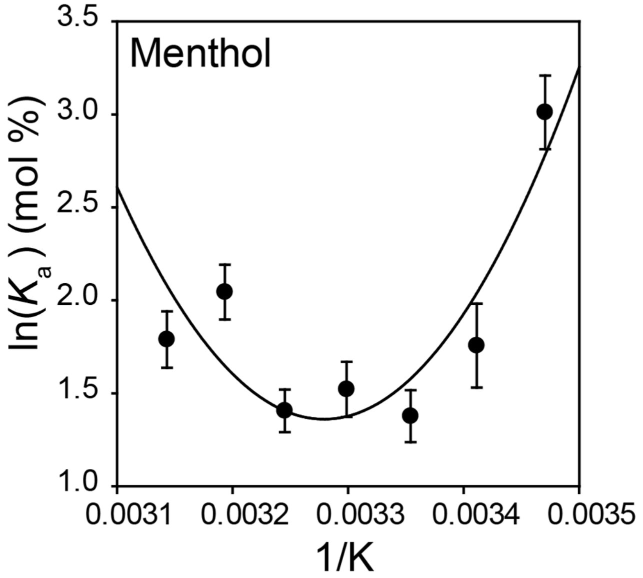

**Figure S12. Menthol binding is coupled to temperature sensitivity.** Van't Hoff plots were construcfor menthol binding. Menthol shows a van't Hoff plot with a significant deviation from linearity, with a calculated $\Delta C_p$ = 2.64 kcal/K.

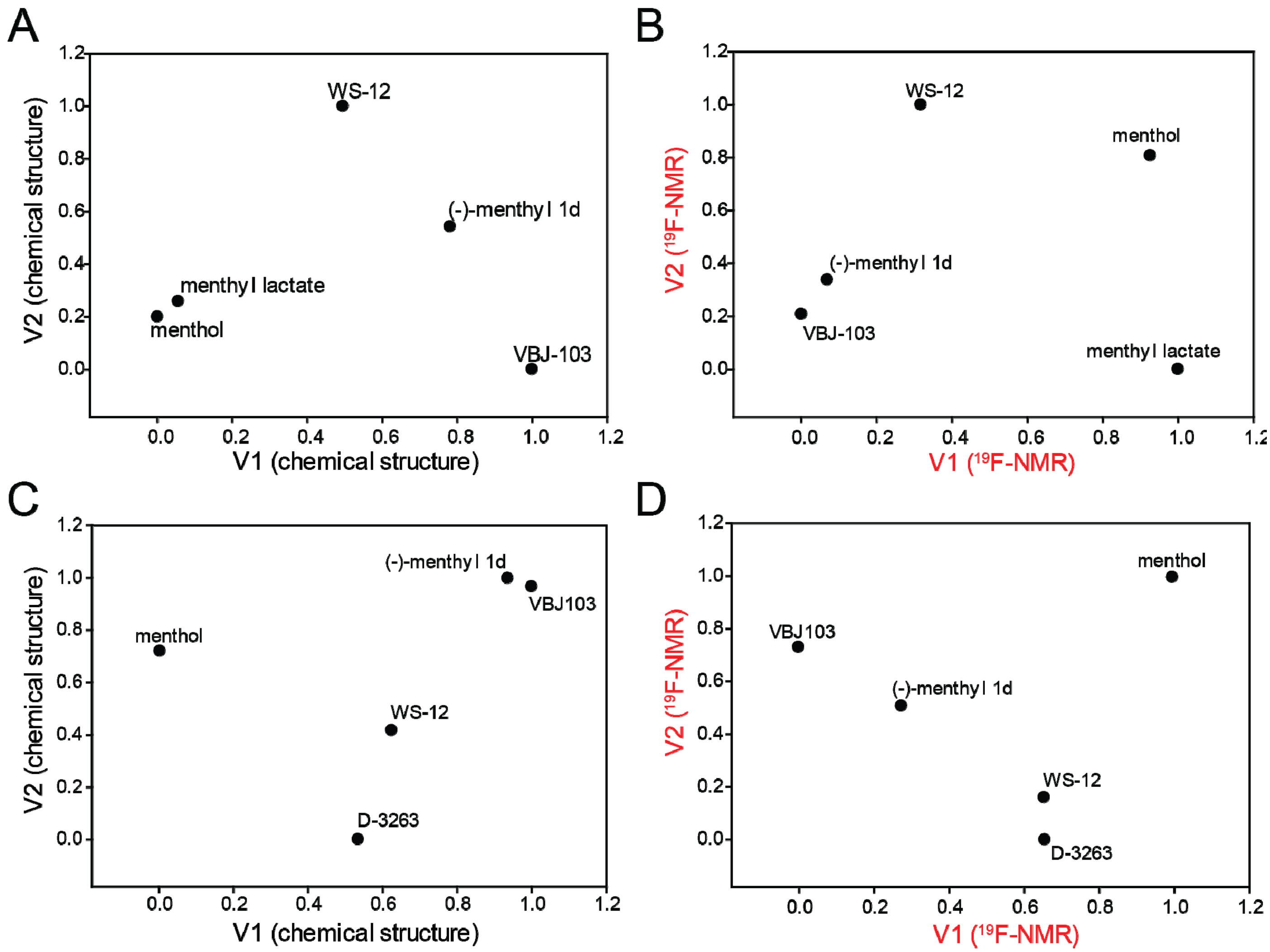

**Figure S13: Multidimensional scaling analysis for all chemical structures and ligand-bound hTRPM8 $^{19}$F NMR spectra**. MDS 2D plots for the first and second rounds of chemical compounds (A and C, respectively) and ligand-bound hTRPM8 $^{19}$F NMR (B and D, respectively).

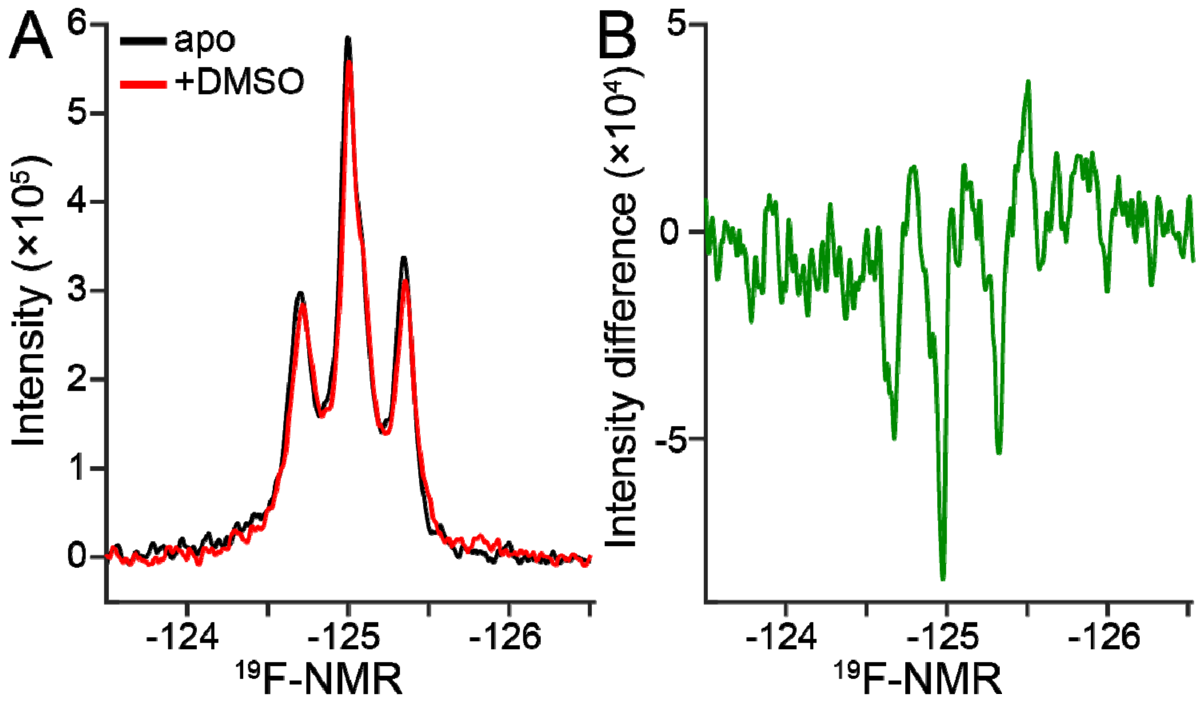

**Figure S14. DMSO induced $^{19}$F NMR hTRPM8-VSLD spectral changes**. (A) $^{19}$F NMR spectral overlay of apo (black) and in the presence of 0.6% DMSO (red).Changes in chemical shift (~Δδ: 0.007-0.016) and intensity (5% to 8% decrease) were observed between apo (black) and in the presence of DMSO (0.6%). (B) $^{19}$F NMR difference spectra was produced by subtracting apo from +DMSO.

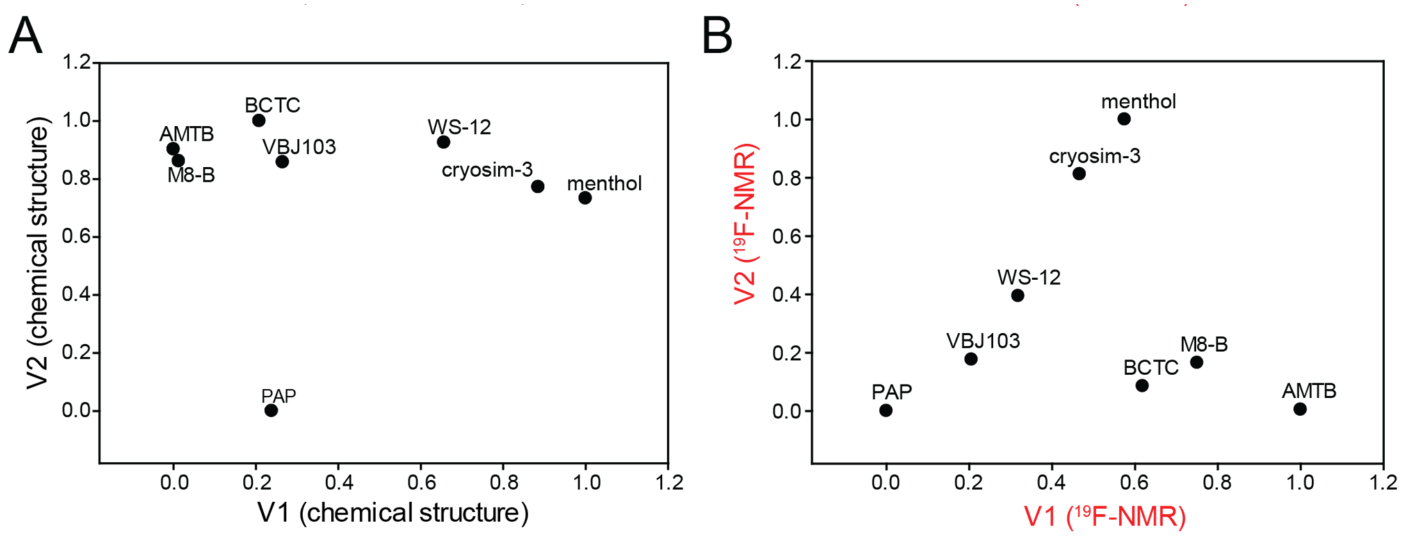

**Figure S15: Round three multidimensional scaling analysis for all chemical structures and ligand-bound hTRPM8 $^{19}$F NMR spectra**. MDS 2D plots for the third rounds of chemical compounds (A) and ligand-bound hTRPM8 $^{19}$F NMR (B).

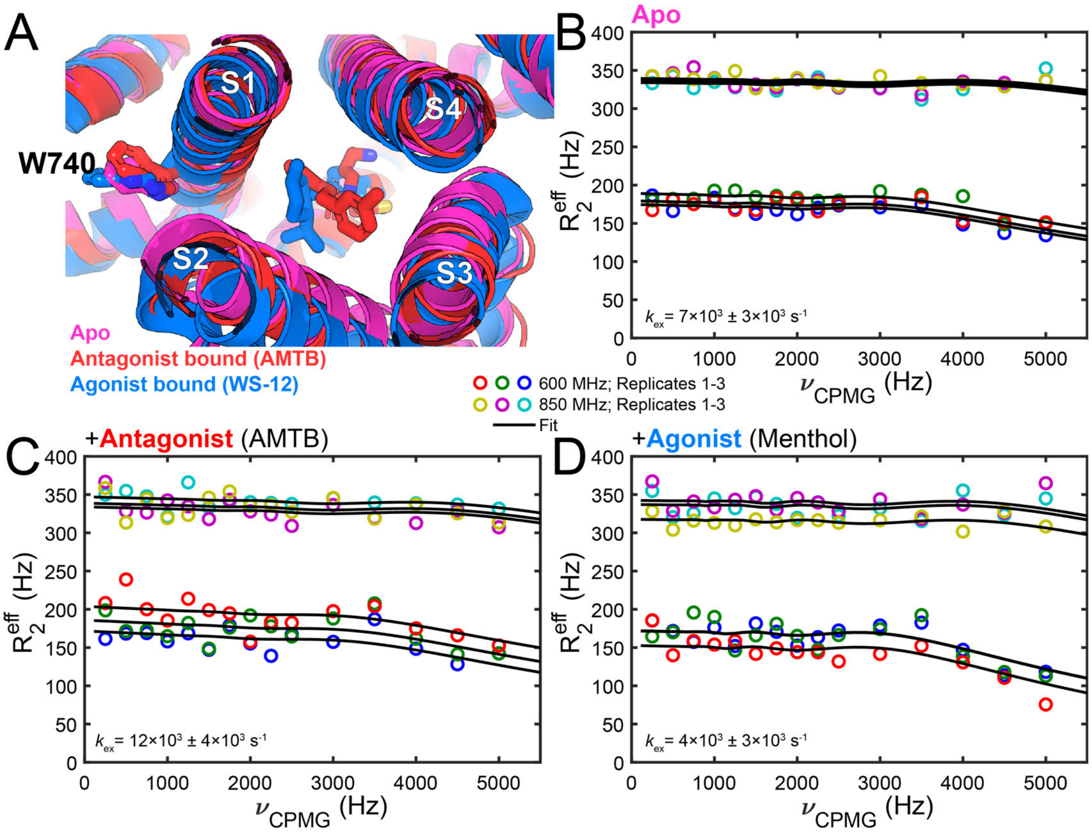

**Figure S16. $^{19}$F NMR reveals distinct dynamics near the binding site in the presence agonist verses antagonist bound states.** (A) Structural alignment of Apo human TRPM8 structure (pdb 8BDC, magenta) with a representative orthosteric ligand-bound TRPM8 structures: antagonist-bound AMTB (pdb 6O6R, blue) and agonist-bound WS-12 (pdb 6NR2, blue). $^{19}$F CPMG relaxation dispersion data for 5-fluorotryptophan-labeled W740 residue of the hTRPM8 VSLD in (B) apo, (C) AMTB-bound (D) menthol-bound states. The unbound (apo) exchange rate ($k_{ex}$, $7\times10^3 \pm 3\times10^3$ s$^{-1}$) increases by ~1.7 fold in the presence of AMTB antagoinist ($k_{ex}$ = $12\times10^3 \pm 4\times10^3$ s$^{-1}$), and the canonical agonist menthol attenuates the apo $k_{ex}$ by ~1.6 fold to $4\times10^3 \pm 3\times10^3$ s$^{-1}$. These data show a ligand-dependent trend in W740 dynamics near the orthosteric binding site.

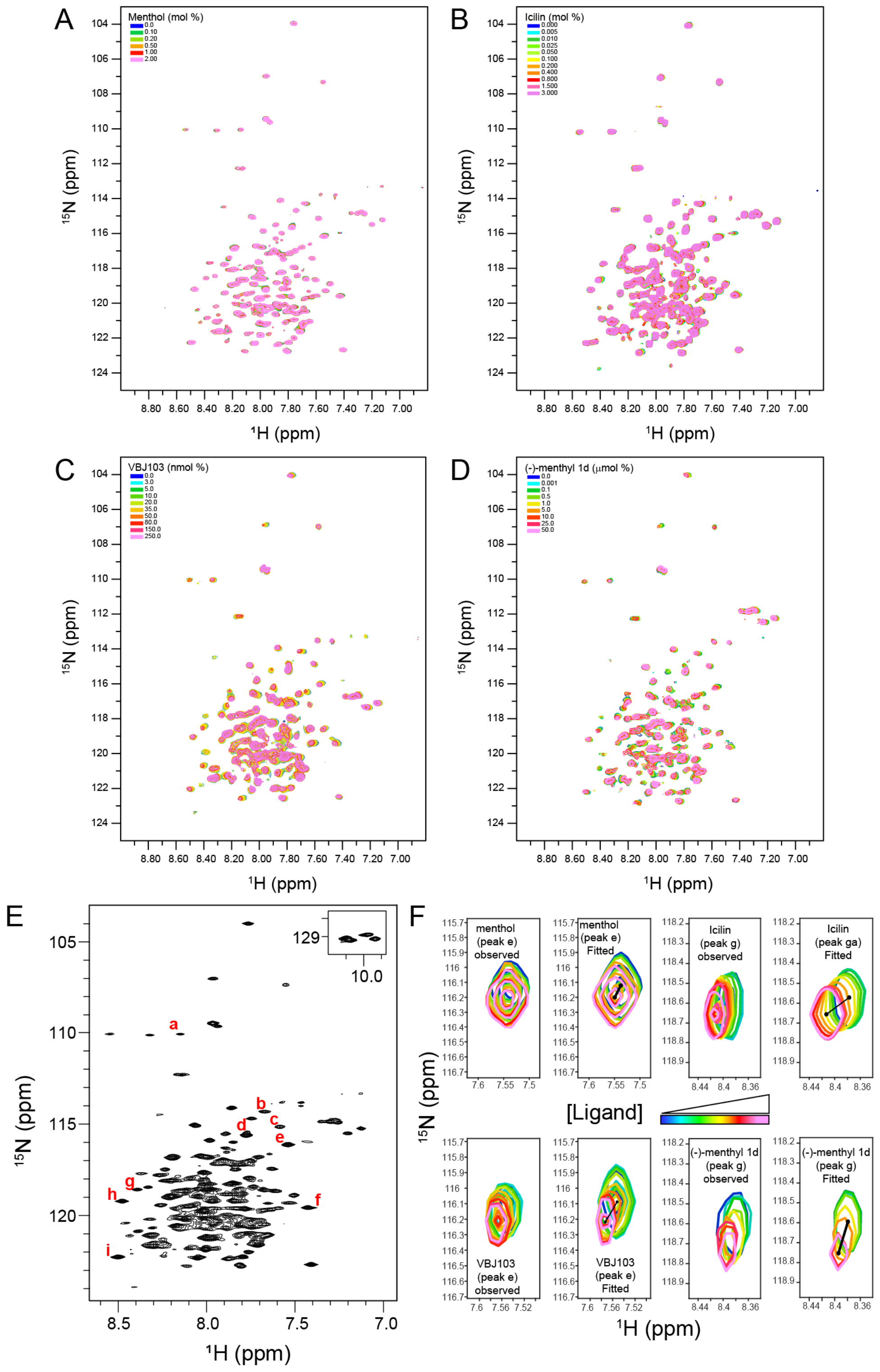

**Figure S17. 2D $^1$H-$^{15}$N HSQC Lineshape analysis shows ligand-dependent backbone dynamics in hTRPM8-VSLD.** (A)$^1$H-$^{15}$N TROSY-HSQC spectra for (A) menthol, (B) icilin, (C) VBJ103, and (D) (-)-menthyl 1d titration data. (E) Representative 2D $^1$H-$^{15}$N HSQC spectrum of hTRPM8-VSLD, highlighting nine well-isolated peaks (labeled a-i) distributed across the spectrum. (F) Examples of observed and fitted resonance from titrations with agonists (menthol and icilin) and antagonists (VBJ103 and (-)-menthyl 1d). The selected peaks exhibit distinct spectral differences between agonist and antagonist binding, indicating differential effects on local backbone dynamics.

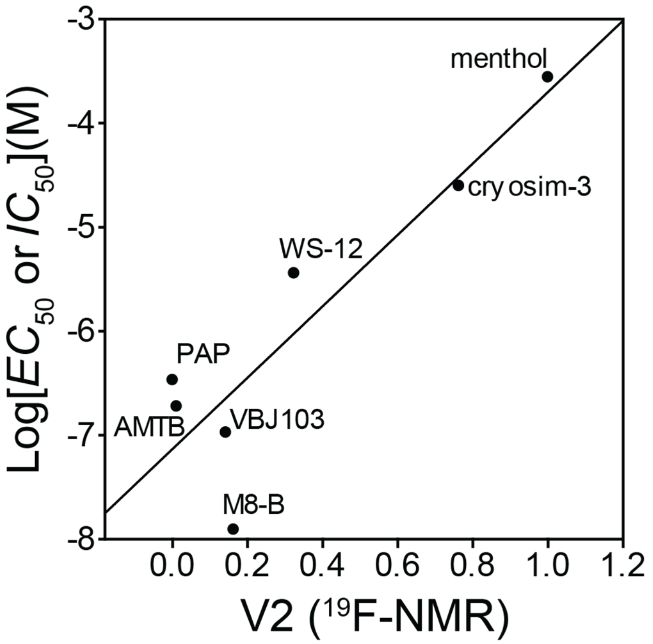

**Figure S18. hTRPM8-VSLD conformational dynamics correlates with function.** Correlation plot between functional data ($EC_{50}$ or $IC_{50}$) and hTRPM8-VSLD conformational dynamics data for menthol, cryosim-3, WS-12, PAP, VBJ103, AMTB, and M8-B with a $R^2$ of 0.791 and p-value of 0.0074.

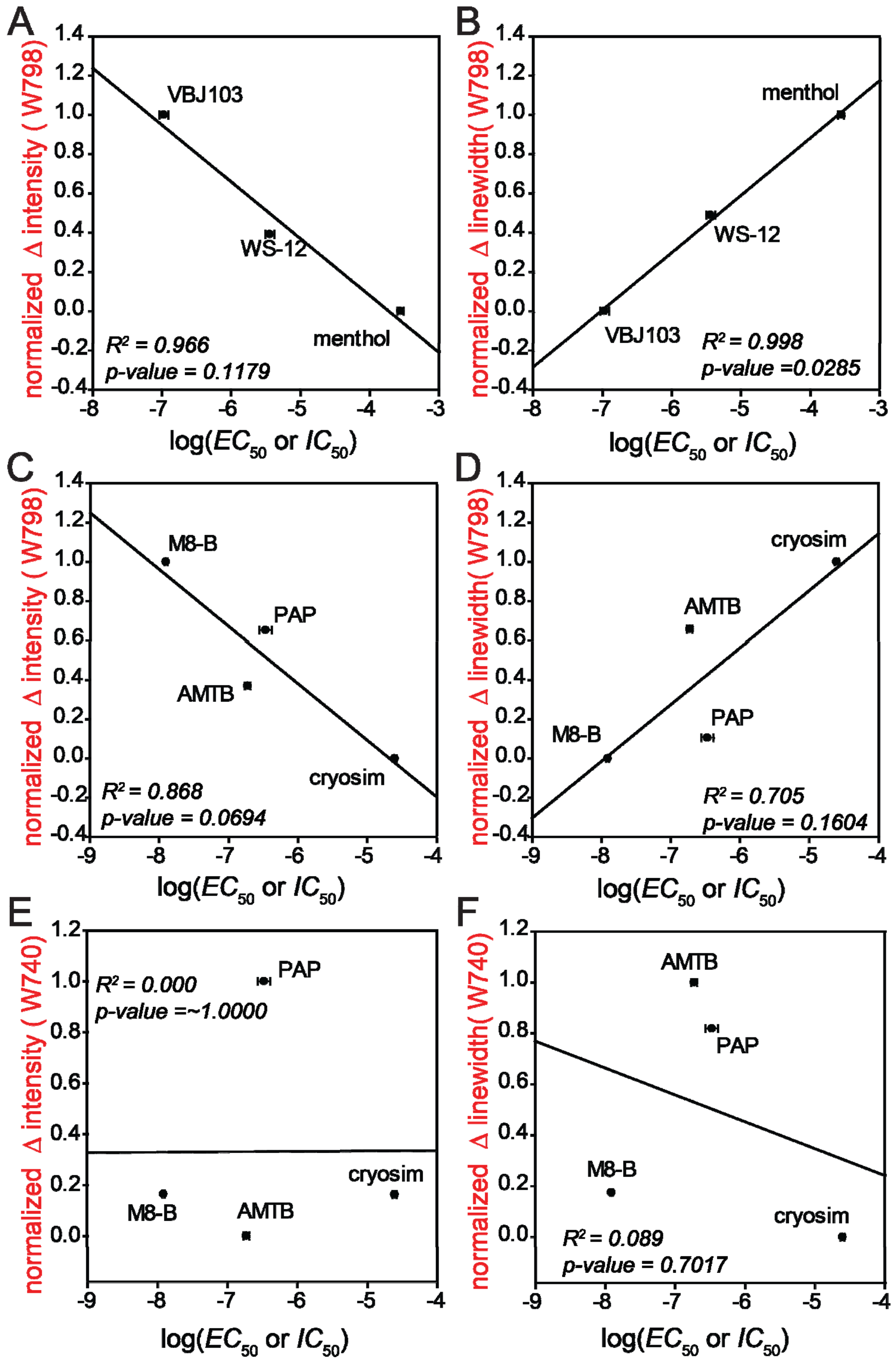

**Figure S19. W798 $^{19}$F NMR peak intensity and linewidth correlate with cellular function**. Correlation between W798 peak intensity and linewidth from round 1 (A and B) and round 3 (C and D) show strong correlation with APC $EC_{50}$ and $IC_{50}$. However, no correlation is observed for resonance features of W740 (E and F) or other resonances (not shown).

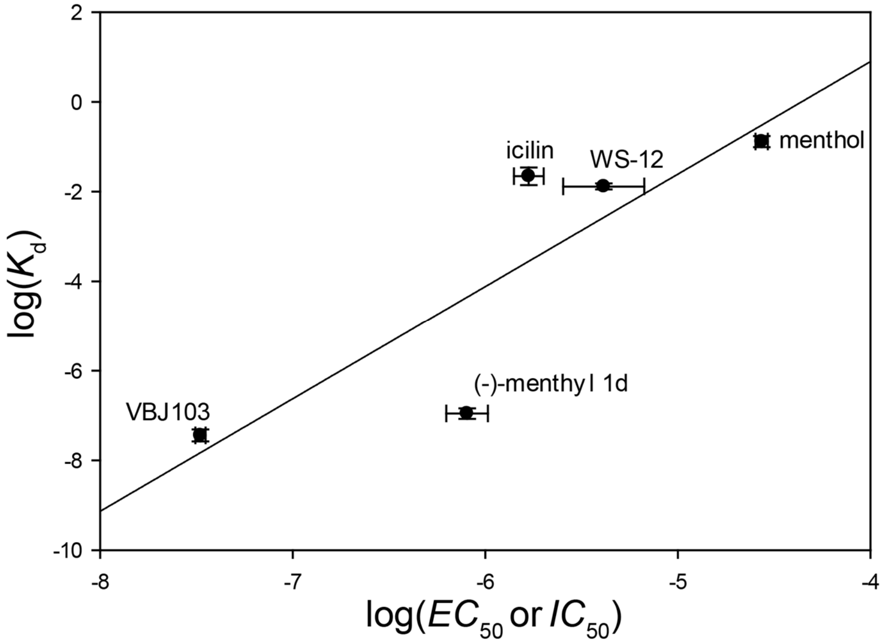

**Figure S20: Correlation between functional activity and binding affinity used to estimate $K_d$ values.** A correlation plot illustrates the relationship between functional activity ($EC_{50}$ and $IC_{50}$) and binding affinity ($K_d$) for several TRPM8 targeting agonist and antagonist compounds. The plot includes values from the literature and includes data generated as part of this study for $EC_{50}$ for menthol[8–10], icilin[9], and WS-12[9,11] as well as $IC_{50}$ values for VBJ103[12] and (-)-menthyl 1d[13] . A linear model was fitted with a $R^2$ 0.722 and p-value 0.0683. This correlation plot was used to estimate $K_d$ values using known $EC_{50}$ for menthyl lactate[14] and cryosim-3[3,15] and $IC_{50}$ values for BCTC[16–20], M8-B[21,22], AMTB[23–25], and PAP[26].

**Table S1. Cryo-EM structures of TRPM8 orthologs**

| Species | Resolution (Å) | Orthosteric Ligand | Other Ligands | PDB ID | Year, Reference |
|---|---|---|---|---|---|
| Human | 2.65 | n/a | $Na^+$, CHS, 9PE, and other undecane molecules | 8BDC | 2023[1] |
| *M. musculus* (mouse) | 3.30 | n/a | CHS | 9B6D | [a]2024[2] |
| | 2.91 | n/a | TC-I 2014, CHS | 9B6E | |
| | 3.42 | n/a | AMG2850, CHS | 9B6F | |
| | 2.81 | AMTB | CHS | 9B6G | |
| | 2.76 | Cryosim-3 | TC-I 2014, CHS | 9B6H | |
| | 3.53 | n/a | PI(4,5)$P_2$, $Ca^{2+}$ | 9B6J | |
| | 4.13 | n/a | $Ca^{2+}$ | 9B6K | |
| *P. major* (Great tit) | 3.26 | n/a | TC-I 2014, CHS, $Ca^{2+}$ | [b]9B6I | |
| *F. albicollis* (Avian, flycatcher) | 3.51 | n/a | PI(4,5)$P_2$ | [b]8E4Q | 2022[3] |
| *M. musculus* (mouse) | 3.59 | n/a | n/a | 8E4P | |
| | 3.43 | n/a | PI(4,5)$P_2$ | 8E4O | |
| | 3.07 | n/a | PI(4,5)$P_2$ | 8E4N | |
| | 3.44 | Cryosim-3 | PI(4,5)$P_2$ | 8E4M | |
| | 3.32 | Cryosim-3 | PI(4,5)$P_2$, AITC | 8E4L | |
| *M. musculus* (mouse) | 2.98 | n/a | n/a | 7WRA | 2022[4] |
| | 2.88 | n/a | $Ca^{2+}$ | 7WRB | |
| | 2.98 | Icilin | $Ca^{2+}$ | 7WRD | |
| | 3.21 | Icilin | PI(4,5)$P_2$, $Ca^{2+}$ | 7WRC | |
| | 2.52 | Icilin | $Ca^{2+}$ | 7WRE | |
| | 3.04 | Icilin | PI(4,5)$P_2$, $Ca^{2+}$ | 7WRF | |
| [b]*P. major* (Great tit) | 3.6 | n/a | n/a | 6O6A | 2019[5] |
| | 3.2 | AMTB | $Na^+$, CHS, 9PE, LQ7, and other undecane molecules | 6O6R | |
| | 3.0 | TC-I 2014 | n/a | 6O72 | |
| | 3.2 | n/a | $Ca^{2+}$ | 6O77 | |
| *F. albicollis* (collared flycatcher) | 3.4 | Icilin | PI(4,5)$P_2$, $Ca^{2+}$ | [c]6NR3 | 2019[6] |
| | 4.3 | Icilin | PI(4,5)$P_2$, $Ca^{2+}$ | [c]6NR4 | |
| | 4.0 | WS-12 | PI(4,5)$P_2$ | [b]6NR2 | |
| [b]*F. albicollis* (collared flycatcher) | 4.1 | n/a | n/a | 6BPQ | 2017[7] |

*[a]TC-2014 binding site discrepancy reported between pdb 9B6I and 6O72 .[b]Avian species, natively icilin insensitive, [c]an Ala805Gly mutation was incorporated to endow icilin sensitivity.*

**Table S2. Pairwise RMSD analysis of TRPM8-VSLD (S1-S4; corresponding to hTRPM8 S733-L853)**

| [a]PDB | 8BDC | 6NR2 | 6NR3 | 6NR4 | 6O6R | 6O72 | 7WRC | 7WRD | 7WRE | 7WRF | 8E4L | 8E4M | 9B6E | 9B6F | 9B6G | 9B6H | 9B6I |
|---|---|---|---|---|---|---|---|---|---|---|---|---|---|---|---|---|---|
| **8BDC** | 0.00 | 0.87 | 1.16 | 0.77 | 0.64 | 0.51 | 0.75 | 0.71 | 0.76 | 0.70 | 1.08 | 0.82 | 0.62 | 0.87 | 0.84 | 0.71 | 0.90 |
| **6NR2** | 0.87 | 0.00 | 0.73 | 0.51 | 0.90 | 0.80 | 0.88 | 0.92 | 0.92 | 0.90 | 1.11 | 0.92 | 0.95 | 0.84 | 0.98 | 0.91 | 1.10 |
| **6NR3** | 1.16 | 0.73 | 0.00 | 0.75 | 1.06 | 0.99 | 0.72 | 0.83 | 0.85 | 0.85 | 0.85 | 0.89 | 1.03 | 0.91 | 1.13 | 1.04 | 0.99 |
| **6NR4** | 0.77 | 0.51 | 0.75 | 0.00 | 0.88 | 0.79 | 0.93 | 0.97 | 0.94 | 0.91 | 1.21 | 0.84 | 0.94 | 0.92 | 1.12 | 0.95 | 1.03 |
| **6O6R** | 0.64 | 0.90 | 1.06 | 0.88 | 0.00 | 0.48 | 0.97 | 0.99 | 0.99 | 0.95 | 1.20 | 1.14 | 0.98 | 1.05 | 1.07 | 0.93 | 0.95 |
| **6O72** | 0.51 | 0.80 | 0.99 | 0.79 | 0.48 | 0.00 | 0.75 | 0.79 | 0.80 | 0.74 | 1.02 | 0.90 | 0.83 | 0.96 | 0.96 | 0.84 | 0.82 |
| **7WRC** | 0.75 | 0.88 | 0.72 | 0.93 | 0.97 | 0.75 | 0.00 | 0.19 | 0.31 | 0.27 | 0.66 | 0.77 | 0.49 | 0.58 | 0.63 | 0.51 | 0.59 |
| **7WRD** | 0.71 | 0.92 | 0.83 | 0.97 | 0.99 | 0.79 | 0.19 | 0.00 | 0.28 | 0.21 | 0.70 | 0.76 | 0.47 | 0.60 | 0.58 | 0.49 | 0.60 |
| **7WRE** | 0.76 | 0.92 | 0.85 | 0.94 | 0.99 | 0.80 | 0.31 | 0.28 | 0.00 | 0.17 | 0.76 | 0.87 | 0.52 | 0.65 | 0.55 | 0.51 | 0.55 |
| **7WRF** | 0.70 | 0.90 | 0.85 | 0.91 | 0.95 | 0.74 | 0.27 | 0.21 | 0.17 | 0.00 | 0.78 | 0.83 | 0.49 | 0.65 | 0.51 | 0.49 | 0.53 |
| **8E4L** | 1.08 | 1.11 | 0.85 | 1.21 | 1.20 | 1.02 | 0.66 | 0.70 | 0.76 | 0.78 | 0.00 | 0.96 | 0.89 | 0.93 | 0.98 | 0.92 | 0.86 |
| **8E4M** | 0.82 | 0.92 | 0.89 | 0.84 | 1.14 | 0.90 | 0.77 | 0.76 | 0.87 | 0.83 | 0.96 | 0.00 | 1.06 | 0.97 | 1.18 | 1.04 | 1.17 |
| **9B6E** | 0.62 | 0.95 | 1.03 | 0.94 | 0.98 | 0.83 | 0.49 | 0.47 | 0.52 | 0.49 | 0.89 | 1.06 | 0.00 | 0.49 | 0.41 | 0.30 | 0.60 |
| **9B6F** | 0.87 | 0.84 | 0.91 | 0.92 | 1.05 | 0.96 | 0.58 | 0.60 | 0.65 | 0.65 | 0.93 | 0.97 | 0.49 | 0.00 | 0.63 | 0.52 | 0.84 |
| **9B6G** | 0.84 | 0.98 | 1.13 | 1.12 | 1.07 | 0.96 | 0.63 | 0.58 | 0.55 | 0.51 | 0.98 | 1.18 | 0.41 | 0.63 | 0.00 | 0.29 | 0.57 |
| **9B6H** | 0.71 | 0.91 | 1.04 | 0.95 | 0.93 | 0.84 | 0.51 | 0.49 | 0.51 | 0.49 | 0.92 | 1.04 | 0.30 | 0.52 | 0.29 | 0.00 | 0.43 |
| **9B6I** | 0.90 | 1.10 | 0.99 | 1.03 | 0.95 | 0.82 | 0.59 | 0.60 | 0.55 | 0.53 | 0.86 | 1.17 | 0.60 | 0.84 | 0.57 | 0.43 | 0.00 |

*[a]All listed structures have ligand bound in the orthosteric site except 8BDC (apo human TRPM8), which is used as a reference and 9B6E, 9B6F, and 9B6I, which have antagonists bound allosterically.*

**Table S3. Pairwise RMSD analysis of all TRPM8 channel structures**

| | 8BDC | 6BPQ | 6NR2 | 6NR3 | 6NR4 | 6O6A | 6O6R | 6O72 | 6O77 | 7WRA | 7WRB | 7WRC | 7WRD | 7WRE | 7WRF | 8E4L | 8E4M | 8E4N | 8E4O | 8E4P | 8E4Q | 9B6D | 9B6E | 9B6F | 9B6G | 9B6H | 9B6I | 9B6J | 9B6K |
|---|---|---|---|---|---|---|---|---|---|---|---|---|---|---|---|---|---|---|---|---|---|---|---|---|---|---|---|---|---|
| **8BDC** | 0 | 2.714 | 3.372 | 3.224 | 2.871 | 1.756 | 1.633 | 1.845 | 4.410 | 3.065 | 3.093 | 3.102 | 3.013 | 3.053 | 3.041 | 3.317 | 3.712 | 3.161 | 3.254 | 2.301 | 2.835 | 4.241 | 3.392 | 4.247 | 3.454 | 3.613 | 3.379 | 2.755 | 4.322 |
| **6BPQ** | 2.714 | 0 | 3.135 | 3.397 | 3.508 | 3.710 | 3.290 | 3.007 | 4.272 | 4.256 | 4.246 | 4.678 | 4.248 | 3.390 | 4.692 | 3.774 | 3.721 | 4.046 | 3.848 | 2.833 | 2.884 | 4.427 | 4.172 | 4.956 | 3.903 | 3.535 | 3.540 | 3.252 | 4.508 |
| **6NR2** | 3.372 | 3.135 | 0 | 2.782 | 1.948 | 3.531 | 3.975 | 3.474 | 4.179 | 4.029 | 3.470 | 3.842 | 3.763 | 3.929 | 3.943 | 3.787 | 4.132 | 3.644 | 3.642 | 1.924 | 2.456 | 3.699 | 3.731 | 3.489 | 3.736 | 4.067 | 4.424 | 3.108 | 3.981 |
| **6NR3** | 3.224 | 3.397 | 2.782 | 0 | 2.826 | 3.900 | 3.957 | 3.863 | 4.219 | 2.859 | 2.870 | 2.930 | 2.975 | 2.730 | 3.067 | 3.727 | 2.919 | 2.448 | 2.432 | 2.868 | 2.861 | 3.100 | 2.847 | 2.960 | 2.856 | 4.437 | 3.422 | 2.562 | 3.162 |
| **6NR4** | 2.871 | 3.508 | 1.948 | 2.826 | 0 | 3.214 | 3.945 | 3.412 | 4.116 | 4.096 | 4.116 | 3.595 | 4.586 | 3.793 | 3.387 | 2.701 | 3.412 | 3.647 | 3.350 | 2.471 | 2.512 | 3.717 | 3.124 | 3.552 | 3.763 | 3.660 | 4.274 | 3.043 | 3.325 |
| **6O6A** | 1.756 | 3.710 | 3.531 | 3.900 | 3.214 | 0 | 1.594 | 2.121 | 4.031 | 3.371 | 3.387 | 3.378 | 3.373 | 3.408 | 3.396 | 3.525 | 3.795 | 3.572 | 3.991 | 3.327 | 3.292 | 4.186 | 3.435 | 4.461 | 4.120 | 4.361 | 3.842 | 2.754 | 4.214 |
| **6O6R** | 1.633 | 3.290 | 3.975 | 3.957 | 3.945 | 1.594 | 0 | 1.903 | 3.939 | 3.423 | 3.583 | 3.448 | 3.358 | 3.457 | 3.361 | 3.324 | 3.666 | 4.136 | 3.701 | 2.516 | 3.186 | 4.659 | 4.571 | 4.464 | 4.445 | 4.365 | 3.934 | 3.074 | 4.206 |
| **6O72** | 1.845 | 3.007 | 3.474 | 3.863 | 3.412 | 2.121 | 1.903 | 0 | 3.787 | 3.723 | 3.765 | 3.566 | 3.399 | 3.378 | 3.566 | 3.859 | 3.586 | 3.515 | 3.759 | 3.116 | 3.100 | 4.493 | 3.534 | 4.387 | 3.541 | 4.154 | 4.084 | 3.059 | 3.957 |
| **6O77** | 4.410 | 4.272 | 4.179 | 4.219 | 4.116 | 4.031 | 3.939 | 3.787 | 0 | 3.133 | 3.100 | 3.103 | 3.683 | 3.221 | 3.409 | 3.250 | 3.079 | 2.966 | 3.144 | 3.815 | 4.225 | 4.036 | 3.765 | 3.884 | 3.410 | 3.106 | 3.464 | 3.733 | 3.686 |
| **7WRA** | 3.065 | 4.256 | 4.029 | 2.859 | 4.096 | 3.371 | 3.423 | 3.723 | 3.133 | 0 | 0.323 | 1.001 | 0.596 | 0.692 | 0.665 | 3.053 | 2.755 | 2.231 | 1.948 | 2.707 | 2.950 | 2.747 | 1.831 | 2.880 | 2.093 | 2.158 | 3.463 | 2.358 | 2.425 |
| **7WRB** | 3.093 | 4.246 | 3.470 | 2.870 | 4.116 | 3.387 | 3.583 | 3.765 | 3.100 | 0.323 | 0 | 0.929 | 0.539 | 0.613 | 0.594 | 2.301 | 2.966 | 2.247 | 2.530 | 2.728 | 3.002 | 2.904 | 1.953 | 2.665 | 2.109 | 2.153 | 3.176 | 2.356 | 2.425 |
| **7WRC** | 3.102 | 4.678 | 3.842 | 2.930 | 3.595 | 3.378 | 3.448 | 3.566 | 3.103 | 1.001 | 0.929 | 0 | 0.582 | 0.594 | 0.582 | 1.968 | 2.783 | 1.979 | 1.826 | 2.587 | 2.878 | 3.118 | 1.951 | 3.109 | 2.244 | 2.307 | 2.724 | 2.245 | 2.652 |
| **7WRD** | 3.013 | 4.248 | 3.763 | 2.975 | 4.586 | 3.373 | 3.358 | 3.399 | 3.683 | 0.596 | 0.539 | 0.582 | 0 | 0.328 | 0.292 | 2.146 | 3.217 | 2.180 | 1.869 | 2.709 | 2.985 | 3.214 | 1.980 | 2.780 | 2.201 | 2.270 | 3.534 | 2.298 | 2.562 |
| **7WRE** | 3.053 | 3.390 | 3.929 | 2.730 | 3.793 | 3.408 | 3.457 | 3.378 | 3.221 | 0.692 | 0.613 | 0.594 | 0.328 | 0 | 0.261 | 2.236 | 3.442 | 2.230 | 1.931 | 2.706 | 2.940 | 2.798 | 2.073 | 2.732 | 2.221 | 2.280 | 2.471 | 2.375 | 2.575 |
| **7WRF** | 3.041 | 4.692 | 3.943 | 3.067 | 3.387 | 3.396 | 3.361 | 3.566 | 3.409 | 0.665 | 0.594 | 0.582 | 0.292 | 0.261 | 0 | 2.261 | 2.924 | 2.176 | 1.902 | 2.691 | 2.935 | 2.837 | 2.052 | 2.694 | 2.227 | 2.293 | 3.267 | 2.396 | 2.579 |
| **8E4L** | 3.317 | 3.774 | 3.787 | 3.727 | 2.701 | 3.525 | 3.324 | 3.859 | 3.250 | 3.053 | 2.301 | 1.968 | 2.146 | 2.236 | 2.261 | 0 | 2.565 | 2.358 | 2.400 | 3.178 | 3.814 | 3.317 | 3.120 | 3.500 | 4.191 | 3.535 | 3.134 | 2.501 | 3.649 |
| **8E4M** | 3.712 | 3.721 | 4.132 | 2.919 | 3.412 | 3.795 | 3.666 | 3.586 | 3.079 | 2.755 | 2.966 | 2.783 | 3.217 | 3.442 | 2.924 | 2.565 | 0 | 1.674 | 1.814 | 3.453 | 4.184 | 4.474 | 2.917 | 3.072 | 2.865 | 2.900 | 3.909 | 1.717 | 3.221 |
| **8E4N** | 3.161 | 4.046 | 3.644 | 2.448 | 3.647 | 3.572 | 4.136 | 3.515 | 2.966 | 2.231 | 2.247 | 1.979 | 2.180 | 2.230 | 2.176 | 2.358 | 1.674 | 0 | 0.858 | 3.531 | 3.931 | 3.663 | 3.047 | 3.817 | 3.280 | 3.116 | 3.351 | 2.233 | 3.514 |
| **8E4O** | 3.254 | 3.848 | 3.642 | 2.432 | 3.350 | 3.991 | 3.701 | 3.759 | 3.144 | 1.948 | 2.530 | 1.826 | 1.869 | 1.931 | 1.902 | 2.400 | 1.814 | 0.858 | 0 | 3.569 | 3.572 | 3.346 | 2.951 | 3.057 | 2.607 | 3.417 | 3.190 | 1.994 | 3.289 |
| **8E4P** | 2.301 | 2.833 | 1.924 | 2.868 | 2.471 | 3.327 | 2.516 | 3.116 | 3.815 | 2.707 | 2.728 | 2.587 | 2.709 | 2.706 | 2.691 | 3.178 | 3.453 | 3.531 | 3.569 | 0 | 2.628 | 3.485 | 3.287 | 3.336 | 3.413 | 3.547 | 3.871 | 2.557 | 3.458 |
| **8E4Q** | 2.835 | 2.884 | 2.456 | 2.861 | 2.512 | 3.292 | 3.186 | 3.100 | 4.225 | 2.950 | 3.002 | 2.878 | 2.985 | 2.940 | 2.935 | 3.814 | 4.184 | 3.931 | 3.572 | 2.628 | 0 | 2.835 | 3.883 | 3.699 | 3.764 | 3.832 | 4.399 | 4.149 | 4.383 |
| **9B6D** | 4.241 | 4.427 | 3.699 | 3.100 | 3.717 | 4.186 | 4.659 | 4.493 | 4.036 | 2.747 | 2.904 | 3.118 | 3.214 | 2.798 | 2.837 | 3.317 | 4.474 | 3.663 | 3.346 | 3.485 | 2.835 | 0 | 1.867 | 1.325 | 2.240 | 2.285 | 2.825 | 3.598 | 2.510 |
| **9B6E** | 3.392 | 4.172 | 3.731 | 2.847 | 3.124 | 3.435 | 4.571 | 3.534 | 3.765 | 1.831 | 1.953 | 1.951 | 1.980 | 2.073 | 2.052 | 3.120 | 2.917 | 3.047 | 2.951 | 3.287 | 3.883 | 1.867 | 0 | 1.325 | 1.031 | 1.101 | 3.153 | 3.099 | 2.005 |
| **9B6F** | 4.247 | 4.956 | 3.489 | 2.960 | 3.552 | 4.461 | 4.464 | 4.387 | 3.884 | 2.880 | 2.665 | 3.109 | 2.780 | 2.732 | 2.694 | 3.500 | 3.072 | 3.817 | 3.057 | 3.336 | 3.699 | 1.325 | 1.325 | 0 | 2.532 | 2.679 | 3.309 | 3.502 | 2.556 |
| **9B6G** | 3.454 | 3.903 | 3.736 | 2.856 | 3.763 | 4.120 | 4.445 | 3.541 | 3.410 | 2.093 | 2.109 | 2.244 | 2.201 | 2.221 | 2.227 | 4.191 | 2.865 | 3.280 | 2.607 | 3.413 | 3.764 | 2.240 | 1.031 | 2.532 | 0 | 1.219 | 3.309 | 3.116 | 2.432 |
| **9B6H** | 3.613 | 3.535 | 4.067 | 4.437 | 3.660 | 4.361 | 4.365 | 4.154 | 3.106 | 2.158 | 2.153 | 2.307 | 2.270 | 2.280 | 2.293 | 3.535 | 2.900 | 3.116 | 3.417 | 3.547 | 3.832 | 2.285 | 1.101 | 2.679 | 1.219 | 0 | 2.396 | 3.523 | 2.177 |
| **9B6I** | 3.379 | 3.540 | 4.424 | 3.422 | 4.274 | 3.842 | 3.934 | 4.084 | 3.464 | 3.463 | 3.176 | 2.724 | 3.534 | 2.471 | 3.267 | 3.134 | 3.909 | 3.351 | 3.190 | 3.871 | 4.399 | 2.825 | 3.153 | 3.309 | 3.309 | 2.396 | 0 | 3.440 | 2.743 |
| **9B6J** | 2.755 | 3.252 | 3.108 | 2.562 | 3.043 | 2.754 | 3.074 | 3.059 | 3.733 | 2.358 | 2.356 | 2.245 | 2.298 | 2.375 | 2.396 | 2.501 | 1.717 | 2.233 | 1.994 | 2.557 | 4.149 | 3.598 | 3.099 | 3.502 | 3.116 | 3.523 | 3.440 | 0 | 3.401 |
| **9B6K** | 4.322 | 4.508 | 3.981 | 3.162 | 3.325 | 4.214 | 4.206 | 3.957 | 3.686 | 2.425 | 2.425 | 2.652 | 2.562 | 2.575 | 2.579 | 3.649 | 3.221 | 3.514 | 3.289 | 3.458 | 4.383 | 2.510 | 2.005 | 2.556 | 2.432 | 2.177 | 2.743 | 3.401 | 0 |

**Table S4. Expression levels of TRPM8 S1–S4 domain under various conditions.**[a]

| | [IPTG] (mM) | Cell Line | | | | | | |
|---|---|---|---|---|---|---|---|---|
| | | 0 | 1 | 2 | 3 | 4 | 6 | 8 |
| 18 °C | 0.3 | 0.21 | 0.06 | 0.26 | 0.39 | 0.24 | 0.42 | 0.52 |
| | 0.5 | 0.24 | 0.06 | 0.28 | 0.49 | 0.43 | 0.59 | 0.72 |
| | 1 | 0.26 | 0.06 | 0.41 | 0.64 | 0.66 | 0.79 | 0.79 |
| 25 °C | 0.3 | 0.34 | 0.21 | 0.56 | 0.77 | 0.72 | 1 | 0.86 |
| | 0.5 | 0.3 | 0.14 | 0.36 | 0.49 | 0.47 | 0.52 | 0.53 |
| | 1 | 0.35 | 0.13 | 0.18 | 0.36 | 0.34 | 0.42 | 0.41 |
| 37 °C | 0.3 | 0.37 | 0.29 | 0.24 | 0.38 | 0.24 | 0.51 | 0.47 |
| | 0.5 | 0.58 | 0.53 | 0.28 | 0.45 | 0.35 | 0.57 | 0.5 |
| | 1 | 0.79 | 0.79 | 0.24 | 0.54 | 0.62 | 0.75 | 0.51 |

[a]*Signal intensity was quantified using ImageJ. The numbers were normalized to the highest value, and the color gradient from purple to red to yellow to represent the lowest to highest signal intensity. Cell lines are numbered as in Figure S5.*

**Table S5. Menthol or Menthol+VBJ103 induced chemical shift change (Δδ)**

| Residue | Menthol (\|Δδ\| ppm) relative to apo | Menthol+VBJ103 (\|Δδ\| ppm) relative to apo |
|---|---|---|
| W725 | 0.045 | 0.013 |
| W740 | 0.073 | 0.036 |
| W786 | 0.049 | 0.046 |
| W798 | 0.039 | 0.022 |

**Table S6. $^{1}H$ lineshape analysis of $^{1}H$-$^{15}N$ HSQC based agonist and antagonist titration**

| | $^{1}H$ | | | | | | | |
|---|---|---|---|---|---|---|---|---|
| | Menthol | | Icilin | | VBJ103 | | Menthyl 1d | |
| Resonance | Δλ (Hz) | Std Error | Δλ (Hz) | Std Error | Δλ (Hz) | Std Error | Δλ (Hz) | Std Error |
| a | 0 | 35 | -454 | 59 | -50 | 28 | 142 | 393 |
| b | 2 | 26 | -232 | 44 | -306 | 39 | 203 | 421 |
| d | -4 | 4 | -59 | 10 | 156 | 3 | 223 | 370 |
| e | 2 | 11 | -94 | 20 | -128 | 9 | 177 | 398 |
| f | 4 | 20 | -152 | 25 | 747 | 20 | 153 | 406 |
| h | 2 | 11 | -33 | 16 | 834 | 8 | 213 | 379 |
| c | 4 | 20 | -142 | 25 | 38 | 9 | 154 | 386 |
| g | 2 | 41 | -80 | 23 | -40 | 23 | 79 | 355 |
| i | 5 | 12 | -308 | 42 | 135 | 12 | 181 | 380 |
| $\Delta\lambda_{Avg}$ | 2 | 8 | -172 | 11 | 154 | 7 | 169 | 129 |

**Table S7. $^{15}N$ lineshape analysis of $^{1}H$-$^{15}N$ HSQC based agonist and antagonist titration**

| | $^{15}N$ | | | | | | | |
|---|---|---|---|---|---|---|---|---|
| | Menthol | | Icilin | | VBJ103 | | Menthyl 1d | |
| Resonance | Δλ (Hz) | Std Error | Δλ (Hz) | Std Error | Δλ (Hz) | Std Error | Δλ (Hz) | Std Error |
| a | -3 | 22 | -201 | 28 | -141 | 12 | 121 | 341 |
| b | 16 | 19 | -108 | 32 | 24 | 10 | 122 | 352 |
| d | 5 | 5 | -68 | 17 | 64 | 4 | 64 | 379 |
| e | 0 | 9 | -34 | 18 | 46 | 5 | 166 | 345 |
| f | -3 | 17 | -70 | 26 | 338 | 9 | 25 | 310 |
| h | 1 | 9 | -6 | 20 | 166 | 6 | 182 | 391 |
| c | -1 | 25 | -88 | 29 | 16 | 7 | 51 | 288 |
| g | 5 | 33 | -34 | 24 | 4 | 16 | 131 | 370 |
| i | 0 | 11 | -55 | 13 | -53 | 5 | 129 | 367 |
| $\Delta\lambda_{Avg}$ | 2 | 6 | -74 | 8 | 52 | 3 | 110 | 117 |

### REFERENCES

(1) Palchevskyi, S.; Czarnocki-Cieciura, M.; Vistoli, G.; Gervasoni, S.; Nowak, E.; Beccari, A. R.; Nowotny, M.; Talarico, C. Structure of Human TRPM8 Channel. *Commun Biol* **2023**, *6* (1), 1065. https://doi.org/10.1038/s42003-023-05425-6.

(2) Yin, Y.; Park, C.-G.; Zhang, F.; Fedor, J. G.; Feng, S.; Suo, Y.; Im, W.; Lee, S.-Y. *Mechanisms of Sensory Adaptation and Inhibition of the Cold and Menthol Receptor TRPM8*; 2024; Vol. 10. https://www.science.org.

(3) Yin, Y.; Zhang, F.; Feng, S.; Butay, K. J.; Borgnia, M. J.; Im, W.; Lee, S.-Y. Activation Mechanism of the Mouse Cold-Sensing TRPM8 Channel by Cooling Agonist and PIP $_2$. *Science (1979)* **2022**, *378* (6616), eadd1268. https://doi.org/10.1126/science.add1268.

(4) Zhao, C.; Xie, Y.; Xu, L.; Ye, F.; Xu, X.; Yang, W.; Yang, F.; Guo, J. Structures of a Mammalian TRPM8 in Closed State. *Nat Commun* **2022**, *13* (1), 3113. https://doi.org/10.1038/s41467-022-30919-y.

(5) Diver, M. M.; Cheng, Y.; Julius, D. Structural Insights into TRPM8 Inhibition and Desensitization. *Science (1979)* **2019**, *365* (6460), 1434–1440. https://doi.org/10.1126/science.aax6672.

(6) Yin, Y.; Le, S. C.; Hsu, A. L.; Borgnia, M. J.; Yang, H.; Lee, S.-Y. Structural Basis of Cooling Agent and Lipid Sensing by the Cold-Activated TRPM8 Channel. *Science (1979)* **2019**, *363* (6430), eaav9334. https://doi.org/10.1126/science.aav9334.

(7) Yin, Y.; Wu, M.; Zubcevic, L.; Borschel, W. F.; Lander, G. C.; Lee, S.-Y. Structure of the Cold- and Menthol-Sensing Ion Channel TRPM8. *Science (1979)* **2018**, *359* (6372), 237–241. https://doi.org/10.1126/science.aan4325.

(8) Voets, T.; Owsianik, G.; Janssens, A.; Talavera, K.; Nilius, B. TRPM8 Voltage Sensor Mutants Reveal a Mechanism for Integrating Thermal and Chemical Stimuli. *Nat Chem Biol* **2007**, *3* (3), 174–182. https://doi.org/10.1038/nchembio862.

(9) Bödding, M.; Wissenbach, U.; Flockerzi, V. Characterisation of TRPM8 as a Pharmacophore Receptor. *Cell Calcium* **2007**, *42* (6), 618–628. https://doi.org/10.1016/j.ceca.2007.03.005.

(10) Luu, D. D.; Ramesh, N.; Kazan, I. C.; Shah, K. H.; Lahiri, G.; Mana, M. D.; Ozkan, S. B.; Van Horn, W. D. Evidence That the Cold- and Menthol-Sensing Functions of the Human TRPM8 Channel Evolved Separately. *Sci Adv* **2024**, *10* (25), 9228. https://doi.org/10.1126/sciadv.adm9228.

(11) Sherkheli, M. A.; Vogt-Eisele, A. K.; Bura, D.; Beltrán Márques, L. R.; Gisselmann, G.; Hatt, H. Characterization Of Selective TRPM8 Ligands And Their Structure Activity Response (S.A.R) Relationship. *Journal of Pharmacy & Pharmaceutical Sciences* **2010**, *13* (2), 242. https://doi.org/10.18433/J3N88N.

(12) Journigan, V. B.; Feng, Z.; Rahman, S.; Wang, Y.; Amin, A. R. M. R.; Heffner, C. E.; Bachtel, N.; Wang, S.; Gonzalez-Rodriguez, S.; Fernández-Carvajal, A.; Fernández-Ballester, G.; Hilton, J. K.; Van Horn, W. D.; Ferrer-Montiel, A.; Xie, X.-Q.; Rahman, T. Structure-Based Design of Novel Biphenyl Amide Antagonists of Human Transient Receptor Potential Cation Channel Subfamily M Member 8 Channels with Potential Implications in the Treatment of Sensory Neuropathies. *ACS Chem Neurosci* **2020**, *11* (3), 268–290. https://doi.org/10.1021/acschemneuro.9b00404.

(13) Journigan, V. B.; Alarcón-Alarcón, D.; Feng, Z.; Wang, Y.; Liang, T.; Dawley, D. C.; Amin, A. R. M. R.; Montano, C.; Van Horn, W. D.; Xie, X. Q.; Ferrer-Montiel, A.; Fernández-Carvajal, A. Structural and in Vitro Functional Characterization of a Menthyl TRPM8 Antagonist Indicates Species-Dependent Regulation. *ACS Med Chem Lett* **2021**, *12* (5), 758–767. https://doi.org/10.1021/acsmedchemlett.1c00001.

(14) Bharate, S. S.; Bharate, S. B. Modulation of Thermoreceptor TRPM8 by Cooling Compounds. *ACS Chem Neurosci* **2012**, *3* (4), 248–267. https://doi.org/10.1021/cn300006u.

(15) Yang, J. M.; Li, F.; Liu, Q.; Rüedi, M.; Wei, E. T.; Lentsman, M.; Lee, H. S.; Choi, W.; Kim, S. J.; Yoon, K. C. A Novel TRPM8 Agonist Relieves Dry Eye Discomfort. *BMC Ophthalmol* **2017**, *17* (1), 101. https://doi.org/10.1186/s12886-017-0495-2.

(16) Liu, T.; Fang, Z.; Wang, G.; Shi, M.; Wang, X.; Jiang, K.; Yang, Z.; Cao, R.; Tao, H.; Wang, X.; Zhou, J. Anti-Tumor Activity of the TRPM8 Inhibitor BCTC in Prostate Cancer DU145 Cells. *Oncol Lett* **2016**, *11* (1), 182–188. https://doi.org/10.3892/ol.2015.3854.

(17) Valenzano, K. J.; Grant, E. R.; Wu, G.; Hachicha, M.; Schmid, L.; Tafesse, L.; Sun, Q.; Rotshteyn, Y.; Francis, J.; Limberis, J.; Malik, S.; Whittemore, E. R.; Hodges, D. N-(4-Tertiarybutylphenyl)-4-(3-Chloropyridin-2-Yl)Tetrahydropyrazine-1(2H)-Carbox-Amide (BCTC), a Novel, Orally Effective Vanilloid Receptor 1 Antagonist with Analgesic Properties: I. In Vitro Characterization and Pharmacokinetic Properties. *Journal of Pharmacology and Experimental Therapeutics* **2003**, *306* (1), 377–386. https://doi.org/10.1124/jpet.102.045674.

(18) Behrendt, H. J.; Germann, T.; Gillen, C.; Hatt, H.; Jostock, R. Characterization of the Mouse Cold-Menthol Receptor TRPM8 and Vanilloid Receptor Type-1 VR1 Using a Fluorometric Imaging Plate Reader (FLIPR) Assay. *Br J Pharmacol* **2004**, *141* (4), 737–745. https://doi.org/10.1038/sj.bjp.0705652.

(19) Liu, Y.; Lubin, M. Lou; Reitz, T. L.; Wang, Y.; Colburn, R. W.; Flores, C. M.; Qin, N. Molecular Identification and Functional Characterization of a Temperature-Sensitive Transient Receptor Potential Channel (TRPM8) from Canine. *Eur J Pharmacol* **2006**, *530* (1–2), 23–32. https://doi.org/10.1016/j.ejphar.2005.11.033.

(20) Weil, A.; Moore, S. E.; Waite, N. J.; Randall, A.; Gunthorpe, M. J. Conservation of Functional and Pharmacological Properties in the Distantly Related Temperature Sensors TRVP1 and TRPM8. *Mol Pharmacol* **2005**, *68* (2), 518–527. https://doi.org/10.1124/mol.105.012146.

(21) Fakih, D.; Baudouin, C.; Goazigo, A. R. Le; Parsadaniantz, S. M. TRPM8: A Therapeutic Target for Neuroinflammatory Symptoms Induced by Severe Dry Eye Disease. *Int J Mol Sci* **2020**, *21* (22), 1–21. https://doi.org/10.3390/ijms21228756.

(22) Miller, S.; Rao, S.; Wang, W.; Liu, H.; Wang, J.; Gavva, N. R. Antibodies to the Extracellular Pore Loop of TRPM8 Act as Antagonists of Channel Activation. *PLoS One* **2014**, *9* (9), e107151. https://doi.org/10.1371/journal.pone.0107151.

(23) Lashinger, E. S. R.; Steiginga, M. S.; Hieble, J. P.; Leon, L. A.; Gardner, S. D.; Nagilla, R.; Davenport, E. A.; Hoffman, B. E.; Laping, N. J.; Su, X. AMTB, a TRPM8 Channel Blocker: Evidence in Rats for Activity in Overactive Bladder and Painful Bladder Syndrome. *American Journal of Physiology-Renal Physiology* **2008**, *295* (3), F803–F810. https://doi.org/10.1152/ajprenal.90269.2008.

(24) Yapa, K. T. D. S.; Deuis, J.; Peters, A. A.; Kenny, P. A.; Roberts-Thomson, S. J.; Vetter, I.; Monteith, G. R. Assessment of the TRPM8 Inhibitor AMTB in Breast Cancer Cells and Its Identification as an Inhibitor of Voltage Gated Sodium Channels. *Life Sci* **2018**, *198*, 128–135. https://doi.org/10.1016/j.lfs.2018.02.030.

(25) Liu, Y.; Leng, A.; Li, L.; Yang, B.; Shen, S.; Chen, H.; Zhu, E.; Xu, Q.; Ma, X.; Shi, P.; Liu, Y.; Liu, T.; Li, L.; Li, K.; Zhang, D.; Xiao, J. AMTB, a TRPM8 Antagonist, Suppresses Growth and Metastasis of Osteosarcoma through Repressing the TGFβ Signaling Pathway. *Cell Death Dis* **2022**, *13* (3), 288. https://doi.org/10.1038/s41419-022-04744-6.

(26) Luyts, N.; Daniluk, J.; Freitas, A. C. N.; Bazeli, B.; Janssens, A.; Mulier, M.; Everaerts, W.; Voets, T. Inhibition of TRPM8 by the Urinary Tract Analgesic Drug Phenazopyridine. *Eur J Pharmacol* **2023**, *942*, 175512. https://doi.org/10.1016/j.ejphar.2023.175512.
